# Thalamic oscillations distinguish natural states of consciousness in humans

**DOI:** 10.1101/2025.01.28.635248

**Authors:** Aditya Chowdhury, Xiongbo Wu, Tara Beilner, Thomas Schreiner, Thomas Koeglsperger, Jan-Hinnerk Mehrkens, Jan Remi, Christian Vollmar, Elisabeth Kaufmann, Tobias Staudigl

## Abstract

Natural states of consciousness are thought to be regulated by deep brain structures such as the thalamus. However, very little is known about the underlying electrophysiology in humans. Here, using a rare opportunity to directly record from the human thalamus, we identify a hitherto-unreported brain-state-specific oscillation of approximately 19-45 Hz. This oscillation is present only during Rapid Eye Movement (REM) sleep and wakefulness, while being absent during Non-REM sleep. The 19-45 oscillation further distinguishes REM sleep microstates, co-occurring with bursts of eye movements, and is specific to the Central Thalamus, a structure implicated in causing global brain state transitions. The discovery of a distinct oscillatory signature in the Central Thalamus that distinguishes conscious states opens up avenues to further investigate thalamic contributions to states of consciousness in humans and potentially to refine interventions to treat disorders of consciousness.

## 1 Introduction

Thalamocortical networks orchestrate neural oscillations over a range of frequencies, serving to selectively synchronize different brain regions to meet the specific requirements of a given brain state [see e.g. refs^1,2^ for reviews]. For example, during Non-Rapid Eye Movement (NREM) sleep, thalamocortical sleep spindles at 11-17 Hz are generated via a mechanism driven by inhibitory neurons in the thalamic reticular nucleus^3,4^; such oscillations are hypothesized to selectively activate various brain networks required for cognitive processes^5,6^. Indeed, sleep spindles form a crucial part of the systems memory consolidation hypothesis that depends on oscillations-mediated communication between the thalamus, the hippocampus, and the cortex^6^. While sleep spindles in the thalamus of model animals have been studied for many decades now, similar research in the human thalamus has been possible only relatively recently. Several researchers have used recordings from patients with bilateral Deep Brain Stimulation (DBS) electrodes^7^ or stereoelectroencephalography from the thalamus to advance our understanding of thalamic contributions to sleep oscillations in humans during NREM sleep^8–11^. However, the relation of thalamic activity to other key natural brain states such as wakefulness and REM sleep has remained severely underexplored [but see refs.^12,13^].

Studies in non-human primates point to a critical role of the Central Thalamus in regulating brain states^14–16^. For example, electrical stimulation of the Central Thalamus can lead to arousal in anesthetized animals^17–19^. In humans, stimulating the Central Thalamus has shown promising but mixed results in reversing the loss of consciousness [see refs.^20,21^ for reviews]. It is currently unclear why DBS of the Central Thalamus leads to a clear improvement in levels of consciousness in certain patients but not in others. One crucial limiting factor here has been the lack of characterization of how thalamic electrophysiology indicates natural states of consciousness in humans, especially in cohorts without disorders of consciousness.

Here we use direct electrophysiology of the thalamus in a cohort of 17 epilepsy patients (age: 35 ± 10 years, 9 males) implanted with DBS electrodes, with continuous recordings typically much longer than one night (median recording duration of ∼ 40 hr, see Methods), to identify a ubiquitous fast rhythm (19-45 Hz) in the thalamus during both wakefulness and REM sleep. We find that the bursts of this state-specific thalamic oscillatory activity are tightly coupled to eye movements during REM sleep, thus further distinguishing between the phasic and the tonic microstates of REM sleep. We further show that the Wake- and REM sleep- specific fast thalamic oscillations have burst amplitudes that correlate with those of sleep spindles detected during NREM sleep, indicative of a common generation mechanism. Finally, we show that the probability of detecting this electrophysiological marker of the state of consciousness in the human thalamus is significantly higher when the electrode contact is in close proximity to the Central Thalamus.

## 2 Results

### 2.1 A fast oscillation in the human thalamus distinguishes wakefulness and REM sleep from NREM sleep

We aimed to understand the spectrotemporal properties of thalamic field potential as a function of natural states of consciousness. The patients in this study were bilaterally implanted with DBS electrodes via a trans-ventricular approach, with four potential thalamic contacts on each hemisphere (Medtronic 3387; see Figure 1(a) for an example electrode trajectory). We obtained the thalamic field potential by bipolarly rereferencing the voltages recorded in neighboring contacts of the DBS electrodes, obtaining a total of 105 channels (see Methods).

**Figure 1:**
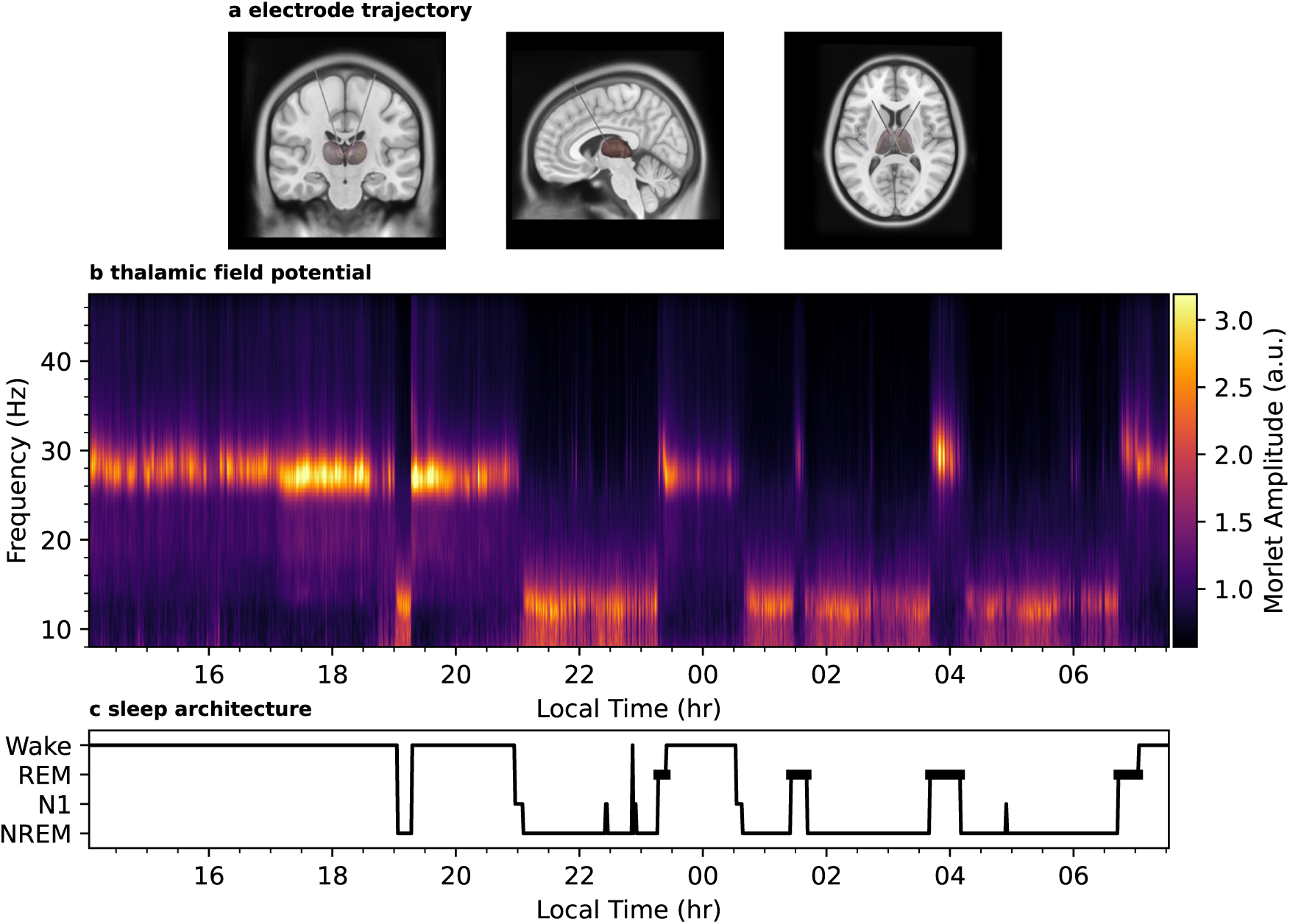
Example electrode trajectory and thalamic field potential recordings over extended time periods. Panel **(a)** illustrates the bilateral implantation scheme used for the DBS electrodes for all patients in our study. The brown region indicates the thalamus. The visualization was generated using LeadDBS^22^. Panel **(b)** shows the thalamic field potential from an example bipolarly referenced chan-nel, as a function of frequency and time, over a continuous period of *≈* 16 hr. Panel **(c)** shows the sleep state classification for the same duration. The consistent presence of sleep spindles at *≈* 11 *−* 15 Hz during NREM sleep and a faster oscillatory activity at *≈* 28 *−* 34 Hz during both wakefulness *and* REM sleep can be clearly seen.

We observed, throughout our cohort, a remarkably consistent pattern of oscillatory activity in different brain states. Figure 1(b) shows the Morlet-transformed thalamic field potential from a single bipolar channel, throughout a continuous period of ≈ 16 hrs that includes a night of sleep. Besides the prominent presence of sleep spindles (≈ 12 Hz) during NREM sleep that has been described in the human thalamus before^11^, the figure shows the ubiquitous presence of a faster oscillation (≈ 28 Hz) during both wakefulness and REM sleep that, to the best of our knowledge, has not been reported in the human thalamus before.

We systematically characterized the oscillatory activity in each of the thalamic contacts by detecting bursts of oscillatory activity across the entire recording epochs. The burst detection was based on the Morlet amplitudes, at a temporal resolution of 5 milliseconds, which were then used to obtain a *burst rate* as a function of time and frequency for the entire duration of the recording (see Methods for a detailed description). Next, we estimated the mean burst rate as a function of frequency for each of the three natural brain states of interest here: (i) wakefulness, (ii) NREM sleep, and (iii) REM sleep. Finally, we estimated, for each of the brain states separately, the frequency ranges with significantly elevated burst probabilities (see Methods). Figure 2(a) illustrates this entire procedure for the same thalamic channel as in Figure 1(b).

**Figure 2:**
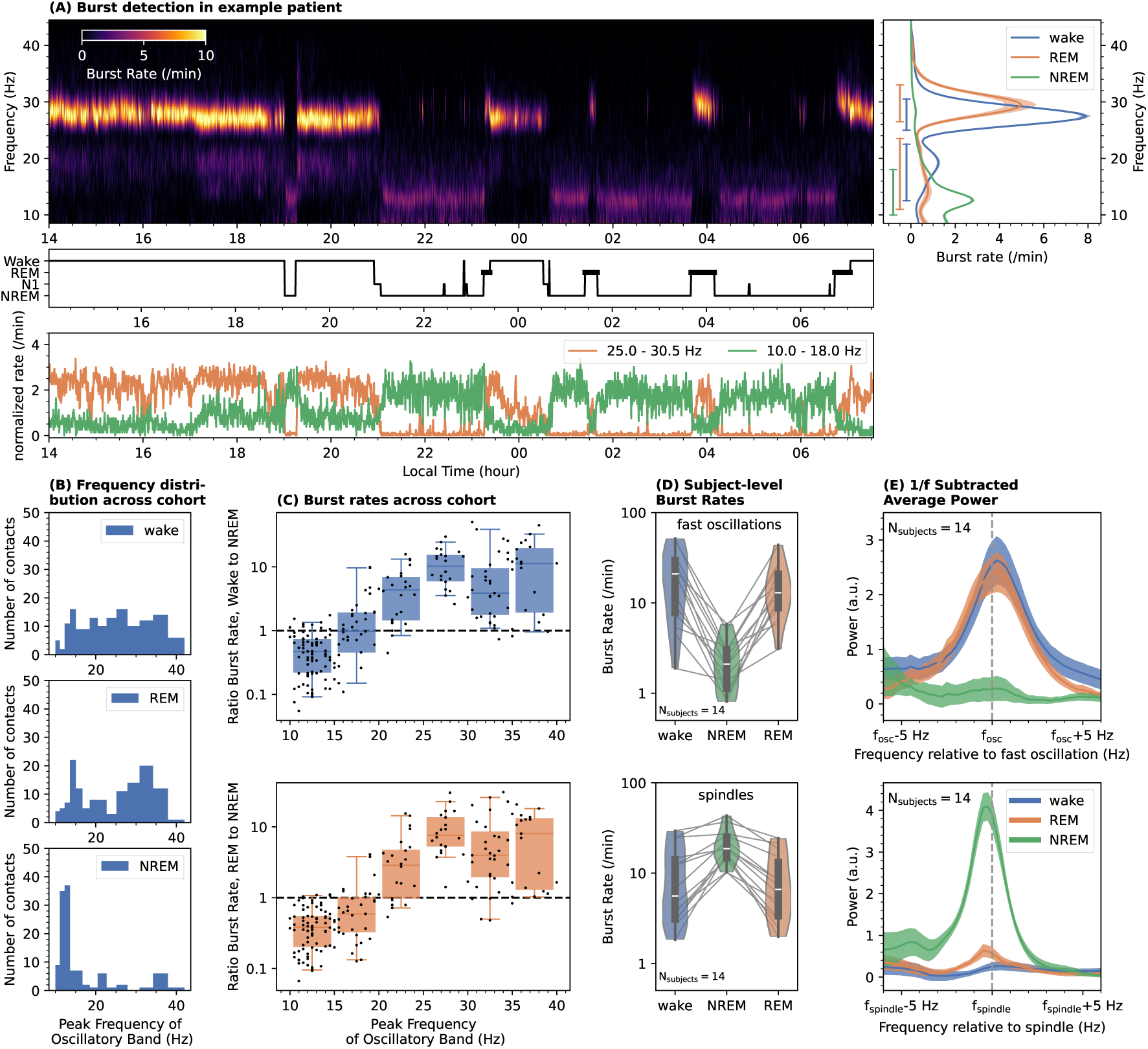
Thalamic oscillatory activity distinguishes natural brain states. Panel **(a)** shows an example illustrating our procedure for detecting frequency bands with significant oscillatory activity. The top panel on the left shows, in color, the burst rate as a function of time and frequency for the same thalamic channel as in Figure 1(b). The middle panel shows the sleep stage classification during the recording length. The panel on the right shows the average burst rate in the three brain states of interest: wakefulness, REM, and NREM. The shaded regions around the curves indicate the 99% confidence interval, derived using bootstrap resampling. The bars on the left of the panel show the frequency ranges detected to have significantly elevated burst rates. Finally, the normalized burst rate, as a function of time, in the spindle frequency band and the band with the fast thalamic oscillations is shown in the bottom panel of (a). Panel **(b)** shows the frequency distribution of N=213 oscillatory bands in thalamic contacts across our entire cohort, again for the three brain states of interest. Panel **(c)** shows, as a function of frequency for the detected oscillatory bands, the ratio of burst rate in wakefulness (top) and REM sleep (bottom) to that in NREM sleep. In both panels **(c)** and **(d)**, the boxes extend from the first quartile to the third quartile, with the line indicating the median; the whiskers indicate either the full range of the distribution or 1.5 times the interquartile range, whichever is smaller. Panel **(d)** shows the subject-level average burst rates, as a function of brain states, in frequency bands of the fast thalamic oscillations (top) and of NREM sleep spindles (bottom) in the 14 patients where we detected the fast thalamic oscillations. Panel **(e)** shows the subject-level average 1/f subtracted power spectra for the fast thalamic oscillations (top) and NREM sleep spindles (bottom). The shaded regions around the curves show the standard error of the mean. The x-axes on these panels indicates the frequency relative to the frequency of the peak burst rate. Both panels (b) and (c) demonstrate the high thalamic oscillatory activity in wakefulness and REM sleep, at frequencies above the typical spindle frequencies. The same can be seen at the subject level in panel (d). The presence of an oscillatory peak during wakefulness and REM sleep is clear on the top panel of (e).

The distribution of peak frequency of all oscillatory activity detected in the thalamic contacts of our cohort is shown, separately for each of the three natural brain states, in Figure 2(b). In line with expectations, the figure shows that the oscillatory activity in NREM sleep is dominated by sleep spindles, with NREM oscillations at 11 − 17 Hz being ubiquitously detected in 87 of the 105 thalamic contacts in our study. Conversely, we find a significant detection probability of bands of oscillatory activity at frequencies higher than the typical spindle band (19 − 45 Hz) during both REM sleep (0.62±0.09 per thalamic contact, subject-level mean and standard deviation using block bootstrapping, see Methods) and wakefulness (0.69±0.09 per thalamic contact). Indeed, these detection probabilities were significantly higher than the probability of detecting an oscillatory band at 19 − 45 Hz during NREM sleep (p *<* 10*^−^*^5^, block bootstrapping with N=17). Finally, we note that an overwhelming majority (90%) of oscillatory bands detected between 19 − 45 Hz in REM sleep had overlapping oscillatory bands detected in wakefulness as well, i.e., most of the oscillatory activity in these frequency ranges was specific to both wakefulness *and* REM sleep.

Next, we analyzed whether the differences in the detection probabilities represented actual enhancements of oscillatory activity, i.e., burst rates, in the respective states. For this analysis, we computed, for each thalamic contact, the time-averaged burst rates during wakefulness, REM sleep, and NREM sleep, for all oscillatory bands detected in the particular contact. We did this irrespective of the state(s) in which the oscillatory band was detected. Figure 2(c) shows, as a function of frequency for each detected oscillatory band in each thalamic contact, the ratio of the burst rate in wakefulness (top panel) and REM sleep (bottom panel) to that in NREM sleep. The figure clearly shows that, for frequencies above the spindle band (19−45 Hz), the rate of oscillatory activity during wakefulness and REM sleep is significantly higher than in NREM sleep (p= 1.2×10*^−^*^4^ for both cases, group-level paired two-sided Wilcoxon Rank Test with 14 subjects: statistic (dof=13)=0 for both wake vs. NREM and REM vs. NREM). Conversely, in line with expectations, the oscillatory bands in the typical spindle frequency range have significantly reduced burst rates in both wakefulness and REM sleep, when compared to NREM sleep (p*<* 3×10*^−^*^4^ for both cases, group-level paired two-sided Wilcoxon Rank Test with 17 subjects: statistic(dof=16)=7, p= 2.9 × 10*^−^*^4^ for wake vs. NREM and statistic(dof=16)=0, p= 1.5 × 10*^−^*^5^ for REM vs. NREM). Overall, we identified significant Wake-and REM sleep- specific oscillations above the spindle frequency band in the thalamic field potentials of 14 out of the 17 patients in our cohort. Figure 2(D, top panel) shows, separately for the three brain-states of interest, the subject-level average burst rates in the 54 thalamic contacts where we detected the fast oscillatory activity (see Methods). The figure clearly shows the enhanced burst rates during both wakefulness and REM sleep compared to those in NREM sleep. The converse can be seen for the frequency bands of sleep spindles detected in the same thalamic contacts in Figure 2(D, bottom panel).

Oscillatory activity in neural time series is often detected as “peaks” in the power spectrum after subtracting out the scale-free aperiodic component^23^. We thus also checked if the frequency ranges for which we detected an elevated burst probability in the thalamic contacts correspond to peaks in the power spectra. Figure 2(E, top panel) shows the average 1/f subtracted power spectrum for the thalamic contacts across the 14 patients where we detected the Wake- and REM sleep-specific oscillations. The average was obtained after aligning the power spectra to their respective peak frequency of the burst rate (see Methods). The figure clearly demonstrates that during both wakefulness and REM sleep, the frequency ranges of the elevated burst probability correspond to a clear peak in the 1/f-subtracted power spectrum, centered at the frequency of the peak burst rate. However, no such peak is seen during NREM sleep. The bottom panel of Figure 2(e) shows the average 1/f-subtracted power spectrum, but now averaged after aligning to the peak frequency of NREM sleep spindles. As expected, the panel shows a clear oscillatory peak during NREM sleep but not during REM sleep and wakefulness.

### 2.2 Eye movements during REM sleep predict bursts of oscillatory activity in the human thalamus

Having established brain-state-specific oscillatory activity in the thalamus, we focused on REM sleep to understand how the fast oscillatory activity relates to microstates. REM sleep is thought to be comprised of two distinct microstates: (i) *phasic* REM, characterized by bursts of eye movements and an elevated arousal threshold and (ii) *tonic* REM, characterized by epochs of marked reduction in eye movements and a reduced arousal threshold^24^. Given that the oscillatory activity in the thalamus at higher frequency bands is present in both wakefulness *and* REM sleep, we hypothesized that the oscillatory activity might be closely linked to bursts of eye movements in REM sleep, during which the brain is thought to simulate wakefulness while being decoupled from actual sensory inputs^25,26^. To test the above hypothesis, we analyzed the correlation between the burst probability of the fast oscillations and eye movements during REM sleep. This approach entirely circumvents the need to define various thresholds for the classification of REM microstates and provides a more basic understanding of how the oscillatory bursts are tied to the fundamental behavioural readout of REM sleep, the rapid eye movements themselves. We detected eye movements during REM sleep from simultaneously recorded frontal electroencephalography (EEG) electrodes (bipolar channel F7-F8, see Methods for a detailed description of the detection algorithm). The middle panel of Figure 3(a, b, c) shows three example traces of the voltage on F7-F8 for three different patients. The traces illustrate the typical bursty nature of eye movements during REM sleep; overlaid on the panels are the eye movements (EMs) detected by our algorithm. The top panels in these examples show the simultaneous Morlet amplitudes in a thalamic contact, illustrating the enhancement of fast oscillatory activity during the burst of rapid EMs. We quantified the relation between rapid EMs and oscillatory bursts by computing the peri-event histogram of the timing of oscillatory bursts in the specific frequency band detected in the thalamic contact relative to the timing of the EMs. An important confound in interpreting such peri-event histograms is the probability distribution of rapid EMs themselves. We thus also computed the probability of other rapid EMs around each detected rapid EM. The bottom panels in Figure 3(a, b, c) show, for the three example thalamic contacts, both the probability of fast oscillatory bursts (orange curve) and rapid EMs (gray curve) relative to the time of rapid EMs. The panels show a clear enhancement in the burst probability around the rapid EMs, which strongly correlates with how the probability of EMs themselves is temporally distributed.

**Figure 3:**
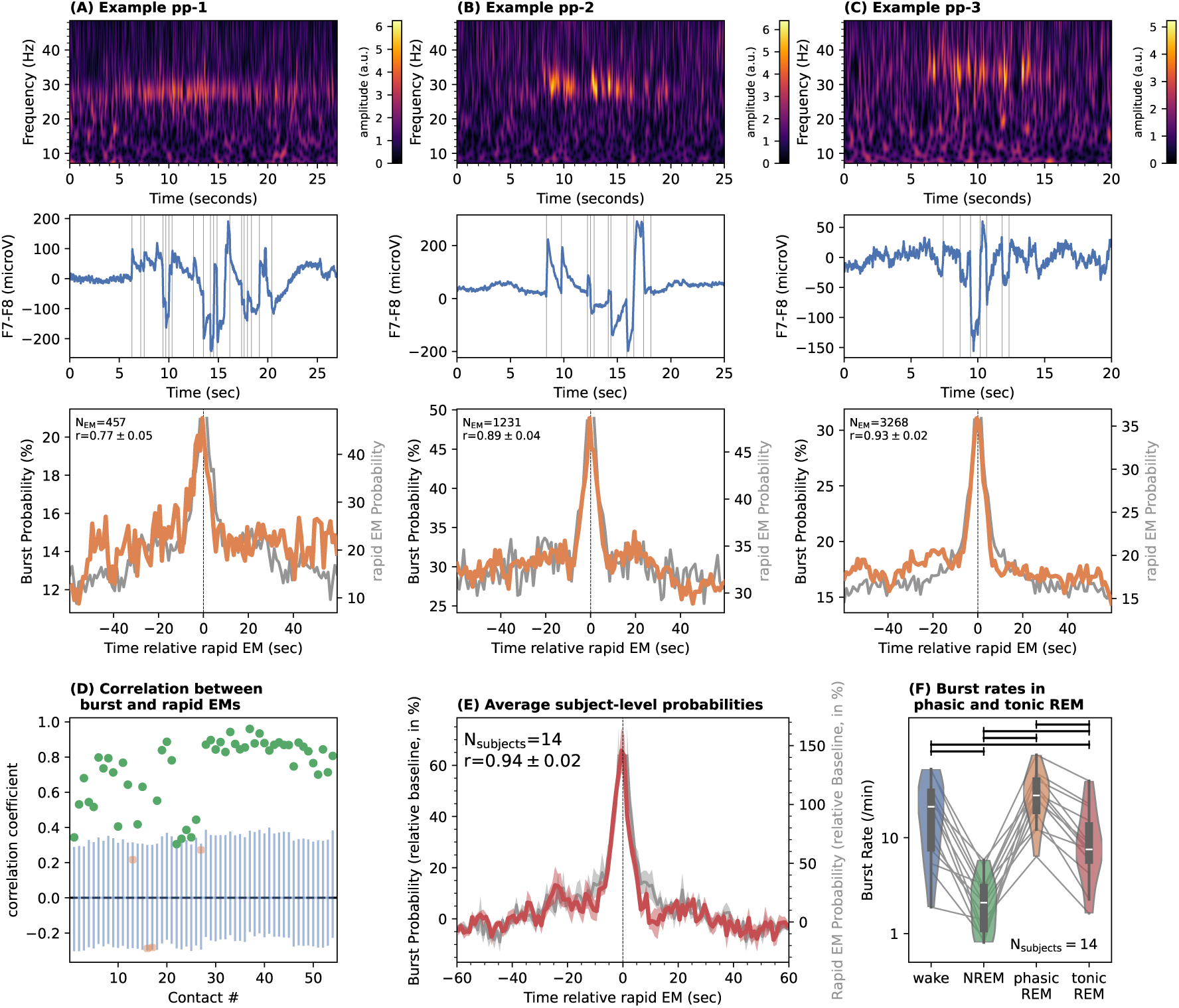
Bursts of the thalamic oscillations co-occur with eye movements during REM sleep Panels. **(a)**, **(b)**, and **(c)** illustrate the correlation between thalamic oscillatory activity and the bursts of EMs during REM sleep for three different patients in our cohort. The top panels show, in color, the Morlet amplitude as a function of time and frequency for an epoch of ongoing EMs detected in the frontal electrodes (shown in the middle panels, with detected rapid EMs marked with vertical lines). The bottom panel for each example patient shows, in orange, the average peri-event histogram of occurrence probability of thalamic bursts (in %) relative to the time of rapid EMs, for the entire duration of REM sleep in each patient. The grey curves indicate the probability distribution of other rapid EMs relative to the current EM. Panel **(d)** shows, for each of the 54 thalamic contacts across the 14 patients in which a Wake- and REM sleep- specific oscillatory band was detected, the correlation coefficient between the peri-event histogram of the thalamic bursts with the temporal distribution of the EMs themselves (e.g. the correlation coefficient between the orange and grey curves in panels A, B, C). The blue bars show the 99.91% intervals of null distributions of r values obtained using a Monte Carlo procedure (see Methods). Contacts with significant correlations (one-sided test for positive correlation using shuffling to estimate the null distribution, p*_corr_* < 0.05 after correcting for 54 comparisons using the Bonferroni method) are marked in green, while those without significant correlations are marked in red. The oscillatory bands were sorted by their peak frequencies. Panel **(e)** shows the subject-level averages (N=14) of the two peri-event histograms, with the shaded bands indicating the standard errors on the mean. The panel shows a clear enhancement of thalamic burst activity that is tightly coupled to eye movements during REM sleep. Panel **(f)** shows the distribution of burst rates of the fast thalamic oscillation as a function of brain state, after classifying REM into the phasic and tonic microstates. Significant differences in burst rates (two-sided Wilcoxon Rank Test at the group level, N=14, p*_corr_ <* 0.002, corrected for 6 comparisons using the Bonferroni method) between pairs of brain states are indicated with horizontal lines on the top. In panel **(f)**, the boxes extend from the first quartile to the third quartile, with the line indicating the median; the whiskers indicate either the full range of the distribution or 1.5 times the interquartile range, whichever is smaller.

Figure 3(d) shows the Pearson correlation coefficient between the probability of rapid EMs and oscillatory bursts for all thalamic contacts where we detected the Wake- and REM sleep- specific oscillations (see Methods). We find that ≈ 92% of the frequency bands show a significant correlation between burst probability and rapid EM probability (one-sided test for positive correlation using shuffling to estimate the null distribution, p*_corr_ <* 0.05 after correcting for 54 comparisons using the Bonferroni method, see Methods). We emphasize that this result is significant *individually* for each of the 14 patients in this analysis. Figure 3(e) shows the subject-level average thalamic burst probability and rapid EM probability, demonstrating a clear correlation between the two (*r* = 0.94 ± 0.02). We thus find that thalamic oscillations not only distinguish brain states at the macro level but also microstates at second timescales, specifically showing a significant enhancement in occurrence probabilities with epochs of enhanced rapid EM probability during REM sleep.

Finally, we also computed the burst rates after classification of REM sleep into the phasic and tonic microstates. The classification was done following criteria similar to those in the literature^13^, where epochs of REM sleep were further divided into 4-second-long segments. Segments containing at least two consecutive rapid EMs were labelled as Phasic REM, while those containing no rapid EMs were labelled as Tonic REM. Figure 3(f) shows that the average burst rate of the oscillation is significantly higher in phasic REM than in tonic REM (two-sided paired Wilcoxon Rank Test at the group level, N=14, p= 1.2 × 10*^−^*^4^, statistic=0). This is consistent with our finding that the burst probabilities are significantly elevated during rapid EMs. We further find that the burst rate of the oscillation during wakefulness is significantly higher than during tonic REM (two-sided paired Wilcoxon Rank Test at the group level, N=14, statistic(dof=13)=1, p= 2.4 × 10*^−^*^4^) but that it is not significantly different from the burst rate during phasic REM (two-sided Wilcoxon Rank Test at the group level, N=14, statistic(dof=13)=29, p= 0.15).

### 2.3 The detection probability of Wake- and REM sleep- specific oscillations is highest in the Central Thalamus

The thalamus is a diverse structure with its different subregions having specialized anatomical connectivity as well as functional roles^27^. Thus, an important question with regard to the wake- and REM- oscillations reported here is if they are specific to one of the thalamic subregions. Specifically, we hypothesized that the oscillatory signal is specific to the Central Thalamus, a key structure that is widely implicated in the regulation of arousal and states of consciousness^14^. We tested if the detection probability of the oscillatory activity above the spindle band that correlates with eye movements in REM sleep depends on the proximity of the electrode contact to the Central Thalamus via a univariate logistic regression (see Methods for detailed description and Supplementary Information for a detailed discussion on our analysis approach). We find that the detection probability of the thalamic signal is significantly correlated with the distance to the Central Thalamus (logistic regression slope= −0.41 ± 0.15, p = 0.0013, group-level statistic using block Bootstrapping with N=17, see Methods). We also performed a control analysis by repeating the logistic regression for the main clinical target of DBS in these patients, the Anteroventral (AV) nuclei. Figure 4 shows the best-fit regression for both thalamic regions. We do not find a dependence of detection probability on the proximity to the AV (logistic regression slope= 0.09 ± 0.10, p = 0.453, block Bootstrapping with N=17 patients). Further, we find that the slope of the logistic regression for proximity to the Central Thalamus is significantly steeper than that for the AV nuclei (p = 0.0026, group-level block Bootstrapping with N=17 patients, see Methods). Overall, our data is thus consistent with the hypothesis that the wake- and REM-sleep specific fast oscillations are specific to the Central Thalamus.

**Figure 4:**
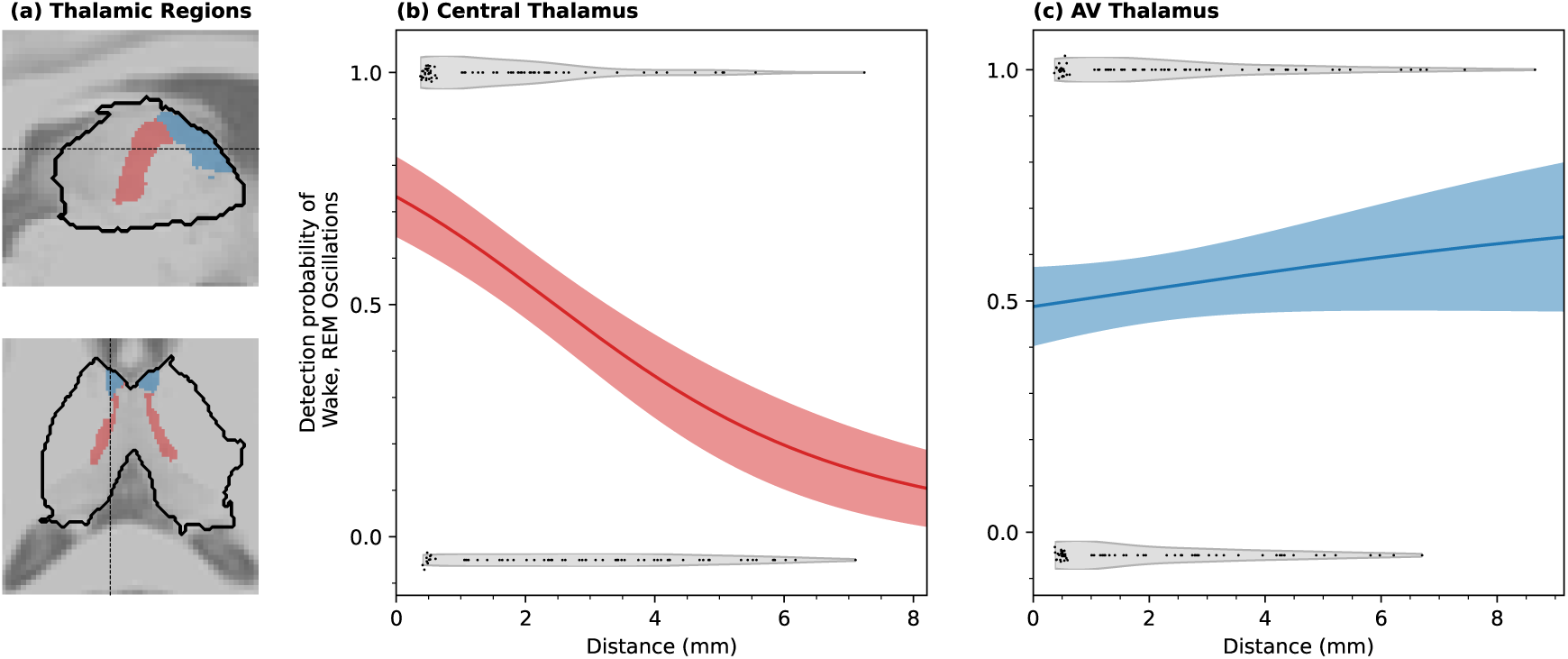
Central thalamus proximity predicts the detection probability of the Wake-and REM sleep- specific oscillations. Panel **(a)** shows, for visualization purposes, the location of the Central Thalamus (red) and the Anteroventral (AV; blue) overlaid on structural MRI slices. The dashed lines in each panel represent the line along which the slice shown in the other panel is taken. The middle and right panels show, for each of **(b)** Central Thalamus and **(c)** AV thalamus, the probability of detecting the Wake- and REM sleep- specific oscillations as a function of distance from the thalamic subregions. The solid curves in each plot show the logistic regression curve fitted to the data, with the shaded area indicating the estimated standard deviation. The black dots indicate our measurements; a y-axis value near 1.0 would imply that the Wake- and REM sleep- specific oscillatory signal was detected in the contact while a y-axis value close to 0.0 would imply no detection. The points clustered around 0.5 mm on the x-axis are those where the electrode contact was inside the subregion (see Methods); we added a scatter to these points to aid visualization. The violin plots around the points show the distribution of the distances. Note that all DBS contacts for all 17 patients were included in the analysis of this figure. The plot shows that the detection probability of the signal is strongly dependent on the distance of the contact to the Central Thalamus (logistic regression slope= *−*0.41 *±* 0.15, p = 0.0013, two-sided group-level estimate using block Bootstrapping where the distribution of surrogate cohorts was used to estimate the null distribution, N=17 patients, see Methods) but not on the distance to the AV (logistic regression slope= 0.09 *±* 0.10, p = 0.453 with N=17, two-sided group-level estimate using block Bootstrapping where the distribution of surrogate cohorts was used to estimate the null distribution, N=17 patients, see Methods).

We investigated whether the anatomical localizations for the DBS contacts of the 3 out of 17 patients in our cohort where we did not detect the Wake- and REM- sleep- specific thalamic oscillations, were systematically not in the Central Thalamus. Indeed, for two of these patients, there were no contacts localized to the Central Thalamus (see Supplementary Figure 11). It could thus be that the location of the DBS contacts affected the detectability of the fast oscillations in these two patients. The reason for not detecting the fast oscillation in the remaining patient remains unclear, given that the patient had two DBS contacts in the central thalamus. It is difficult to conclusively establish why this was the case, but one possibility could be that the signal is intrinsically much weaker in the particular patient.

### 2.4 Functional connectivity between the thalamus and the cortex as a function of global brain states

The thalamus is known to have extensive structural and functional connectivity with the cortex^27^. We used the simultaneously recorded scalp EEG data in our cohort to investigate whether the Wake- and REM sleep- specific thalamic oscillations can be detected on the scalp as either a direct power modulation or as a modulation of functional connectivity. First, we ran the same algorithm for the detection of frequency ranges with elevated probability of oscillatory bursts on each of the 10 scalp electrodes that were consistently available (see Methods). We found that only 4 out of the 14 patients showed clear evidence for a Wake- and REM sleep- specific oscillation in at least one scalp electrode and in a frequency range that overlapped with the frequency range of thalamic activity. Further, we also checked for oscillatory peaks in the 1/f-subtracted power spectra on the scalp electrodes. Extended Data Figure 1 shows the average 1/f-subtracted power spectra, after aligning the individual power spectra of each patient to the peak frequency of oscillatory activity in the thalamic iEEG channels (see Methods). The figure illustrates the lack of a clear oscillatory peak, as is seen in the thalamic contacts, during either wakefulness or REM sleep in any of the scalp EEG electrodes. Next, we checked for functional connectivity by computing the weighted Phase Lag Index (wPLI, see Methods) between the thalamic iEEG contacts and the scalp electrodes (see Methods).

The average wPLI for the 14 patients in the frequency range where the fast thalamic oscillatory activity was detected is shown in Extended Data Figure 2(a). The figure shows that the wPLI between thalamic contacts and scalp electrodes in these frequency ranges during REM sleep and wakefulness is significantly higher than during NREM sleep for all three scalp EEG regions (for all three effects, p_corr_ *<* 0.006, group-level paired two-sided Wilcoxon Rank Test with 14 subjects, corrected for 3 comparisons using the Bonferroni method: statistic(dof=13)=6, p_corr_=0.005 for Frontal, statistic(dof=13)=5, p_corr_=0.003 for Central, and statistic(dof=13)=3, p_corr_=0.002 for Occipital). We also repeated the above for the frequency ranges of NREM sleep spindles in the thalamic contact (Extended Data Figure 2(b)). For all three groups of scalp electrodes, we find that the wPLI in frequency ranges of sleep spindles is significantly higher during NREM sleep than during REM sleep and wakefulness (for all three effects, p_corr_ *<* 0.002, group-level paired two-sided Wilcoxon Rank Test with 14 subjects, corrected for 3 comparisons using the Bonferroni method: statistic(dof=13)=0, p_corr_=0.0004 for Frontal, statistic(dof=13)=2, p_corr_=0.001 for Central, and statistic(dof=13)=2, p_corr_=0.001 for Occipital). Finally, Extended Data Figure 2(c) shows the clear presence of a peak in the frequency-resolved wPLI at the peak frequency of the oscillatory activity detected in the thalamic contacts. We thus find clear evidence for brain-state-specific thalamocortical connectivity in the frequency ranges of the fast oscillations during REM sleep and wakefulness as well as of sleep spindles during NREM sleep (see also Supplementary Information for results and discussion of a Granger Causality analysis).

We note that coherence measures such as wPLI have been shown to depend on power and phase-locking in the sender, as well as connectivity between the two regions^28^. Assuming that the thalamus is the sender in our case, there could be coherence between the regions without a clear increase in power in the receiver (the cortex). This could happen, for example, if the signal is only weakly detectable in the scalp EEG, and the phase information from the thalamic signal helps boost its detectability when computing coherence. However, we would like to emphasize that coherence between two regions does not necessarily imply neuronal communication between them.

## 3 DISCUSSION

The central finding of this study is the detection and characterization of oscillatory activity in the human thalamus, at frequencies above the spindle band, that specifically occur during two natural states of consciousness - wakefulness *and* REM sleep. We find that bursts of the thalamic oscillations correlate strongly with phases of rapid eye movements during REM sleep; these oscillations thus further distinguish between the phasic and tonic microstates in REM sleep. Finally, we find that the detection probability of the oscillatory activity is significantly enhanced when the DBS contact is close to the Central Thalamus, a region of the thalamus that has previously been implicated in regulating states of consciousness.

A clear electrophysiological signature for the state of consciousness in the Central Thalamus of humans opens up promising lines of investigation in both understanding the contributions of the thalamus to states of consciousness as well as in refining clinical interventions, such as thalamic DBS, for disorders of consciousness. The Central Thalamus has been shown to play a key role in arousal regulation^14,16^. Research in non-human primates shows that DBS of central thalamic nuclei can reverse the loss of consciousness in anesthetized animals^17–19^ as well as improve behavioral performances by regulating arousal levels^15,29^. Indeed, in two patients where we had simultaneous recordings with video-based eyetracking available, we find a significant correlation between the burst rate of the fast thalamic oscillation and the pupil diameter, a widely used metric of arousal^30^ (see Extended Data Figure 3 and Methods). This clear correlation between pupil diameter and thalamic burst rates suggests that the fast thalamic oscillations are not only related to global brain states but also to levels of arousal within wakefulness.

In humans, several clinical studies show that DBS of the Central Thalamus can lead to improvements in patients suffering from disorders of consciousness [see refs.^20,21,31^ for reviews]. Interestingly, one of these studies^32^ reported the progressive emergence of REM sleep in a minimally conscious patient after 18 months of Central Thalamus DBS. Despite the convincing causal evidence for the role of the Central Thalamus in regulating arousal and consciousness, a mechanistic understanding remains elusive^14,16,33^. Our discovery of a fast oscillation specific to the Central Thalamus during wakefulness raises the possibility of a causal connection between the DBS-triggered arousal described above and the activation of the central thalamic oscillatory activity. Indeed, previous research^17^ has found that the DBS frequency most effective in causing arousal in anaesthetized animals of 50 Hz approximately matched the frequency of thalamic activity when animals were awake; these authors thus speculated that the DBS was effectively “mimicking” the intrinsic activity during wakefulness. However, there is no broad consensus on how the efficacy of DBS in causing arousal depends on the stimulation frequency. This is also the case for the clinical use of thalamic DBS to reverse the loss of consciousness in humans, where the efficacy of the intervention varies widely with reported studies [see ref.^21^ for a meta-analysis]. If a potential mechanism via which thalamic DBS causes transitions in conscious states is mimicking the intrinsic oscillatory activity of the thalamus, the efficacy of DBS might critically depend on the relation between (i) the stimulation frequency and the intrinsic frequency of the wakefulness-specific oscillations and (ii) the stimulation site and the source of the oscillations. Future studies should thus systematically study how DBS at different frequencies and in different thalamic nuclei impacts the intrinsic fast oscillations. In addition, a characterization of oscillatory activity in the thalamus of patients with disorders of consciousness might provide valuable insights into how future therapeutic interventions could be optimized for different patient cohorts (see, e.g., ref.^34^).

The distance to the Central Thalamus significantly predicted the detection probability of Wake-and REM sleep- specific oscillatory bursts. This was not the case for any of the other thalamic regions that were sampled by our data (AV, Mediodorsal medial, Ventral Anterior, or Ventral Lateral Thalamus). However, it is important to note that the specificity of the signal to the Central Thalamus does not necessarily implicate the region as the source of the oscillation. For example, the generation or sustainment of the oscillation may depend on thalamocortical networks. In this context, it is interesting to note that research with model animals (specifically cats) shows that thalamocortical networks can produce fast oscillations, both *in vivo* and *in vitro*^35,36^. The typical frequencies of the oscillatory activities, as well as their brain-state specificity^37,38^, are similar to the oscillatory activity we detect in the human thalamus. Interestingly, these studies indicate that the thalamocortical networks that maintain sleep spindles can switch to sustaining faster rhythms depending on the type of ion channels that were activated^1,2^ and different neuromodulatory conditions, e.g., driven by cholinergic input from the brain stem^39^ . Further, computational modeling of thalamocortical networks also produces similar transitions in dominant oscillatory activity from spindle-like oscillations under neuromodulatory conditions of NREM sleep to faster oscillations in the beta-gamma frequency ranges under conditions of wakefulness^40^. Indeed, it is interesting to note that we find the amplitudes of NREM sleep spindles and the wake- and REM- specific fast oscillations to be correlated across contacts and patients, indicative of an underlying relationship between these oscillations (see Supplementary Figure 2).

We found a strong correlation between epochs of rapid EMs in REM sleep and the burst probability of the thalamic oscillation. The burst rates of the fast oscillation were thus different between the two REM microstates, phasic and tonic, with an elevated burst rate in phasic REM. We note that Simor et al.^13^ also used DBS electrodes in the thalamus of patients with pharmacoresistant epilepsy to study differences in power spectra between tonic and phasic REM sleep. However, a direct comparison between their and the results of this work is made difficult by the very different focuses of their study and this work; the primary being (i) Simor et al. focused on the overall power spectrum in the two REM microstates, after averaging across subjects, while we focused primarily on oscillatory power across multiple brain states (NREM, wake, tonic and phasic REM) at the individual subject level; our burst detection technique to differentiate between oscillatory power and aperiodic power allowed us to detect patient-specific frequency bands with elevated oscillatory power and (ii) Simor et al. investigated the power spectra in the anterior thalamus only, while we included all available thalamic contacts, and found that the probability of detecting the oscillatory activity depends on the distance to the Central Thalamus but not the Anterior Thalamus.

The thalamus is a key regulator of brain states, controlling the flow of information between the sensory periphery and the cortex as well as between different regions of the cortex^41^. It is thus intriguing that the mesoscale activity of the thalamus is similarly synchronized via fast oscillations across two very different states of consciousness: wakefulness, where alertness to external sensory information is maintained, as well as REM sleep, where external sensory information is processed differently (see e.g., ref.^42^). Our observations here would be consistent with the hypothesis that the brain is involved in a simulation of actions and their consequences during REM sleep^26^. For example, studies in rodents show that eye movements during REM sleep meaningfully update the animal’s head direction system, similar to actual head turns during wakefulness^25^. A potential function for the thalamic oscillations reported here would thus be in mediating action-related information. Indeed, research in non-human primates has shown a key role of higher-order thalamic nuclei, such as those in the Central Thalamus, in communicating efference copies of actions^41,43^. Specifically, during REM sleep, the fast thalamic oscillations could synchronize the flow of information related to internal simulations of actions in ways similar to those during wakefulness. In addition, the Central Thalamus has regions that overlap with the “occulomotor thalamus” which is known to play a role in the generation of eye movements in animals^44^. It is thus possible that these oscillations are also related to the generation of eye movements during REM sleep. Overall, these hypotheses would be consistent with our findings that the oscillatory activity is significantly enhanced during bursts of eye movements in REM sleep.

A potential limitation of the study is the possibility of abnormalities in thalamocortical structural connectivity[45], which have been described previously for such cohorts with pharmacoresistant epilepsies. Further, administration of anti-seizure medication (ASM) is also known to have an impact on sleep architecture [46]. As with any study that utilizes intracranial data from such cohorts, the extent to which the results generalize to healthy populations remains an open question. However, multiple factors, such as (i) the relatively large size of our cohort (compared to similar studies), (ii) the use of group-level statistics throughout the study, (iii) the heterogeneity of epilepsy types (see Extended Data Table 1), (iv) the lack of correlation between thalamic burst rates and seizure rates as well as between the burst rates and epilepsy type (see Supplementary Figure 4), and (v) heterogeneity of ASM regimes (see Extended Data Table 1), makes it unlikely that the main findings of this work are driven by pathology.

An important question for future research would be to understand how the thalamic oscillations reported here compare with cortical oscillatory activities reported at similar frequencies that researchers have hypothesized to be thalamocortical in nature. Specifically, the thalamic oscillations bear interesting resemblances to 40 Hz oscillations detected in Magnetoencephalography during REM sleep^47^, beta-band power enhancements^48^ and oscillatory activity in REM sleep^49,50^, beta bursts during wake^51^, and narrowband EEG oscillations at 25-35 Hz in children and young adults^52^. We note that while our current results show a clear modulation of brain-state-specific functional connectivity at the frequencies of thalamic oscillations, the low-density and sparse coverage of the simultaneously recorded scalp EEG was a limitation of our study. A clear understanding of the connection with cortical activity may only be possible with simultaneous intracranial cortical and central thalamic recordings.

While our study establishes that thalamic oscillations are correlated with different states of consciousness and provides evidence that the fast thalamic oscillations reported in this work are related to arousal levels in wake, it is important to note that the connection of the oscillation to the broader phenomenon of consciousness, specifically with *contents* of consciousness^16,53^ remains an open issue. Future experiments specifically designed to address this important open issue would be of much interest. If the thalamic oscillation reported here also relates to the contents of consciousness, it may help to carve out potentially conscious contents from brain states that are notoriously difficult to access without disruption. REM sleep, and particularly phasic REM, has been reported to produce vivid dreams that involve conscious experience^54^. That the thalamic oscillation reported here is specific to phasic REM and wake might thus speak in favor of the two states being similar with respect to conscious access. Further, our discovery could have implications for the efficacy and side effects of thalamic DBS therapies for various disorders^34,55,56^. Finally, our findings inform current theories of the neural correlates of consciousness^57–60^ by highlighting a critical role of the Central Thalamus and may also provide testable predictions about how to interfere with and causally test them^61^.

## 4 METHODS

### 4.1 Participants

Simultaneous intracranial and scalp EEG recordings were obtained from pharmacoresistant epilepsy patients bilaterally implanted with DBS electrodes for chronic stimulation of their Anterior Thalamus^62^. The data was recorded at the Epilepsy Center, Department of Neurology, Ludwig–Maximilian Universitäat (Munich, Germany), during post-surgical externalization of the DBS electrodes, prior to connecting them to the pulse generator. Participants did not receive compensation for their participation.

No statistical methods were used to pre-determine sample sizes, but our sample sizes are larger than those reported in previous publications^13^.

Our primary inclusion criterion was the availability of intracranial thalamic recordings with at least one full night of sleep and daytime wakefulness. The recordings for a total of 24 patients implanted at the center met these criteria.

We excluded 7 patients from the main analysis due to various reasons. These exclusions can be broadly classified into four categories: (i) one patient with reduced vigilance, (ii) two patients due to excessive Epileptic Spiking in the scalp electrodes and the inability to confidently classify sleep stages, (iii) three patients due to very short duration of REM sleep (≲ 10 minutes) in the only night of recording that was available, and (iv) one patient with clear recording system malfunction during the night of recording. Overall, our main analysis is carried out on a total of 17 patients (see Extended Data Table 1 for details on patient characteristics).

We note that for the one patient with reduced vigilance, we could not find clear signatures of different sleep stages when looking at the scalp EEG. Further, the thalamic field potentials also showed no clear sign of spindles in the 16 hr of data that were recorded from the patient. While we did not see a clear indication of the presence of a fast thalamic oscillation in this patient, the thalamic field potential does show some switching between slow oscillation-dominated epochs and epochs with some enhancement in higher frequency power. However, the inability to unambiguously classify sleep stages based on scalp EEG makes it difficult to draw any conclusions.

One patient in our main cohort was reimplanted due to clinical reasons. The DBS electrodes were externalized after both implantations. The recorded data in both epochs met our inclusion criteria. We thus include the data from both epochs, but given that the contact locations were changed in the reimplantation, we treat the data from the two epochs as arising from separate thalamic channels.

The data for seven patients in our main cohort partially overlap with the data used in a previous study^11^.

The study received permission from the local ethics committee (Ethics Committee of the Faculty of Medicine, LMU Munich, project number 24-0454) and all participants consented to participation in the study.

### 4.2 Data Acquisition

The patients in this study were implanted transventricularly with two DBS electrodes, one for each hemisphere, and with each electrode having four intracranial contacts (platinum–iridium contacts, 1.5 mm wide with 1.5 mm edge-to-edge distance; Medtronic 3387). The data was recorded using XLTEK Neuroworks software (Natus Medical, San Carlos, CA, USA) and an XLTEK EMU128FS amplifier. Electrodes placed on the scalp were recorded simultaneously. The scalp electrodes were positioned according to the international 10-20 system. However, the scalp EEG coverage was variable with the number of available scalp electrodes varying between 11 and 22. Despite the variability, a total of 10 scalp EEG electrodes were consistently available for all except two patients in the main cohort. The consistently available scalp EEG channels were: (i) Four frontal electrodes F7, F8, Fp1, and Fp2, (ii) Four central electrodes P7, P8, T7, and T8, and (iii) Two occipital electrodes O1 and O2. For two patients in the cohort, all of the above electrodes except Fp1 were available. All channels were originally referenced to the scalp electrode CPz and sampled at 200 Hz (1 patient), 256 Hz (9 patients), and 1024 Hz (7 patients).

### 4.3 iEEG and scalp EEG Preprocessing

All EEG analyses were performed using the MNE toolbox^63^, with additional custom routines written in Python. The recordings for all patients were downsampled to a uniform sampling rate of 200 Hz, using an anti-aliasing filter. A notch filter was used to suppress power at the electrical power line frequency of 50 Hz. The intracranial contacts were bipolarly rereferenced with neighboring contacts, producing a total of six intracranial channels per patient (L1-L2, L2-L3, L3-L4, R1-R2, R2-R3, R3-R4). For one patient, only one of the two DBS electrodes was connected; the voltages from the electrode in the right hemisphere were not recorded. The online reference Cpz was retained for the scalp EEG channels.

We used an automated algorithm to detect Interictal epileptic discharges (IEDs) in each bipolarly referenced intracranial channel. We high-pass filtered the data above 55 Hz, followed by a Hilbert transform to obtain the envelope of the filtered data. Next, we marked all epochs that exceeded five times the median amplitude of the envelope. We marked a period ±1 sec around each event as an IED. The mean fraction of excluded data was 3.6% per channel. We note that the range of frequencies above 55 Hz was selected to avoid misclassification of thalamic oscillations between 19-50 Hz as IEDs. We further emphasize that our burst detection algorithm (outlined in a later subsection) has built-in safeguards, such as a maximum frequency width for bursts, that would minimize the likelihood of detecting IEDs as oscillatory bursts.

### 4.4 Sleep Staging

We carried out sleep staging using standard procedures^64^, with sleep stages N2 and Slow Wave Sleep (SWS) combined into NREM. The sleep staging was carried out visually in standard 30-second segments by looking only at activity on the scalp EEG channels. The thalamic iEEG channels were not used to sleep score to avoid potential confounds and confirmation biases. The standard AASM guidelines on looking for characteristic oscillations in different sleep stages were followed (spindles, K-complexes in N2 sleep, the presence of slow oscillations in N3 sleep for more than 20% of the 30s epochs in N3 sleep, and signatures of eye rolling and reduction in alpha power for N1). Signatures of eye blinks on the scalp EEG were further used as an indicator for wakefulness. We also note that special attention was paid to ensure that the classification of REM sleep is highly reliable, and that there is no misclassification of REM stages with periods of arousals or NREM sleep; specifically, we ensured that REM epochs overlapped with periods of clear eye-movement-related activity on the frontal contacts, that no characteristic NREM specific oscillations (spindles or K-complexes) were present, and that there were no indications of eye blinks on the frontal scalp electrodes throughout the entire epoch.

### 4.5 Burst Detection from Morlet Spectrogram

Detection of oscillatory bursts, such as spindles, is typically done on a pre-defined frequency band by thresholding on a bandpass filtered and Hilbert transformed time series (e.g., ref.^11^). In this work, we used an improved burst detection algorithm that requires no prior knowledge of frequency band(s) where significant neural oscillations are present. This is critical to the current work as there was substantial individual variability in the frequency of the fast thalamic oscillation during REM sleep and wakefulness.

We performed, for each bipolarly referenced channel, a discrete Morlet transform on the entire time series, with a wavelet width of 10 cycles, over a frequency range of 3-49 Hz (sampling every 0.5 Hz), and with a temporal sampling of 5 ms (equivalent to our downsampled rate of 200 Hz). The Morlet wavelets used in the transform have a normalization that preserves the amplitude of input oscillations, i.e., the transform of sinusoidal waves with identical amplitude would also yield identical amplitudes in the transformed space, irrespective of the frequency of the sine waves. We note that the Morlet transform is computed after masking out time segments of the data affected by IEDs in the particular channel.

We next detected peaks on the two-dimensional Morlet amplitudes that exceeded a predefined threshold of two times the median of all Morlet amplitudes across all time samples and frequencies for the given channel. The temporal and spectral extent of each event detected at this stage was obtained from cuts in the 2D space along both the time and frequency axes. The width of an event was defined using the first instance along each dimension where the Morlet amplitude around these cuts meets any of the following criteria: (i) the amplitude was less than twice the median, or (ii) the amplitude was less than 10% of the peak amplitude, or (iii) the amplitude turns over and starts increasing. Finally, any detected neighboring events with similar peak frequencies (≤1 Hz) such that the stopping point of one and the starting point of the next burst were within 1.5 cycles of each other were merged into a single event.

To ensure that the detected events represent actual oscillatory activity, rather than noise fluctuations or broadband and aperiodic increases in power, we imposed additional criteria that the detected bursts must be (i) at least five oscillatory cycles long, (ii) have a peak amplitude that exceeded three times the median of all Morlet amplitudes in the channel, and (iii) that the width of the burst along the frequency axis was between 2 Hz and 20 Hz. Further, we excluded any detected burst likely to be a primary harmonic of a lower frequency burst; this was accomplished by searching for any coincident burst with a higher amplitude and at half the frequency of the burst. Finally, any pair of bursts with overlap along both the spatial and temporal dimensions was marked as a duplicate, and only the burst with a higher amplitude was retained. Extended Data Figure 4 shows an example that illustrates our burst detection algorithm.

### 4.6 Defining Oscillatory Bands in Different Brain States

We binned the oscillatory bursts detected from Morlet Spectrograms (as described above) to obtain burst rates as a function of time and frequency, for the entire duration of the recordings. The width of the bin along the time axis was 30s, matched to the resolution of sleep scoring^64^. Along the frequency axis, the bin width was 0.5 Hz, and the burst rates were smoothed using a Gaussian kernel with Full Width at Half Maxima (FWHM) of 2 Hz. An example 2D histogram from a single patient and bipolar contact is shown in Figure 2(a).

We averaged the 2D burst rate distribution along the temporal axis to obtain an average burst rate for each of the three brain states of interest: wakefulness, NREM sleep, and REM sleep. To ensure that the oscillatory bands defined represent the actual enhancement of burst probability over the given range of frequencies, we perform peak detection on the average burst rate as a function of frequency, for each bipolar contact and each of the three brain states separately. We only included peaks with a peak burst rate greater than 0.1 bursts/min. The width of the peaks was determined using a combination of the slope of the average burst rate as a function of frequency, as well as the burst rate curve itself. Specifically, around each frequency peak, we first searched for a peak in the first derivative on either side. Next, on the lower (or higher) side of the detected first-derivative peaks, we searched for the first frequency bin where either the burst rate drops to 25% of the peak burst rate or the burst rate decreases by less than 1% per Hz, whichever occurs earlier. Further, we merged any detected bands without a clear trough separating them, such that the ratios of the peak burst rates to trough burst rate were less than 1.2. Finally, we excluded any detected frequency range with a width of less than 1 Hz on either side of the peak frequency. We obtained a total of 591 frequency bands with enhanced burst probability across all states, bipolar contacts, and patients (i.e., an average of ≈ 2 oscillatory bands per bipolar contact per brain state).

Next, we checked if the oscillatory bands thus detected represented frequencies with elevated burst probabilities that were statistically significant. In order to do so, we estimated the errors on the burst rates via bootstrap resampling with replacement, with 10^6^ random realizations. The p-value of the significance of the detected frequency ranges was obtained by computing the fraction of bootstrap realizations where the burst rate at the detected peak frequency was greater than the burst rate at the edges of the frequency band. We find that 409 frequency ranges, across all states, bipolar contacts, and patients, have significantly enhanced burst rates (p-value *<* 0.05*/*591 ∼ 8 × 10*^−^*^5^, correcting for 591 comparisons).

Finally, we removed any frequency band that is very likely to represent harmonic bursts by searching for any lower frequency band that (i) was detected in the same brain state but at sub-harmonic frequencies (specifically, 1/2 and 1/3), (ii) had a higher burst rate, and (iii) had a burst rate that was significantly correlated with the burst rate of the potential harmonic frequency (spearman rank correlation, p*<* 10*^−^*^3^). We note that although our original burst detection routine substantially eliminates detecting bursts at harmonic frequencies, noise fluctuations cause a small fraction of these to get detected. Overall, we found 15 out of the 409 frequency bands that met the above criteria for representing bursts at harmonic frequencies. We remove these to obtain a final set of 394 frequency bands. The frequency distributions of these 394 bands, separately for the three brain states of interest, are shown in Figure 2(b).

We emphasize here that our overall approach, as described above, was to first select frequency bands that may have significantly elevated burst rates based on actual measurements of burst rates and then subject the selected frequency bands to tests for significance. This ensures that the initial detection and characterization of the peaks in burst rate is done independent of any statistical significance criteria, avoiding the common pitfall of obtaining widths using cluster-based tests, such as an artificial modulation of the width that depends on the noise (see e.g ref.^65^).

We determined, for each thalamic contact separately, if a given frequency band was detected in more than one brain state. In such cases, we combined these into a single frequency band, retaining the detection with the narrower frequency width (irrespective of brain state). After combining overlapping bands across brain states, we obtained a total of 213 frequency bands across our cohort (an average of ≈ 2 bands per bipolar thalamic contact). The ratio of burst rates across different brain states for these 213 frequency bands is shown in Figure 2(c).

Finally, for each of the 105 thalamic contacts, we identified the frequency bands of the wakefulness and REM sleep-specific oscillations as those with (i) a peak frequency above the spindle band (*>* 19 Hz), (ii) an elevated burst probability detected in both wakefulness and REM sleep, and (iii) a higher mean burst rate in both wakefulness and REM sleep compared to that in NREM sleep. We find such frequency bands in a total of 54 thalamic contacts across 14 patients, with 7 thalamic contacts having two such distinct frequency bands. For the 7 contacts with two frequency bands, we consistently used the higher frequency band for all analyses. We note that none of the conclusions of this paper depend on this choice; for example, including both frequency bands led to identical conclusions. The subject-level average burst rates, as a function of brain states, for these oscillatory bands are shown in Figure 2(D, top panel). We note that these 54 thalamic contacts across the 14 patients entered further analysis (e.g., that of Figure 3). For comparison, the subject-level average burst rates at spindle frequency bands in the same bipolar channels are shown in Figure 2(D, bottom panel). We note that a frequency band with significant spindle oscillation was identified in 52 of these 54 bipolar channels.

We note that we also detected oscillatory activity in the thalamus during wakefulness or REM sleep that lies in typical alpha and low-beta frequency ranges^66,67^; these can be seen in the distribution of frequencies in Figure 2(b). The frequency ranges of alpha and low-beta partially overlap with the spindle frequency band. These oscillations thus contribute to the burst rates in the spindle frequency ranges during REM sleep or wakefulness. However, this does not impact the analysis or the interpretation of any of the results of this paper.

### 4.7 Block Bootstrap Resampling

Bootstrap resampling with replacement is a widely used resampling technique to estimate measurement uncertainties without having to assume a particular form of the underlying probability distribution [see e.g. ref.^68^ for a discussion specific to neurophysiological data]. Block bootstrapping is an extension of bootstrap resampling with replacement for data that contains correlated measurements. This is especially well-suited for computing group-level statistics for various quantities in this work.

We treated measurements from each subject as one block during the resampling procedure. Specifically, for each Monte Carlo iteration, we obtain a surrogate cohort by randomly drawing a set of subjects equal to the original number of subjects, i.e., effectively performing a bootstrap resampling with replacement at the subject level. We then measure the quantity of interest for these surrogates. This procedure is repeated 10^5^ times to obtain a probability distribution function for the quantity of interest.

When testing the hypothesis that the probability of detecting an oscillatory band at 19 − 45 Hz is greater in REM sleep (wakefulness) than in NREM sleep, we treated each subject as one block. We thus estimated for each surrogate cohort the detection rate in REM sleep (wakefulness) and NREM sleep. The p-value of rejecting the null hypothesis was then obtained by calculating the fraction of surrogate cohorts where the detection rate in NREM sleep was higher than that in REM sleep (wakefulness). We also used the surrogate cohorts to estimate the group-level standard deviations of the detection rates.

### 4.8 Detecting Eye Movements from Frontral Electrodes

We detected horizontal rapid eye movements using the frontal electrodes F7 and F8. We used the bipolar derivative F7-F8 as a substitute for horizontal electrooculogram (EOG). The detection of horizontal REMs from F7-F8 was done using a series of thresholding on both the voltage deflection as well as the rate of change of the voltage deflection. We first bandpass filtered the voltages above 0.1 Hz and below 40 Hz. Next, we computed the rate of change of the voltage deflection (hereafter “velocity”) over a range of temporal scales spanning 20 ms to 250 ms, in steps of 10 ms; this was done such that the detector is sensitive to both fast and slow voltage deflections.

The velocity at each temporal scale required to be z-scored before any thresholding could be carried out. A standard z-scoring procedure over the entire duration of the recording was not found to be suitable for our purposes because (i) the mean and standard deviation of the signal evolve over the very long recording durations of our data and (ii) the mean and standard deviation of the signal would be substantially affected by the presence or absence of an eye-movement-related deflection, thereby limiting the sensitivity of the detector. We thus used a robust and temporally resolved approach to effectively z-score the data by computing the moving median absolute deviation of the signal with a sliding window of 5 minutes.

For each temporal scale, an initial selection of horizontal EM events was defined as those that exceeded two times the median absolute deviation for a duration between 20 ms and 750 ms **and** with a peak velocity that was at least seven times the median absolute deviation. The first and last time instances where the velocity exceeded two times the median absolute deviation were defined as the onset and offset of the event, respectively.

In order to ensure that the detected REMs were highly reliable, we further subjected the initial detections to a series of additional thresholding to arrive at a final set of events. These criteria were essential to avoid including other events, such as IEDs that also have sharp transitions. The additional criteria were as follows.

1. The peak velocity exceeded five times the standard deviation of the velocities ±200 ms around the onset and offset of the event.
2. The magnitude of voltage deflection exceeded five times the standard deviation of the voltages ±200 ms around the onset and offset of the event.
3. The maximum voltage deflection ±200 ms around the onset and offset of the event was less than the voltage deflection associated with the event itself.

The above detection was run on all the velocity time series at all scales (20-250 ms). Events detected at multiple scales were merged together into a single event, with the onsets and offsets derived from the shortest timescale velocity series, so as to retain the most accurate timing information available.

The total number of rapid EMs detected in our cohort varied between 56 and 6964, with a mean rapid EM frequency of 8 ± 5 per minute. Further, the median number of EMs in our cohort is 1101, with 80% of the patients having more than 300 EMs.

The average topography on the scalp EEG of deflection at the detected rapid EMs is shown, for each patient separately, in Extended Data Figure 5. Despite the limited and variable scalp EEG coverage, the figure shows the stereotypical scalp topography associated with horizontal eye movements for each of the 17 patients in our cohort.

Further, for comparison with events detected in a standard horizontal EOG setup, we used data of 11 healthy participants from an independent study where reliable EOG measurements during REM sleep were available simultaneously with a standard 64-channel scalp EEG. We ran the algorithm as described above, with identical parameters, on both horizontal EOG and the bipolar derivative F7-F8. For every subject, we find that *>* 82% of the rapid EMs detected in F7-F8 were also detected in horizontal EOG, without any additional fine-tuning of the parameters for horizontal EOG.

Finally, as an additional comparison, we also used video-based eyetracking data that is available for two of the patients in our cohort (whose pupil data were used for the analysis in Extended Data Figure 3). The two patients had performed an unrelated task, with a total duration of 80-100 min per patient, where we had simultaneous eye-tracker data at 600 Hz, recorded with the Tobii Pro Eyetracker. For this analysis, we first ran the EEG-based eye-detection algorithm on the data from these patients. Next, we computed the eye velocity from the eye-tracking data. Extended Data Figure 6 shows the eye velocity data for the two patients, aligned to the events detected by our algorithm from the scalp-based data for which valid eye-tracker data was available (527 events in patient 1 and 1242 in patient 2). The figure shows that using the scalp derivative F7-F8 lets us reliably detect eye movements, as is evident from the increase in eye velocity around the epoch of detection, as measured with the video-based eyetracker.

### 4.9 Burst Probability and Rapid Eye Movements

We computed the peri-event histogram between the timing of the thalamic oscillatory bursts in the frequency range of interest and the timing of eye movement events detected in the frontal bipolar channel. For the thalamic bursts, we used all oscillatory bursts detected by our algorithm (described in Section 4.5), with the burst time defined as the time of peak Morlet amplitude. For the eye movement events, we used the time when the first derivative of the voltage in the frontal bipolar channel reached its peak; this would yield approximately the time of the peak eye movement velocity. Next, for each eye movement event, we computed the relative time of all thalamic bursts occurring within ±60 seconds of the eye movement event. We computed such relative timing for all detected eye movement events in REM sleep and binned them in 1 second intervals to obtain the peri-event histograms. We followed a similar procedure to compute the peri-event histogram for the distribution of eye-movement intervals, with the relative timing now being between the given eye-movement event and all other eye-movement events within ±60 seconds.

The correlation between the peri-event histograms of eye movements and burst probabilities was obtained by computing the Pearson correlation coefficient between the two histograms. The significance of the correlation was determined using a Monte Carlo procedure. In each iteration of the procedure, we randomly shuffled the peri-event histogram of the burst probabilities and computed the r-value between the shuffled peri-event histogram of the burst probabilities and the true distribution of eye-movement probabilities. We repeated the procedure 2 × 10^5^ times to obtain a distribution of shuffled r-values. The p-value of significance was determined using a one-sided test, estimated as the fraction of shuffled r-values that were larger than the true r-value.

Figure 3(d) shows the r-values for the 54 thalamic channels with the Wake- and REM sleep-specific oscillations that were detected across the 14 patients. We find a significant correlation (p*<* 9 × 10*^−^*^4^) for the bursts in a total of 50 thalamic channels across 14 patients.

### 4.10 Electrode Localization, Thalamic Segmentation **&** Proximity Analysis

To localize the DBS electrodes for each patient, we used a combination of the workflow within the LeadDBS package^22^ and a Freesurfer implementation of the Thalamic Parcellation routine from^69^. Within the LeadDBS workflow, we first co-registered the pre-operative T1 scan with the post-operative CT scan using advanced normalization tools (ANTs^70^). The native-space coordinates of the four contacts on each of the two DBS electrodes were obtained using the PaCER method^71^ and manually refined if necessary.

The boundaries of thalamic sub-nuclei were determined in native space using a thalamic parcellation routine^69^ implemented in Freesurfer. We also tried out other thalamic atlases based on MNI space, using a further coregistration of the preoperative MRI to the MNI template. However, visual inspection by a neuroanatomy expert found that the native space segmentation^69^ was more reliable than the segmentation based on MNI space atlases. Supplementary Figures 12–46 shows the thalamic segmentation for all 17 patients in our cohort, with the localization of the DBS contacts overlaid.

In order to test our hypothesis that the oscillatory signal is specific to the Central Thalamus, we obtained the mask for the region by grouping together the following two nuclei from the segmentation atlas^69^: MedioDorsal lateral (MDl) and Central Lateral (CL); this approximates previous definitions of the central thalamus [see e.g. refs.^72,73^].

For each of the 140 DBS electrode contacts across our patient cohort (15 patients contributed 8 contacts each, one patient where recording for one of the DBS electrodes was not available contributed 4 contacts, and finally, one patient that was reimplanted contributed a total of 16 contacts), we calculated the proximity of the contact to the thalamic subregion of interest. This was defined as the shortest distance, in native space, from the centre of the contact to the boundary of the thalamic subregion of interest. For centres that were inside or within 1 mm (i.e., less than the resolution of the MRI scans) of a thalamic subregion, we set the distance to an arbitrary value of 0.5 mm (the statistical inference from our regression analysis did not depend on the choice of the arbitrary value, e.g. setting it to 0 mm or 1 mm led to identical statistical inferences). Overall, we found that the centre of 42 contacts was within 1 mm of the Central Thalamus. Further, a total of 47 contacts were within 1 mm of the Anteroventral (AV) nuclei, the main clinical target for DBS.

An important consideration for the analysis of detection probabilities is the fact that all analyses of thalamic field potentials were done on bipolar derivatives of neighbouring channels (e.g. R1-R2); any activity detected on a bipolar derivative could arise due to the proximity of either of the two contacts to the source of the signal (e.g. R1 or R2 for the activity detected on R1-R2). In our analysis, we dealt with this issue in an assumption-free way by assigning the signal to both contacts that went into the bipolar channel. We note that this approach does introduce correlations between neighbouring channels; however, our approach of computing uncertainties and significances using block bootstrapping at the group level ensures that these correlations do not affect the statistical inferences. We also repeated our analyses below using midpoints to each bipolar pair. This led to consistent conclusions as our main analysis(see Extended Data Figure 7).

For each of the two thalamic subregions of interest, we fitted a univariate logistic regression to our measurements, with a binary dependent variable indicating whether the oscillation was detected in the contact (1=detection, 0=non-detection) and the distance to the thalamic subregion serving as the independent variable. The logistic regression was performed using the scikit-learn^74^ package in Python, with an L2 penalty and the ‘liblinear’ solver. Note that for the analysis in Figure 4, we labelled a contact as containing a detection if (i) we detected a wake- and REM specific- oscillation on the contact and (ii) the oscillation significantly correlated with eye movements in REM sleep. However, we verified that dropping the second criterion does not alter the conclusions of the analysis. The uncertainties on the regression coefficients were again computed at the group level using block bootstrapping. We created 10^6^ surrogate cohorts and estimated the logistic regression coefficient for each of the surrogate cohorts to obtain a distribution of regression coefficients. The p-value for rejecting the null hypothesis that the detection probability is uncorrelated was obtained by computing the fraction of surrogate cohorts for which the regression coefficient was greater (lesser) than zero. Finally, the p-value for rejecting the null hypothesis that the slope of the regression for distance to the Central Thalamus is not different from that for the distance to the AV nuclei was obtained by computing the fraction of surrogate cohorts for which the regression coefficient for the AV nuclei was greater than (or lesser than) that for the Central Thalamus.

We note that we also performed a logistic regression to test if the detection probability depended on the distance to three nuclei neighbouring the Central Thalamus: (i) Medio Dorsal medial (MDm), (ii) Ventral Lateral (VL), and (iii) Ventral Anterior (VA). We found no statistically significant evidence for the dependence of the detection probability on the distance to the either the MDm (logistic regression slope= −0.25 ± 0.15, p_corr_ = 0.306, block Bootstrapping) or the VL (logistic regression slope= −0.38 ± 0.21, p_corr_ = 0.148, block Bootstrapping) or the VA (logistic regression slope= 0.18 ± 0.15, p_corr_ = 0.651, block Bootstrapping). All p-values above have been Bonferroni corrected for 3 multiple comparisons.

### 4.11 Aperiodic (1/f) Subtracted Power Spectra

We followed standard procedures to obtain the 1/f subtracted power spectrum for each channel^23^. We first divided our recordings into 2.5-second segments and computed a time-resolved power spectrum using the Welch method in the MNE toolbox. Next, we averaged over these 2.5-second segments to obtain an average power spectrum for each of the three brain states of interest: (i) wakefulness, (ii) REM sleep, and (iii) NREM sleep. Finally, we used the fitting oscillations & one over f (FOOOF) toolbox^23^ to estimate and subtract out the aperiodic component of the power spectrum, separately for each channel and brain state. We used the frequency range from 4 Hz to 48 Hz to fit the aperiodic background, a model without a knee, and limiting the widths of the peaks to the range 2 Hz and 25 Hz, with no constraints on the number of peaks.

For each subject, we obtained an average 1/f-subtracted power spectrum of the thalamic contacts where the Wake- and REM sleep- specific oscillatory bursts were detected, after aligning the spectrum of each contact to the frequency of the peak burst rate detected in the given contact. Figure 2(D, top panel) shows the average of the aligned thalamic power spectra of all 14 subjects where we detect the thalamic activity. The subject-level average was computed after normalizing the 1/f-subtracted power spectrum of each subject by its standard deviation.

Further, to test the specificity of the oscillatory signal to the thalamus, we repeated the above exercise for simultaneously recorded scalp EEG of the same patients. Extended Data Figure 1(B, C, D) shows the subject-level average of the power spectra aligned to the frequency of the oscillatory activity in the thalamic contacts. We grouped the consistently available scalp EEG electrodes in three groups: panel-(b) Frontal scalp electrodes (Fp1, Fp2, F7, F8), panel-(c) Central scalp electrodes (T7, T8, P7, P8), and panel-(d) Occipital scalp electrodes (O1 and O2).

### 4.12 Connectivity Analysis

We used the simultaneously recorded scalp and intracranial EEG to compute the weighted Phase Lag Index (wPLI^75^), a metric of functional connectivity, between oscillations in the thalamic contacts and the scalp EEG channels. We choose to use wPLI because of its robustness to volume conductivity; the wPLI between two oscillatory signals is non-zero only if there exists a consistent positive or negative phase difference between the two signals^75^.

The thalamic field potentials used in this analysis are from the same bipolarly-referenced DBS channels. For EEG signals on the scalp, we used the 10 consistently available EEG electrodes (F7, F8, Fp1, Fp2, P7, P8, T7, T8, O1 and O2), except for one patient where Fp2 was not available.

For each pair of bipolarly referenced intracranial channel and scalp EEG electrode, we first obtained a Morlet transform of both signals, with a wavelet width of 10 cycles, over a frequency range of 8 − 49 Hz (sampling every 1 Hz), and with a temporal sampling of 5 ms (equivalent to our downsampled rate of 200 Hz). Next, we obtained the cross-spectral density between the thalamic contact and the scalp contact, as a function of both time (t) and frequency (f), *X*(*t, f*) = *V*_iEEG_(*t, f*)∗ 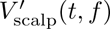, with *V*_iEEG_(*t, f*) referring to the complex-valued time-frequency representation of the signal recorded with intracranial EEG and *V*_scalp_(*t, f*) to the complex conjugate of the time-frequency representation of the signal recorded with scalp EEG. We then obtained, for each of the three brain states, NREM, REM, and wakefulness, the wPLI as a function of frequency by computing the following mean over all epochs in each of the three brain states^75^.

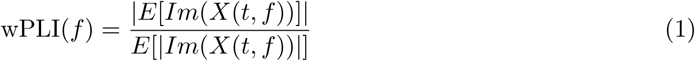

In the above equation, *Im* refers to the imaginary component and *E* refers to the mean across time. The wPLI for the particular Wake- and REM sleep- specific oscillatory band detected in a given thalamic contact was obtained by averaging wPLI(*f*) over the range of frequencies where the oscillatory activity was detected. This procedure thus yielded, for each pair of thalamic channel and scalp EEG channel, three measurements of wPLI in the oscillatory band, one each for NREM sleep, REM sleep, and wakefulness. We average together the wPLI during REM sleep and wakefulness to obtain a single overall estimate of the wPLI for each oscillatory band. We use the measurement of the wPLI in NREM sleep, when the fast oscillatory activity in the thalamus is entirely absent, as a control condition. Finally, given the limited number of scalp EEG electrodes, we group the measurements into the same three broad regions on the scalp: (i) Frontal (F7, F8, Fp1, Fp2), (ii) Central (P7, P8, T7, T8), and (iii) Occipital (O1 and O2).

The average wPLI for the 14 patients in the frequency range where the fast thalamic oscillatory activity was detected is shown in Extended Data Figure 2(a). We detected significantly higher wPLI in REM sleep and wakefulness, compared to NREM sleep, between thalamic contacts and scalp electrodes in the frequency range of the fast thalamic oscillations. For all three effects, p_corr_ *<* 0.006, group-level paired two-sided Wilcoxon Rank Test with 14 subjects, corrected for 3 comparisons using the Bonferroni method. We also repeated the above for the frequency ranges of NREM sleep spindles in the thalamic contact (Extended Data Figure 2(b)). For all three groups of scalp electrodes, we find that the wPLI in frequency ranges of sleep spindles is significantly higher during NREM sleep than during REM sleep and wakefulness (group-level paired two-sided Wilcoxon Rank Test with 14 subjects, corrected for 3 comparisons using the Bonferroni method: statistic(dof=13)=0, p_corr_=0.0004 for Frontal, statistic(dof=13)=2, p_corr_=0.001 for Central, and statistic(dof=13)=2, p_corr_=0.001 for Occipital).

### 4.13 Correlation between Pupil Size and Thalamic Burst Rates

For two of the patients in the current cohort (P15 and P17), where we detected the REM and wake specific fast oscillation, we had eye-tracker data with pupil size tracking available. These data were recorded as part of an unrelated experiment. The eye-tracker data was recorded using Tobii Pro Spectrum at a sampling rate of 600 Hz. For each patient, two sessions were recorded, each lasting ≈ 40 − 50 minutes.

For each of these sessions, we ran our burst detection algorithm on the same bipolarly-referenced channels where we had detected the wake- and REM-specific- fast oscillations. Next, we extracted the timestamps of the bursts that were in the same patient-specific frequency range. We used the timestamps of the detected bursts to obtain a burst rate as a function of time, in 5-second bins. We averaged the burst rates of all thalamic channels where the fast oscillation was detected, based on the analysis of the full wake and sleep data for the patient, to derive an overall thalamic burst rate as a function of time.

The pupil diameter data were extracted and temporally aligned to the thalamic EEG recording. Next, we applied a running-median window of size 100 ms on the pupil diameter measurement of each eye. This was done to reduce noise on the measurement and get a robust estimate of the pupil diameter. The measurements from each eye were then averaged and binned to the same time axis as the burst-rate, i.e., also with 5-second binwidths.

In order to assess the correlation between the pupil diameter and the thalamic burst rate, we performed a standard cross-correlation analysis where the correlation coefficient between the two signals was computed as a function of lag in the range ±300 seconds. We estimate the variance on the cross-correlation thus obtained via a Monte Carlo procedure wherein we repeat the above analysis but after shifting one of the time series by a randomly-drawn lag in the range ±300 seconds. We repeat the procedure 10^4^ times. These shuffled distributions are then used to estimate the confidence interval and p-values of significance.

Extended Data Figure 3 shows, for each of the two patients, the simultaneously recorded pupil size and thalamic burst rates as a function of time. The Figure also shows the correlation coefficient as a function of lag, with the shaded region indicating the 95% ranges of the shuffled distributions. For both patients, we find a significant correlation between the pupil diameter and thalamic burst rate at zero lag (P15: r-value at zero lag = 0.24, *p <* 10*^−^*^4^; P17: r-value at zero lag = 0.57, *p <* 10*^−^*^4^, using shuffled distribution as an estimate of the null distribution).

We note that we also checked for a correlation between the burst rates and saccade rates in the same two patients over the same duration of recording, with eye-tracking data available. However, we found no significant correlation (p*>* 0.05) between the burst rates and saccade rates in the time period that shows a significant correlation between the burst rates and pupil diameter.

### 4.14 Data Availability

The number of patients undergoing DBS for epilepsy at LMU Munich is small, raising the possibility that publicly available raw iEEG data could be used to identify participants who consented to take part in our study. Therefore, the raw data are not shared publicly. Moreover, the thalamic iEEG data used in this work are being used in ongoing projects, which provides an additional reason why the data are not currently being shared. However, subject to applicable data privacy regulations, we will provide the iEEG data to interested researchers for the exclusive purpose of replicating the results reported here. Requests should be addressed to the corresponding author. All source data for generating the figures and the reported statistics are included with the analysis code in the following Github repository: https://github.com/StaudiglLab/code4thalamicbursts.

### 4.15 Code Availability

The code used for all analyses in the manuscript is available at the following GitHub repository: https://github.com/StaudiglLab/code4thalamicbursts

## Acknowledgements

This work was supported by the European Research Council (ERC, ERC-STG Starting Grant 802681, HORIZON-ERC-POC HORIZON ERC Proof of Concept Grants, 101248723, awarded to T. Staudigl) and by the Deutsche Forschungsgemeinschaft (DFG, German Research Foundation) – Project numbers 520617944; 520139811 (awarded to T. Staudigl and E. Kaufmann). We are indebted to all patients who volunteered to participate in this study. E. Kaufmann and T. Koeglsperger were supported by the Medical & Clinician Scientist program (MCSP) scholarship of the LMU Munich. We thank the staff and physicians at the Epilepsy Center, Department of Neurology, Ludwig Maximilians University, Munich, for assistance. A.C. acknowledges discussions with Fabian Schwimmbeck.

## Author Contributions

A.C.: conceptualization, methodology, software, validation, formal analysis, investigation, interpretation of results, data curation, writing — original draft, writing — review and editing, and visualization. X.W.: data collection, writing — review and editing. T.B.: data collection, writing — review and editing. T. Schreiner: writing — review and editing. T.K.: resources and writing — review and editing. J.H.M.: resources and writing — review and editing. J.R.: resources and writing — review and editing. C.V.: resources and writing — review and editing. E.K.: data collection, data curation, methodology, resources, and writing — review and editing. T. Staudigl: conceptualization, methodology, interpretation of results, writing — original draft, writing — review and editing, supervision, project administration, and funding acquisition.

---

**Extended Data Figure 1:**
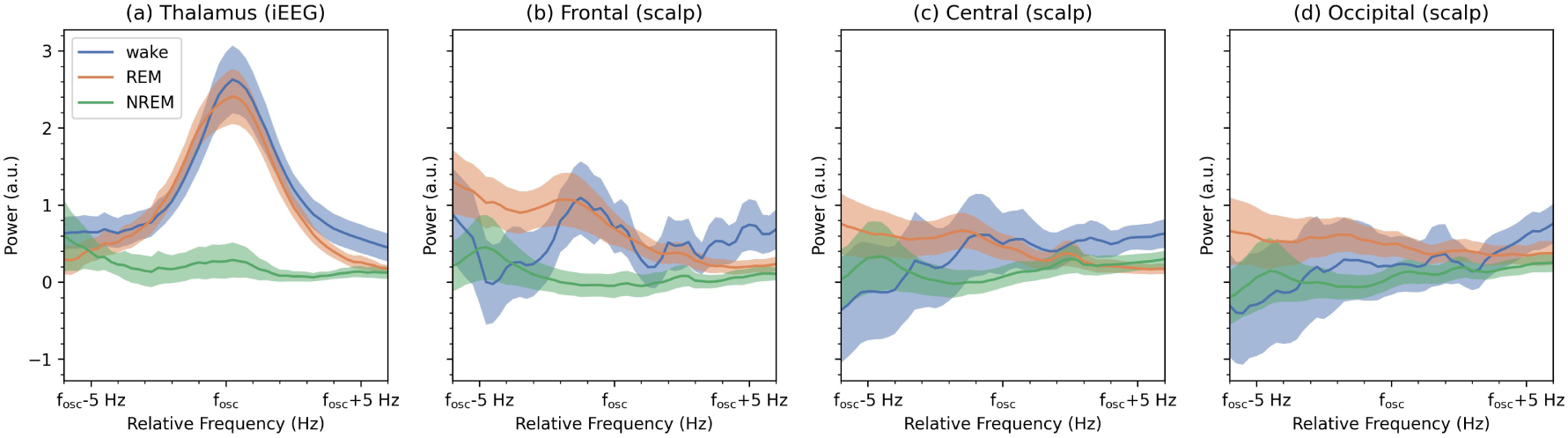
Aperiodic-subtracted Power Spectra for Wake- and REM sleep-specific oscillations (at 19 − 40 Hz). The panels show the power spectra for (a) Thalamic iEEG contacts where the oscillatory bursts were originally detected, (b) Frontal scalp electrodes (Fp1, Fp2, F7, F8), (c) Central scalp electrodes (T7, T8, P7, P8), and (d) Occipital scalp electrodes (O1 and O2). The panels show the subject-level averages of the 1/f subtracted power spectra, separately for (i) wakefulness (blue curve), (ii) REM sleep (orange curve), and (iii) NREM sleep (green curve), with the shaded regions showing the standard error on the mean. The average was obtained by aligning the power spectrum relative to the peak of the frequency band where significant oscillatory activity was detected in the thalamic contacts; the x-axis thus indicates the frequency relative to the frequency of the peak burst rate. The presence of an oscillatory peak during wakefulness and REM sleep is clear on the thalamic contacts, where the oscillations were originally detected. However, no prominent peaks, such as those seen in the thalamic contacts, are seen on the scalp electrodes.

**Extended Data Figure 2:**
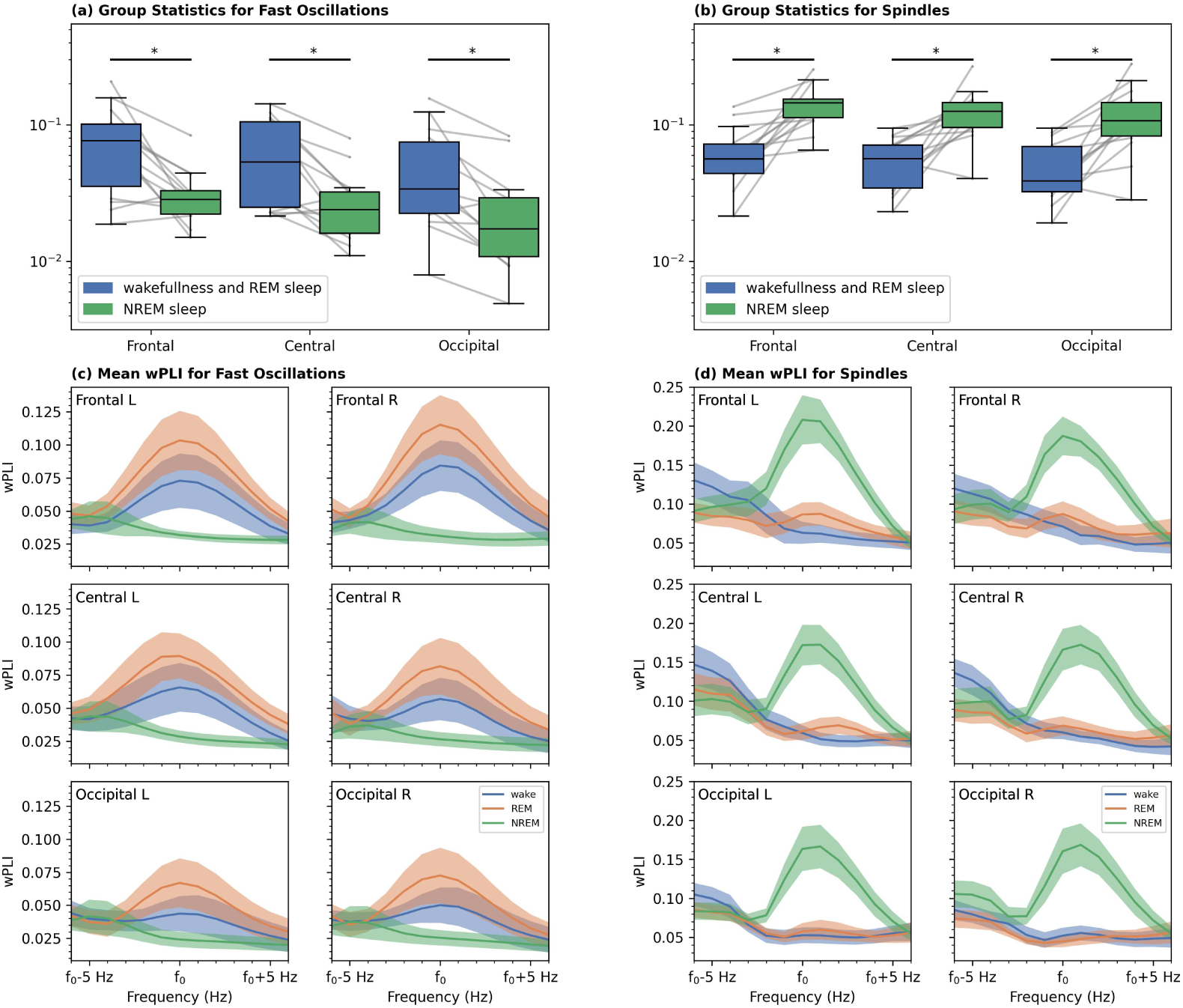
Functional Connectivity between the Thalamic iEEG Contacts and Scalp EEG Electrodes. The figure shows the averaged wPLI (subject-level) between simultaneously recorded thalamic contacts and scalp electrodes in the frequency range of **(a)** the fast thalamic oscillations during REM sleep and wakefulness and **(b)** NREM sleep spindles. Panel **(a)** shows that the wPLI in the frequency range of fast thalamic oscillations for all three regions of scalp electrodes is significantly higher during wakefulness and REM sleep than in NREM sleep. Panel **(b)** shows that the wPLI in the frequency range of sleep spindles is significantly increased for all scalp regions in NREM sleep than in REM sleep and wakefulness. In both panels **(a)** and **(b)**, the boxes extend from the first quartile to the third quartile, with the line indicating the median; the whiskers indicate either the full range of the distribution or 1.5 times the interquartile range, whichever is smaller. Panels **(c)** and **(d)** show, for visualization purposes, the frequency-resolved average wPLI, after aligning to the peak frequency of the thalamic fast oscillations [Panel (c)] and thalamic spindle bands [Panel (d)], separately for each group of scalp electrodes (L: Left Hemisphere, R: Right Hemisphere). The curve in each panel shows the subject-level mean, with the shaded region indicating the standard error on the mean. A clear peak can be seen at the corresponding peak frequencies and for the expected brain states. ^∗^p_corr_ < 0.006, group-level paired two-sided Wilcoxon Rank Test with 14 subjects, corrected for 3 comparisons using the Bonferroni method.

**Extended Data Figure 3:**
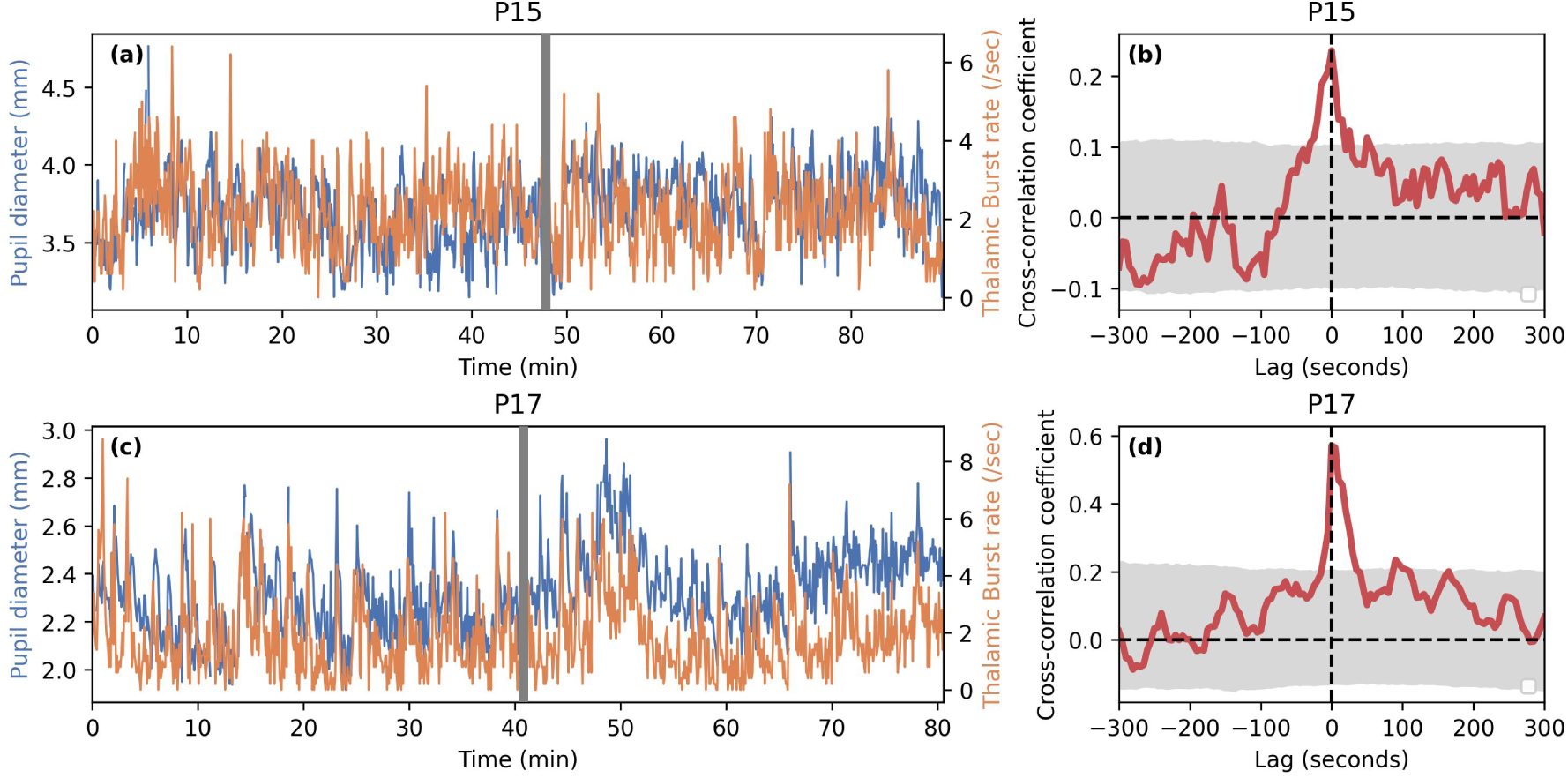
Correlation between pupil diameter and burst rate of the fast thalamic oscillations in two patients. Panels **(a)** and **(c)** show the pupil diameter (in blue) and the thalamic burst rate (in orange) as a function of time. The gray vertical bands show the separation between the two discrete sessions over which the data were acquired (see Methods). Panels **(b)** and **(d)** show the cross-correlation coefficient as a function of lag between the pupil diameter and the thalamic burst rates. The gray shaded regions indicate the 95% confidence interval on the cross-correlation coefficients, obtained via a shuffling procedure (see Methods). For both patients, a significant peak in the cross-correlation can be seen at zero lag (*p <* 10^−4^, one-sided estimate using the shuffled distribution as the null distribution).

**Extended Data Figure 4:**
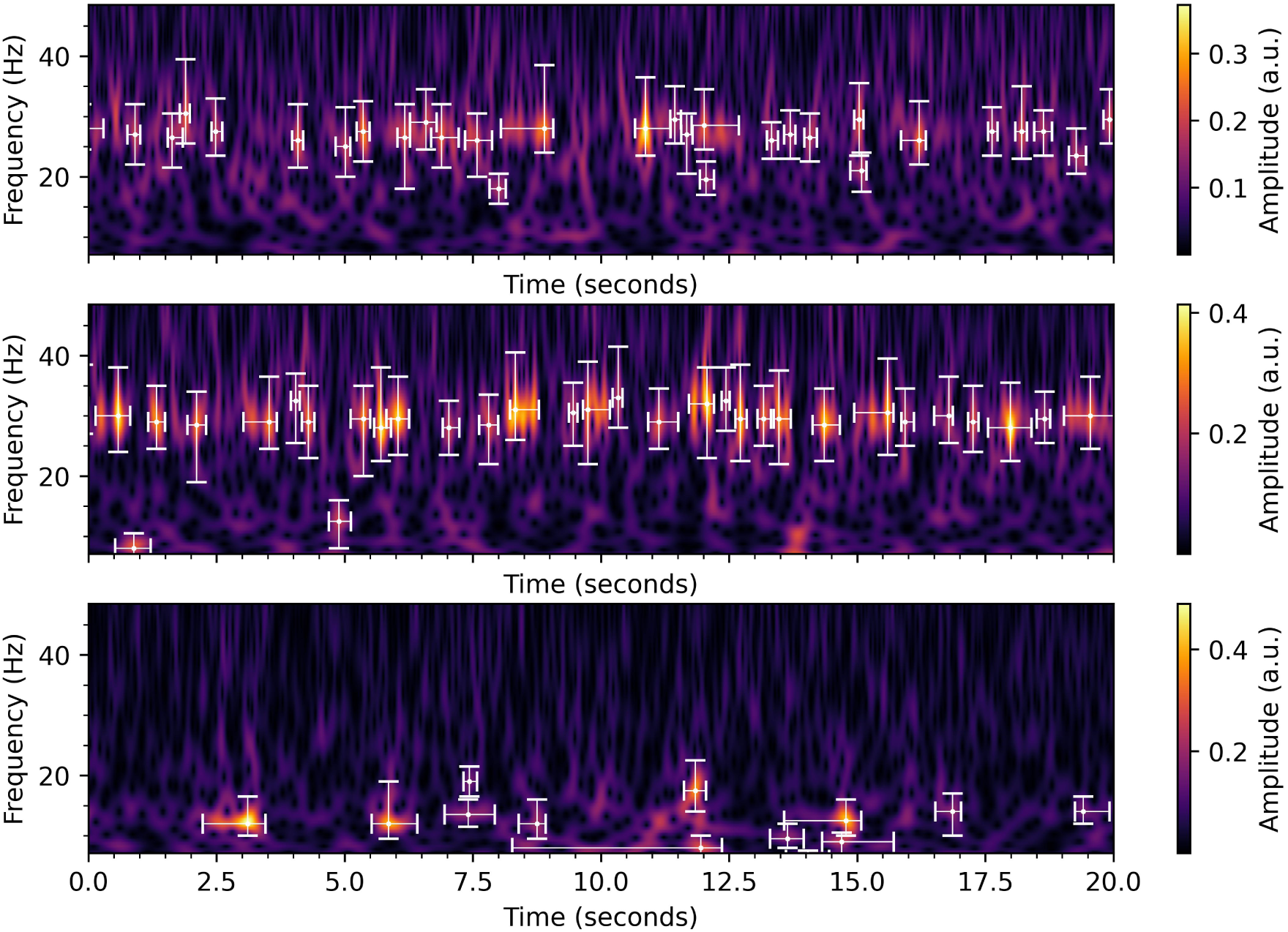
An example illustrating our burst detection algorithm. The panels show 20 seconds of data from an example participant and bipolarly referenced intracranial channel (same as the channel shown in Figure 1(b)), for three different brain states: (top) Wakefulness, (middle) REM sleep, and (bottom) NREM sleep. The Morlet amplitudes as a function of time and frequency are shown in colour with the overlaid white crosshairs indicating the locations of the bursts detected in the data. The extent of the crosshairs along each dimension indicates the extent of the burst that was calculated by our algorithm.

**Extended Data Figure 5:**
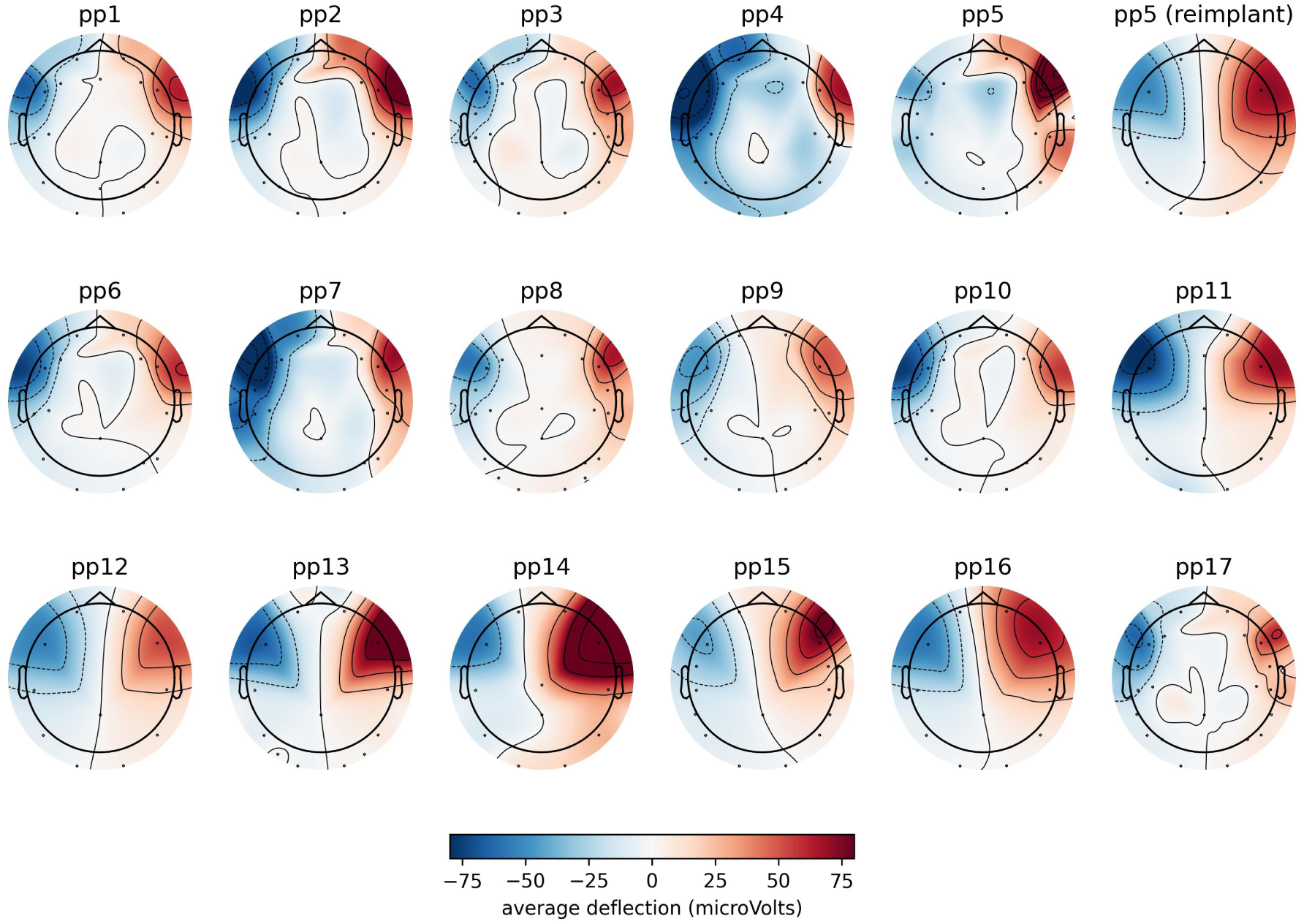
Average scalp topography of voltage deflections during rapid EMs. The topographies are consistent with the stereotypical scalp EEG topography associated with horizontal eye movements. Note that the scalp electrode coverage is variable across our cohort.

**Extended Data Figure 6:**
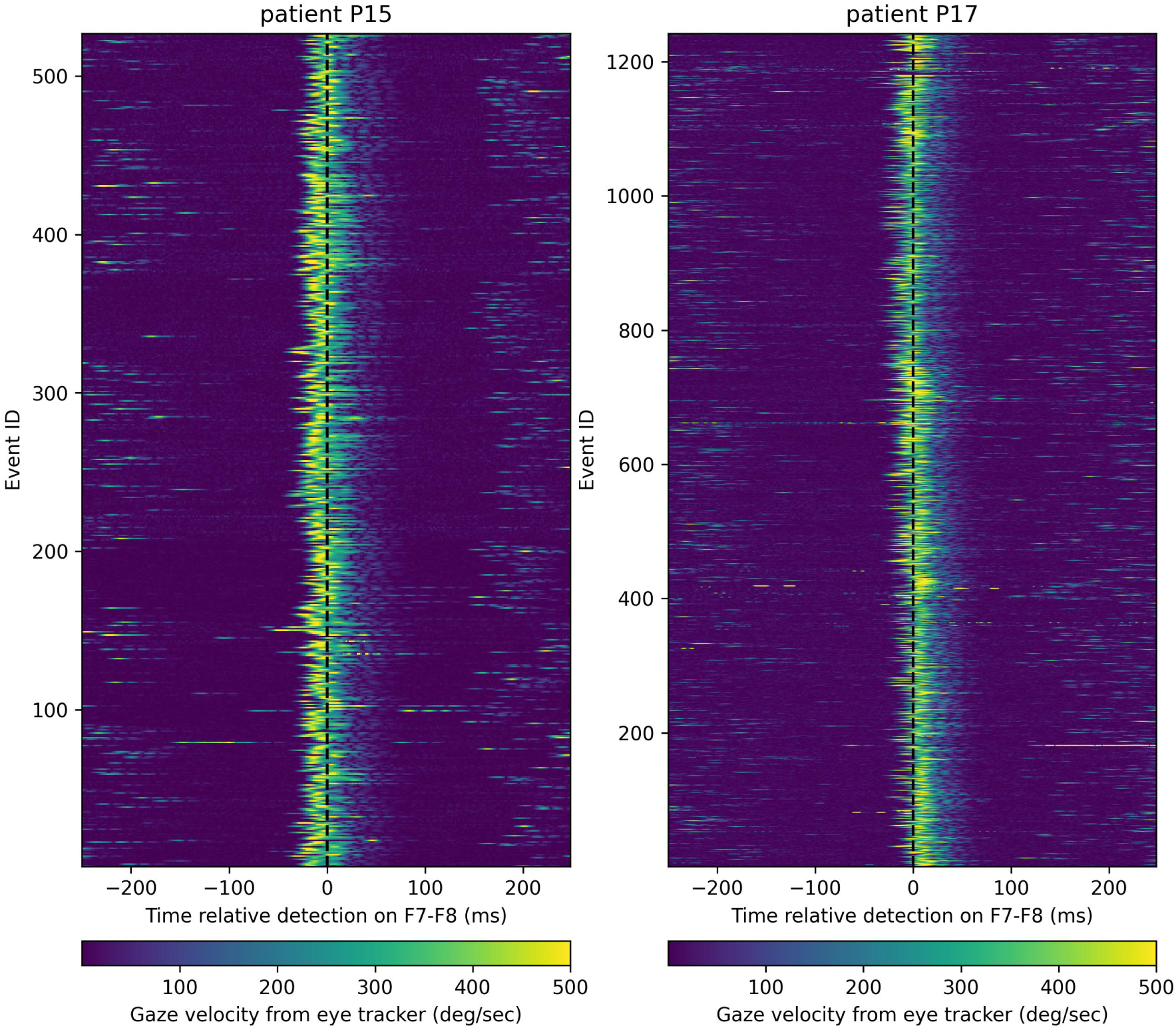
Gaze velocity measurement using a video-based eyetracker locked to the eye-movement events detected in F7-F8. The plot shows a clear enhancement in the gaze velocity, as measured by the video-based eyetracker, around the epoch of events detected on the EEG derivative F7-F8 of the two patients.

**Extended Data Figure 7:**
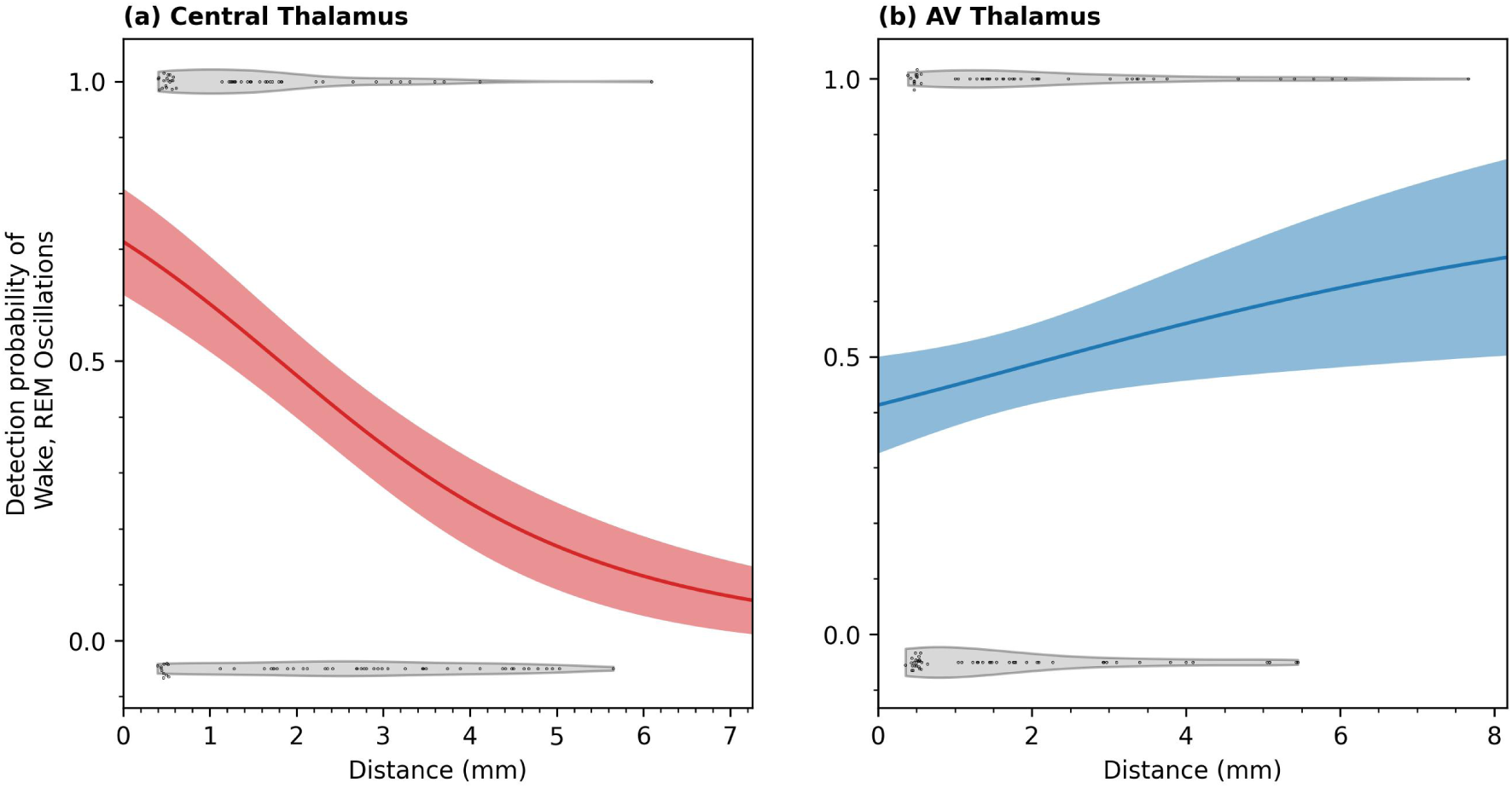
Logistic regression analysis using distance to the midpoint of bipolar contacts. The panels show, for each of (a) Central Thalamus and (b) AV thalamus, the probability of detecting the Wake- and REM sleep- specific oscillations as a function of distance from the thalamic subregions. The solid curves in each plot show the logistic regression curve, with the shaded area indicating the estimated standard deviation. The black dots indicate our measurements; a y-axis value near 1.0 would imply that the Wake- and REM sleep- specific oscillatory signal was detected in the contact, while a y-axis value close to 0.0 would imply no detection. The points clustered around 0.5 mm on the x-axis are those where the midpoint of the pair of electrode contacts was inside the subregion (see Methods); we added a scatter to these points to aid visualization. Note that all DBS contacts for all 17 patients entered the analysis of this figure. The plot shows that the detection probability of the signal is strongly dependent on the distance of the midpoint of the contact to the Central Thalamus (logistic regression slope= −0.51 ± 0.18, p= 7.4 × 10*^−^*^4^, group-level statistic using block Bootstrapping with N=17 patients, see Methods) but not on the distance of the midpoint of the contacts to the AV (logistic regression slope= 0.16 ± 0.14, p= 0.267, block Bootstrapping).

**Extended Data Table 1:** Details on the cohort of patients in the study. Notes: (a) * Patient D was reimplanted due to clinical reasons, (b) For all patients (but one, see note c), the number of bipolarly rereferenced contacts is 6 (3 on each hemisphere), (c) ** For technical reasons, only recording from the right hemisphere was available for Patient M, and hence only 3 bipolar referenced contacts were available, (d) The abbreviations in the table are as follows. SOZ: Seizure Onset Zone, TBI: Traumatic Brain Injury, FLE: Frontal Lobe Epilepsy, FCD: Focal Cortical Dysplasia, FTLE: Frontotemporal Lobe Epilepsy, HS: Hippocampal Sclerosis, MOGHE: Mild malformation of cortical development with oligodendroglial hyperplasia and epilepsy. The seizure onset age for the cohort was 12.3 ± 8.0 yr (mean ± std. dev.).

| Patient | Prior Epilepsy Surgery | SOZ & Etiology | Anti-seizure Medication | Recording Duration (hr) | Number of Thalamic Contacts with detection of fast oscillations | Oscillation on the scalp at similar frequencies (Y/N) |
| --- | --- | --- | --- | --- | --- | --- |
| A | N | multifocal bilateral (dysplasia) | Levetiracetam, Cenobamate | 46.7 | 6 | Y |
| B | Y | multifocal left (unknown) | Lamotrigine, Lacosamide, Clobazam | 32.0 | 1 | N |
| C | N | focal right (double cortex) | Levetiracetam, Lamotrigine | 17.5 | 5 | N |
| D | N | multifocal right (encephalitis) | Levetiracetam, Clobazam, Eslicarbazepine acetate, Phenobarbital | 40.5 | 2 | Y |
| D* | N |  | Phenobarbital, Levetiracetam, Lacosamide, Cenobamate | 40.2 | 6 | N |
| E | Y | Focal right (HS bilateral) | Valproic acid | 112.8 | 3 | N |
| F | Y | FLE bilateral (FCD) | Lamotrigine, Lacosamide, Levetiracetam | 16.7 | 3 | N |
| G | Y | FTLE left (MOGHE) | Brivaracetam, Zonisamide, Lamotrigine, Cenobamate | 102.4 | 3 | N |
| H | N | FLE bilateral (FCD) | Levetiracetam, Lamotrigine, Oxcarbazepin, Zonisamide | 95.6 | - | N |
| I | N | TLE bilateral (HS bilateral) | Lamotrigine, Lacosamide, Zonisamide | 61.6 | 2 | N |
| J | N | Focal left (TBI) | Levetiracetam, Phenytoin, Lamotrigine | 19.7 | 2 | N |
| K | N | multifocal bilateral (FCD) | Levetiracetam, Lacosamide, Cenobamate, Perampanel | 83.5 | - | N |
| L | N | multifocal bilateral (encephalitis) | Levetiracetam, Lamotrigine | 18.5 | 1 | Y |
| M** | Y | TLE bilateral (unknown) | Topiramate | 39.2 | - | N |
| N | N | FLE left (gliosis/encephalitis) | Lacosamide, Oxcarbazepine, Topiramate | 11.2 | 4 | N |
| O | N | multifocal bilateral (FCD, microcephaly) | Lacosamide, Clobazam, Levetiracetam, Phenytoin | 20.1 | 6 | Y |
| P | Y | TLE bilateral (unknown) | Cenobamat, Lacosamide | 47.1 | 4 | N |
| Q | N | multifocal bilateral (unknown) | Brivaracetam, Phenytoin, Clobazam, Eslicarbazepine acetate | 17.3 | 6 | N |

