## Supplementary Information for "Thalamic oscillations distinguish natural states of consciousness in humans"

#### Contents

**1 Granger Causality Analysis** **2**

**2 Correlation between Amplitudes of Spindles and Fast Oscillations** **2**

**3 Thalamic Burst Rates and Pathology** **5**

**4 Rate of Thalamic Fast Oscillations and Spindles in N2 sleep and N3 sleep** **7**

**5 Rate of IEDs across brain states** **8**

**6 Infraslow Oscillations in Thalamic Burst Rates during REM sleep** **8**

**7 Thalamic Bursts and Saccades in Wake** **9**

**8 Electrode Localization and Anatomical Specificity** **9**

### 1 Granger Causality Analysis

To assess the presence of any dominant directionality in the connectivity between the thalamic contacts and the scalp EEG contacts (See Extended Data Figure 2), we perform a Granger Causality analysis. We use the MNE connectivity package[1] to compute frequency-resolved Granger Causality (GC) scores between all pairs of bipolarly-referenced thalamic iEEG contacts and scalp EEG contacts, in 10-second time segments. The package uses a state-space formalism to compute spectral GC[2]. For each pair of channels and time segment, we computed both the GC score for a drive of the scalp EEG channel by the thalamic iEEG channel ($GC_{\mathrm{Thalamus->Scalp}}$) and vice versa ($GC_{\mathrm{Scalp->Thalamus}}$). We compute a net GC score for each pair of channels, defined as:

$$GC_{\mathrm{net}} = GC_{\mathrm{Thalamus->Scalp}} - GC_{\mathrm{Scalp->Thalamus}} \quad (1)$$

A positive $GC_{\mathrm{net}}$ would thus be indicative of a dominant thalamus-to-cortex drive.

The average net GC score for the 14 patients in the frequency range where the fast thalamic oscillatory activity was detected, for the same three groups of scalp EEG electrodes as in Extended Data Figure 2, is shown in Supplementary Figure 1(a). We detected a significantly higher net GC score in REM sleep and wakefulness, compared to NREM sleep, between thalamic contacts and scalp electrodes in the frequency range of the fast thalamic oscillations. For all three effects, $p_{\mathrm{corr}} < 0.004$, group-level paired two-sided Wilcoxon Rank Test with 14 subjects, corrected for 3 comparisons using the Bonferroni method (Frontal scalp electrodes: statistic(dof=13)=5, $p_{corr}$=0.0036; Central scalp electrodes: statistic(dof=13)=0, $p_{corr}$=0.00036; Occipital scalp electrodes: statistic(dof=13)=5, $p_{corr}$=0.0036). The results of this analysis are thus consistent with the thalamus driving the cortex at these frequencies.

A common issue with analysis of GC scores is the presence of correlated noise between pairs of channels that can bias GC scores, leading to false positives[3]. An analysis technique that is sometimes used to obtain robustness against such biases is to do a correction to the net GC score that is based on a time-reversal operation on each of the two time series involved[4]. We again used the MNE connectivity package to compute the time-reversed GC scores, also in both directions, to obtain the following time-reversal corrected net GC score:

$$GCTR_{\mathrm{net}} = (GC_{\mathrm{Thalamus->Scalp}} - GC_{\mathrm{Scalp->Thalamus}}) - (GC^{TR}_{\mathrm{Thalamus->Scalp}} - GC^{TR}_{\mathrm{Scalp->Thalamus}})$$

where $GC^{TR}_{\mathrm{Thalamus->Scalp}}$ and $GC^{TR}_{\mathrm{Scalp->Thalamus}}$ are the GC scores obtained after flipping the time-axis, for a thalamic iEEG to scalp EEG and scalp EEG to thalamic iEEG drive, respectively.

Supplementary Figure 1(b) shows the same as in panel (a) of the Figure, but now for the GC scores with the time-reversal correction ($GCTR_{\mathrm{net}}$).

The evidence of directionality that the standard GC scores show (Panel a and c of Supplementary Figure 1(a)) is no longer statistically significant in the time-reversal corrected scores (Frontal scalp electrodes: statistic(dof=13)=24, $p_{corr}$=0.235; Central scalp electrodes: statistic(dof=13)=20, $p_{corr}$=0.126; Occipital scalp electrodes: statistic(dof=13)=28, $p_{corr}$=0.406, group-level paired two-sided Wilcoxon Rank Test with 14 subjects). It is unclear why the time-reversal correction has this effect on the GC scores; potential causes could be (a) the much lower signal-to-noise ratio of the scalp EEG compared to the thalamic iEEG, leading to a difficulty in computing GC scores for scalp-to-thalamus connectivity, and (b) the presence of bidirectional connectivity between the scalp and thalamic channels which makes the interpretation of the time-reversal correction non-trivial[5]. A clearer understanding of the connectivity would potentially require simultaneous thalamic and cortical iEEG, providing high signal-to-noise ratio recordings in both the thalamus and cortex. Moreover, a conclusive understanding of the drive for the signal may only be possible with further investigation of the signal in animal models.

### 2 Correlation between Amplitudes of Spindles and Fast Oscillations

Given our observations that the thalamic field potential switches between sleep spindles during NREM sleep and faster oscillations during wakefulness and REM sleep, we considered the possibility that the oscillations might have similar amplitudes. To test this, we computed, for each thalamic contact, the average burst profile for the sleep spindles as well as the Wake- and REM sleep- specific oscillations (50 contacts across 14 patients where the thalamic oscillations correlated with rapid EMs). We measured the average amplitude of oscillatory bursts from the average burst profile of

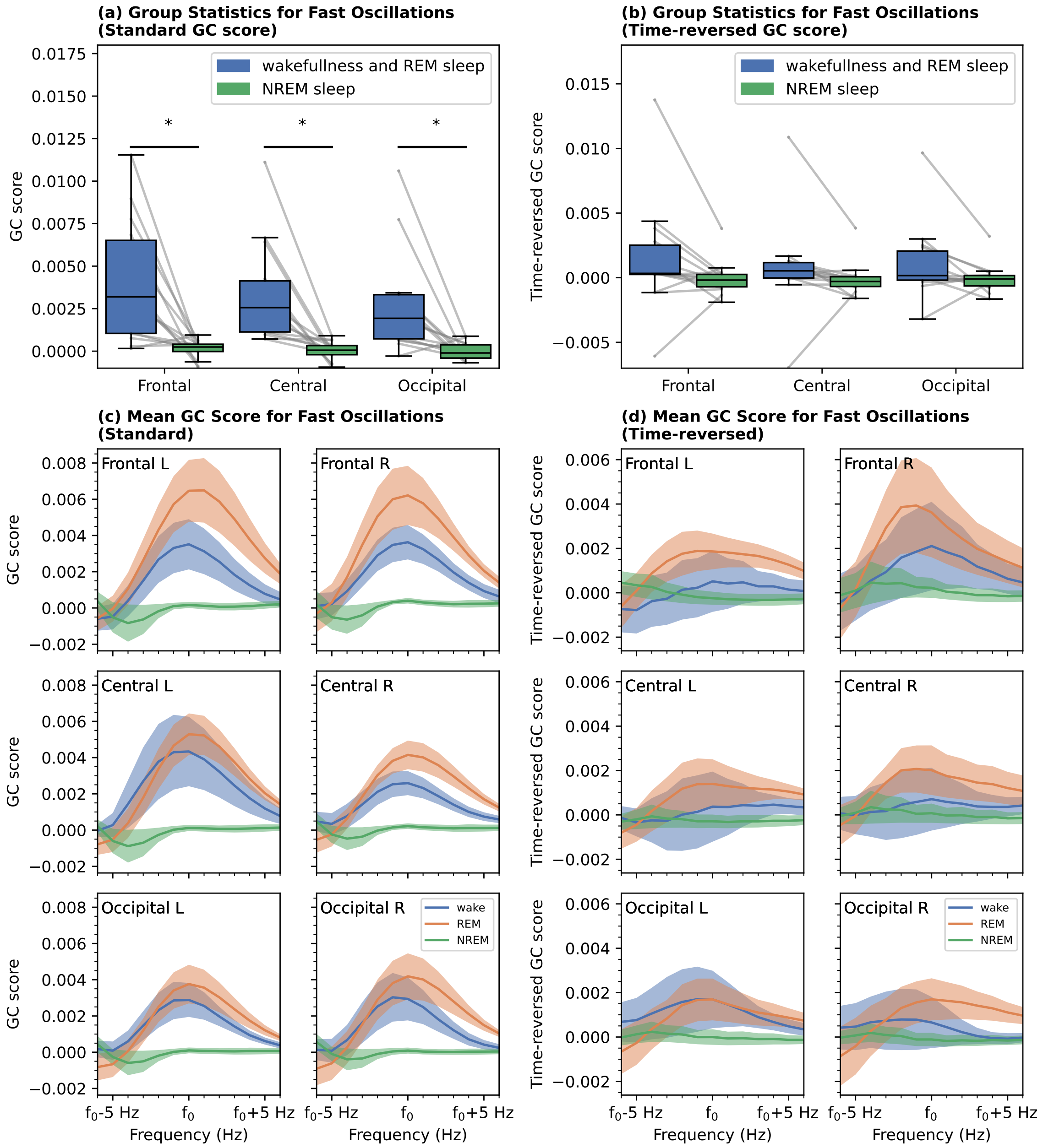


Supplementary Figure 1: **Granger Causality between the Thalamic iEEG Contacts and Scalp EEG Electrodes.** The figure shows the averaged Granger causality scores (subject-level) between simultaneously recorded thalamic contacts and scalp electrodes in the frequency range of the fast thalamic oscillations during REM sleep and wakefulness. Panel **(a)** shows the net Granger Causality score in the frequency range of fast thalamic oscillations for all three regions of scalp electrodes. Panel **(b)** shows the same, but with the scores now having a time-reversal correction[4], $GCTR_{net}$. Panels (c) and (d) show, for visualization purposes, the frequency-resolved average wPLI, after aligning to the peak frequency of the thalamic fast oscillations separately for each group of scalp electrodes (L: Left Hemisphere, R: Right Hemisphere), for the standard GC score and the time-reversed GC scores, respectively. While a clear peak can be seen at the corresponding peak frequencies for the standard GC score, implying a thalamus-to-cortex drive, this is not the case after applying the time-reversal correction. See text for a discussion. (* $p_{corr} < 0.004$, group-level paired two-sided Wilcoxon Rank Test with 14 subjects, corrected for 3 comparisons using the Bonferroni method). In both panels **(a)** and **(b)**, the boxes extend from the first quartile to the third quartile, with the line indicating the median; the whiskers indicate either the full range of the distribution or 1.5 times the interquartile range, whichever is smaller.

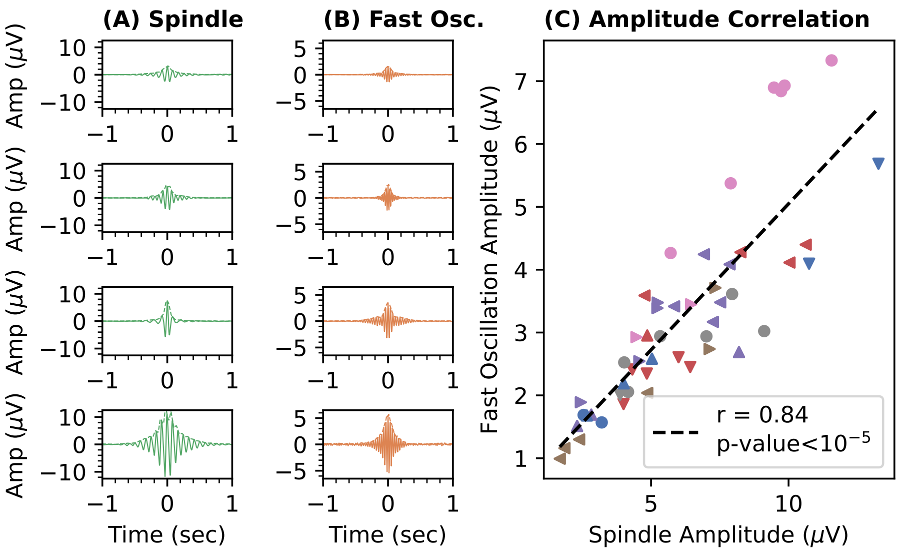


Supplementary Figure 2: **Correlation between average burst amplitudes of sleep spindles in NREM sleep and faster oscillations in REM sleep and wakefulness.** The four rows in Panels **(a)** and **(b)** show, for four different patients, the average sleep spindle profile during NREM sleep and the average profile of fast oscillation during wakefulness and REM sleep, detected in the same thalamic contact. The dashed line in each panel shows the envelope of the burst, obtained via a Hilbert transform. Panel **(c)** shows the amplitude of the average fast oscillation as a function of the amplitude of NREM sleep spindles in the same thalamic contact. The amplitudes of the oscillatory activity in the different brain states are significantly correlated ($p < 10^{-5}$, block bootstrap with N=14 patients). In panel (c), each unique combination of colors and symbols represents measurements from a patient.

the detected bursts in a given frequency range. This was done by taking the phase-coherent average of individual bursts via the following steps. First, for each detected burst in a frequency range of interest, we obtained the bandpass filtered voltage series and determined the time at which the filtered voltage reached its maximum value around ±1 cycle of the peak amplitude detected in the Morlet transformed space (as determined using the detection algorithm, see prior Sections on burst detection). This effectively allowed us to identify the time-point of the burst corresponding to the zero-crossing of the phase. Next, we extracted ±4 sec of voltage (without any filtering) around the identified time-point of each detected burst. We averaged the aligned voltage series across all bursts to obtain an average burst profile for each oscillatory band in each thalamic contact. Any slow modulation in the average burst profile was removed by filtering the time series above 8 Hz. Figure 2(a) and (b) show examples of average burst profiles for four different patients in our cohort. We note that similar results were obtained for various choices of filtering, including filtering the voltages to the individual frequency ranges of detection. The amplitude of the burst was obtained by measuring the peak voltage of the average burst.

Figure 2(c) shows, for the entire cohort, the amplitude of the Wake- and REM sleep- specific fast oscillations as a function of the amplitude of NREM sleep spindles. We find significant correlations for the amplitudes (Pearson $r = 0.833 \pm 0.040$, $p < 10^{-5}$, Block Bootstrap at the group level with 14 subjects) of the average sleep spindles and faster oscillations.

An important consideration for the amplitude correlation is the effect of electrode impedances. We have now computed the amplitude correlation after regressing out the effect of the impedance of each contact. In this analysis, we used measurements of bipolar impedances on each pair of contacts that were made at the time of activating the patient's stimulators; this was typically done 3 weeks after the implantation. These measurements were available for all but one patient in our cohort. We used a partial correlation analysis to regress out the effect of impedance, treating it as a covariate. Supplementary Figure 3(a) shows the amplitudes of the two oscillations after subtracting out the effect of impedances on both the amplitudes of NREM sleep spindles and of the fast oscillations (the individual regressions are shown in panels (b) and (c) of Supplementary Figure 3). We find a statistically significant correlation between the amplitudes of the two oscillations even after regressing out the effect of impedances ($r = 0.828 \pm 0.042$; $p < 10^{-5}$, Block bootstrapping with N=13 patients). The analysis shows that impedance variability across contacts and electrodes is not the primary driver of the high correlation between the amplitudes of the two oscillations ($r = 0.833 \pm 0.040$, see Supplementary Figure 3). We note that DBS contact impedances are known to drift slowly over time (see e.g., ref.[6]); however, this is not expected to affect the conclusions of this analysis due to the short gap of only approximately 3 weeks between our iEEG recordings and the impedance measurements.

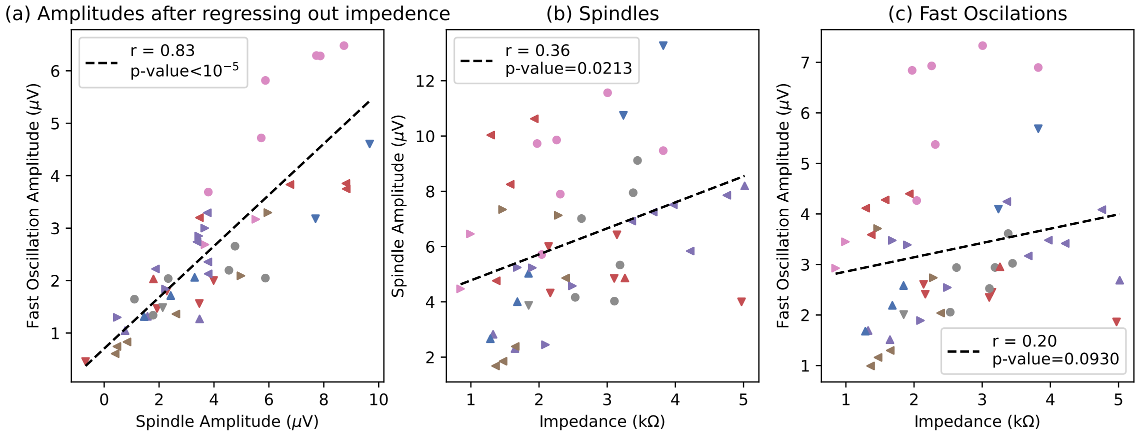


Supplementary Figure 3: **Effect of contact impedances on the correlation between the average burst amplitudes of NREM sleep spindles and fast oscillations in REM sleep and wakefulness.** Panel (a) shows, after regressing out the effect of impedances, the amplitudes of the two oscillations on each bipolar contact. The regression lines used to subtract out the effect of impedances are shown in panels (b) and (c). Note that the negative amplitudes in panel (a) are an effect of subtracting out the regression lines of panels (b) and (c) from the individual amplitude values. We find that the amplitudes of the two oscillations are significantly correlated even after regressing out the effect of contact impedances ($p < 10^{-5}$). All statistics were done at the group-level with N=13, using block bootstrapping. In both panels, each unique combination of colors and symbols represents measurements from a patient.

We also carried out an additional analysis to further understand the issue. A high electrode impedance of an EEG contact leads to high noise levels on the channel[7]. If electrode impedances drive the correlation, this is likely to also cause a related anti-correlation between the amplitudes and the rates of the bursts. For example, an electrode contact with a relatively good impedance should lead to a better detection threshold (in microVolts). The detections of a larger number of weaker events would lead to a higher event rate and a lower average event amplitude. An impedance-based explanation of the amplitude correlation should thus lead to an anti-correlation between mean amplitudes and event rates. We thus tested whether the spindle rates and the average spindle amplitudes are correlated. We found no statistically significant correlation between these two measures (Pearson-r=$0.19 \pm 0.14$, p=0.083, Block Bootstrapping at the group level). We also repeated this for fast oscillations to find a significant correlation instead of an anti-correlation between these measures (Pearson-r=$0.64 \pm 0.20$, p=0.009, Block Bootstrapping at the group level). The results of this analysis further show that the amplitude correlation of Supplementary Figure 2 is unlikely to arise purely due to an impedance effect.

### 3 Thalamic Burst Rates and Pathology

We tested whether the rate of thalamic bursts, both fast oscillations during Wake and REM sleep, as well as spindles during NREM sleep, correlates with the seizure frequency. For this, we used a baseline total seizure frequency measurement one month before the DBS implantation surgery, and hence, the date of recording of the data presented in this work. We found no significant correlation between the baseline seizure rate and the rate of thalamic fast oscillations, nor between the seizure rate and the rate of spindles in the same thalamic contacts (Figure R3 below, panels a and b, for rates of the wake- and REM- specific oscillations Spearman-r=0.01, p=0.98; for rates of NREM sleep spindes, Spearman-r=0.20,p=0.50).

We also investigated any differences in the rates of either fast oscillations or spindles based on epilepsy types. We note that the epilepsy types for the patients in our cohort are very diverse, with different focuses and etiologies (Extended Data Table 1). It is thus unlikely that the presence of the oscillation itself is due to a network pathology. Further, after consulting with an epileptologist (E.K.), we classified the cohort of patients using different criteria: (i) unilateral epilepsies vs. bilateral epilepsies and (ii) Fronto-Temporal Lobe Epilepsies (FTLE) or other types. We then tested whether either the rate of fast oscillations in Wake and REM sleep or the rate of spindles in NREM sleep is different between these different cohorts. For both classifications, we did not find any significant differences in the rates (Supplementary Figure 4 below, panels c-f, Mann-Whitney U test, p> 0.11 in all four cases; Bilateral vs. Unilateral for fast oscillations: statistic(dof=13)=35, p=0.111, Bilateral vs. Unilateral for spindles: statistic(dof=13)=15, p=0.364, FTLE vs. not FTLE for fast oscillations: statistic(dof=13)=22, p=0.805, FTLE vs. not FTLE for spindles: statistic(dof=13)=18, p=0.458).

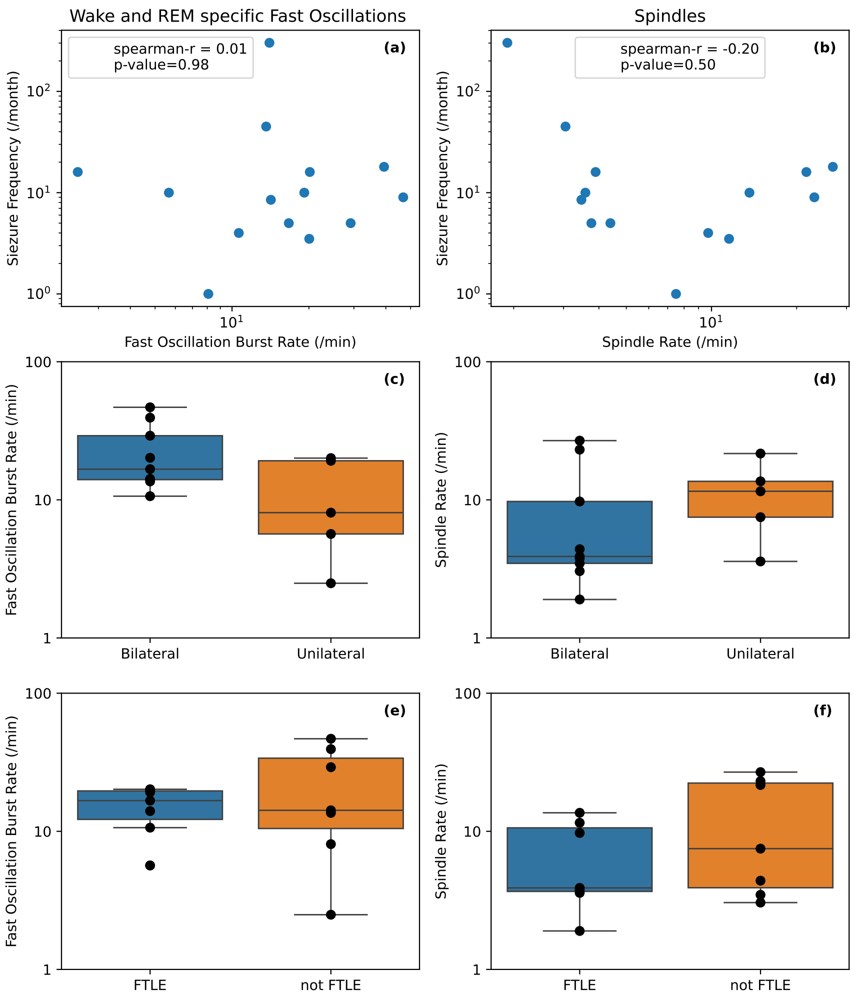


Supplementary Figure 4: **Relationship between fast oscillation (spindle) rates in the thalamic contacts and pathological properties.** Panels (a) and (b) show the baseline seizure frequency, measured one month before the DBS implantation surgery, as a function of the rate of the fast oscillation in Wake and REM sleep (Panel a) and that of the rate of spindles in NREM sleep (Panel b), for the 14 patients where we detected the thalamic fast oscillation. The data in both panels are consistent with there being no correlations between either the rate of fast oscillations or that of spindles and the baseline seizure rate (Spearman-r=0.01, p> 0.98 for fast oscillations and Spearman-r=-0.22, p> 0.45 for spindles). Panels (c) and (d) show the rates of fast oscillations and spindles, respectively, for patients diagnosed with unilateral epilepsies and bilateral epilepsies. Panels (e) and (f) show the same, but now dividing the cohort based on whether they had Fronto-Temporal Lobe Epilepsies (FTLE) or not. We did not find any significant differences between the two cohorts for either of the two classifications, for both fast oscillation rates and spindles (Mann-Whitney U test, p> 0.11 in all four cases). Note that the rates of both fast oscillations and spindles were consistently computed from thalamic iEEG contacts where the fast oscillation was detected. For all boxplots, the boxes extend from the first quartile to the third quartile, with the line indicating the median; the whiskers indicate either the full range of the distribution or 1.5 times the interquartile range, whichever is smaller.

### 4 Rate of Thalamic Fast Oscillations and Spindles in N2 sleep and N3 sleep

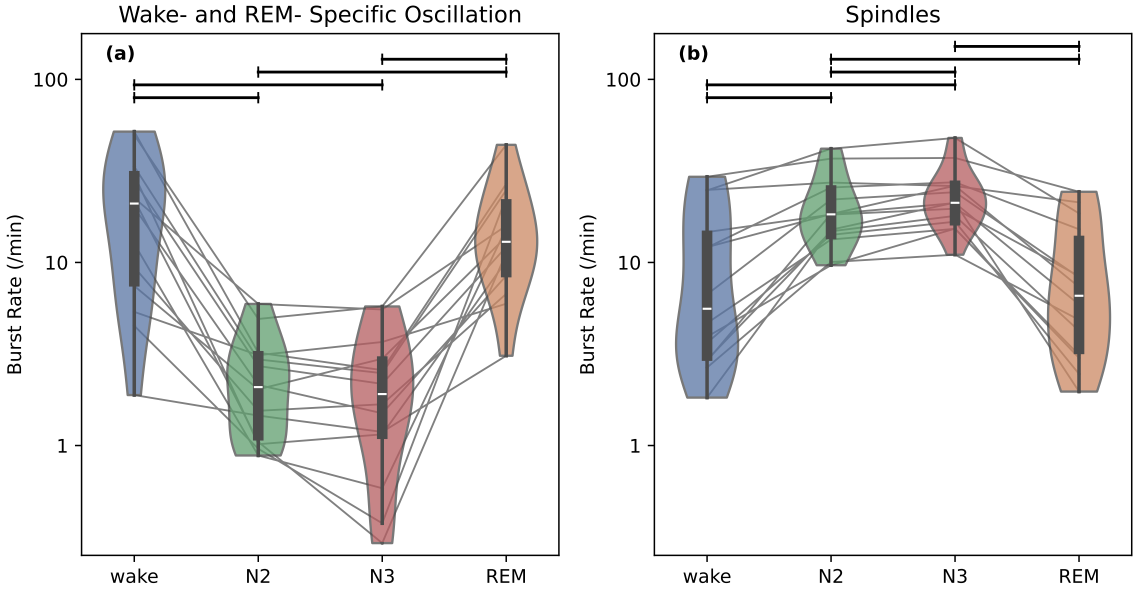


Supplementary Figure 5: **The rate of Wake- and REM- specific Oscillations and NREM Spindles in Thalamic contacts, as a function of sleep stages, with NREM sleep divided into N2 and N3.** Significant differences in burst rates (two-sided Wilcoxon Rank Test at the group level, N=14, $p_{\mathrm{corr}} < 0.05$, after correcting for multiple comparisons) between pairs of brain states are indicated with horizontal lines on the top. In both panels, the boxes extend from the first quartile to the third quartile, with the line indicating the median; the whiskers indicate either the full range of the distribution or 1.5 times the interquartile range, whichever is smaller. For the Wake- and REM- sleep specific fast oscillations, shown in Panel (a), the burst rate is significantly higher in REM and wake compared to both N2 and N3, with no differences in the rates between N2 and N3. For the NREM sleep spindles, shown in Panel (b), the rates in N2 and N3 are significantly higher than in both wake and REM sleep. Further, the rate of spindles in the thalamic contacts is significantly higher in N3 than in N2.

Supplementary Figure 5 shows the burst rates of both oscillations as a function of sleep states, with NREM divided into N2 and N3. For both oscillations, the figure shows the trends expected from our main findings: (i) the burst rate of the thalamic fast oscillations in each of wake and REM is significantly higher than in both N2 and N3 (two-sided Wilcoxon Rank Test at the group level, N=14, statistic (dof=13)=0, $p_{corr}$=0.0007 for all four comparisions, Bonferroni corrected for a total of 6 pair-wise comparisons) and (ii) the burst rate of spindles is significantly higher in each of N2 and N3 than in both wake and REM (two-sided Wilcoxon Rank Test at the group level, N=14, statistic=0, $p_{corr}$=0.0007 for all four comparisons, Bonferroni corrected for 6 pair-wise comparisons). Further, for the fast oscillations, we find no statistically-significant difference in the rates between N2 and N3 (two-sided Wilcoxon Rank Test at the group level, N=14, statistic=25, $p_{corr} = 0.543$).

Interestingly, for sleep spindles, we find that the rate in N3 is significantly higher than in N2 (two-sided Wilcoxon Rank Test at the group level, N=14, statistic=2, $p_c orr = 0.002$). While there is some consensus that spindle rates are higher in N2 than in N3 in the human cortex[8], the same for the rates in the thalamus is unclear. In the few studies that use direct iEEG recordings from the Thalamus[9–11], the rate of sleep spindles between N2 and N3 is either consistent or the rate is found to be significantly higher in N3 than in N2 (similar to our findings here). We note that these papers also studied different regions of the thalamus (e.g., the Pulvinar in ref.[9], ANT and MD in ref.[12]). The prior studies, along with our findings here, are indicative that the relative rates of spindle in N2 and N3 are different in different parts of the brain (and potentially between different parts of the thalamus).

#### 5 Rate of IEDs across brain states

Supplementary Figure 6 shows the IED rates as a function of brain states. We did group-level paired two-sided Wilcoxon Rank Tests on all possible pairs to find that the IED rate is significantly lower in REM sleep than in Wake (two-sided Wilcoxon Rank Test at the group level, N=14, Wake vs. REM: statistic (dof=13)=1, $p_{corr}$=0.0007; NREM vs. REM: statistic=49, $p_{corr}$=1; NREM vs. wake: statistic (dof=13)=21, $p_{corr}$=0.15; Bonferroni corrected for a total of 3 pair-wise comparisons). While the lower IED rate in REM sleep is broadly consistent with findings in the literature[13], we note that the difference in brain regions and epilepsy types could contribute differently to IED rates.

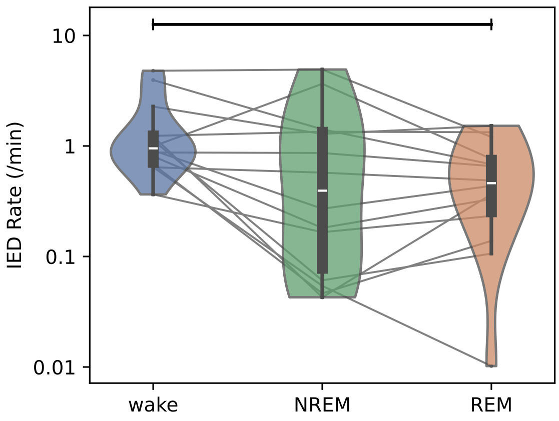


Supplementary Figure 6: **IED rates as a function of brain states.** The boxes extend from the first quartile to the third quartile, with the line indicating the median; the whiskers indicate either the full range of the distribution or 1.5 times the interquartile range, whichever is smaller. We find that the rate of IEDs in REM sleep is significantly less than in wake ($p_{corr} < 0.0007$, group-level paired two-sided Wilcoxon Rank Test with 14 subjects, corrected for the three comparisons).

#### 6 Infraslow Oscillations in Thalamic Burst Rates during REM sleep

In order to check for the presence of an infraslow periodicity in the REM- and wake- specific thalamic bursts, we used the time stamps of the bursts detected by our burst-detection algorithm to get a burst rate as a function of time, in time bins of 0.5 seconds. Next, for each patient, we selected REM sleep segments that lasted for more than 5 minutes. We computed the power spectrum for each of these segments using the Welch method (with a FFT length of 512 samples), in the frequency range 0.008-0.5 Hz. For each patient, we combined all REM segments across all thalamic channels where we had detected the oscillation to get one burst-rate power spectrum per patient. We next fitted a 1/f aperiodic component to this power spectrum, with a spectral knee, and subtracted it out to get a single power spectrum per patient. Supplementary shows the average 1/f-subtracted power spectrum of the fast thalamic bursts in REM sleep (blue curve). We did a one-sided permutation cluster t-test to check for significantly enhanced oscillatory power to find a significant cluster in the range 0.043-0.054 Hz (p=0.036). Given our findings that the thalamic bursts during REM sleep are tightly correlated with epochs of rapid EMs, we also repeated the above procedure to compute a 1/f-subtracted power spectrum for the rate of rapid EMs (with identical choice of bin widths and parameters for the computation of the power spectrum). Supplementary Figure 7 also shows the power spectrum for the rapid EMs (orange curve). We again performed a one-sided permutation cluster t-test to find significantly enhanced power over the frequency range 0.031-0.055 Hz (p=0.0004). Thus, consistent with our findings that the thalamic bursts during REM sleep are tightly correlated with the bursts of rapid eye movements, we find an overlapping range of frequencies over which the power spectrum of both the thalamic burst rate and rate of rapid EMs is higher than what is expected from a purely aperiodic power spectrum. The frequency ranges where we find the significant enhancement correspond to time scales of 20-30 seconds, shorter than the minute time scale oscillations of rapid EMS in REM sleep that have been reported in the literature[14]. This could be due to a difference in the subject cohort for the two studies (five young and healthy subjects in Ktonas et al.[14] vs. 14 epilepsy patients with a large variability in age in this study).

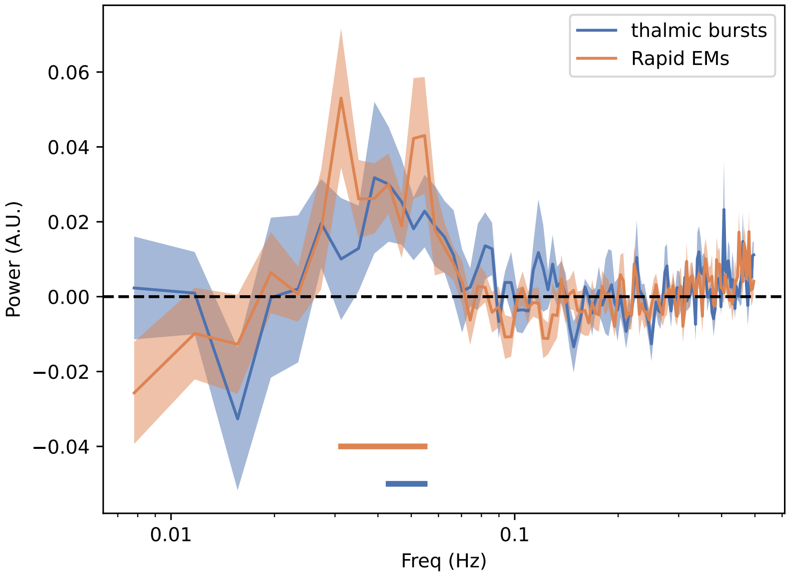


Supplementary Figure 7: **Infraslow oscillatory component (1/f-subtracted) of the power spectrum of the rate of thalamic bursts in REM sleep (blue curve) and the rate of rapid eye movements (orange curve).** The curves show the subject-level means, while the shaded regions indicate the standard errors on the mean. The horizontal bars indicate the frequency ranges with significant enhancements in oscillatory power (one-sided permutation cluster t-test, $p < 0.05$). Both the rate of thalamic bursts and of rapid EMs show significant oscillatory power in an overlapping frequency range.

### 7 Thalamic Bursts and Saccades in Wake

We note that repeating our analysis with wake eye movements comes with multiple challenges. Unlike REM sleep, where eye movements occur in bursts with stereotyped durations, wakefulness can have very different occurrence rates of eye movements depending on the alertness levels and the activity being performed by the patient at the clinic. Further, there would be confounds between periods of open eye wakefulness and closed-eye wakefulness, which would be difficult to disentangle in the absence of video-based eyetracking.

Supplementary Figure 8(a) shows the peri-event histograms of the burst rates relative to time of wake-eye movements as well as the eye movements themselves, for the same thalamic contacts where we detected a correlation between thalamic bursts and rapid EMs in REM sleep. The figure shows that while the peri-event histograms of the thalamic bursts during wakefulness are correlated with the peri-event histograms of the eye movements themselves ($r = 0.84 \pm 0.02$), the enhancement in the burst-rates around the eye movements are both temporally less specific and much weaker compared REM sleep (see Supplemetary Figure 8), peak modulation of $6 \pm 1\%$ over baseline in wake vs. $66 \pm 9\%$ over baseline in REM). There are at least two distinct possibilities for this difference (i) the thalamic activity is directly related to occulomotor activity but the continuous nature of eye movements in wakefulness leads to an elevated baseline level and hence a smaller modulation of the burst rates, and (ii) the thalamic bursts correlate with global states of the brain (such as arousal levels in wakefulness and phasic vs. tonic states in REM) rather than individual eye-movements.

In this context, it is interesting to note that Extended Data Figure 3 shows that the burst rate of the fast thalamic oscillations in wakefulness correlates significantly with pupil diameter (in two patients), a widely used correlate of arousal level, but not with the saccade rate (see Materials and Methods). The data from the two patients thus tentatively support the hypothesis that the central thalamic wake- and REM- specific oscillations are tightly coupled with levels of arousal in wakefulness.

### 8 Electrode Localization and Anatomical Specificity

The segmentation of the human thalamus from T1-weighted MRI images is quite challenging, with a large number of segmentation algorithms that are available, each of which is based on a different

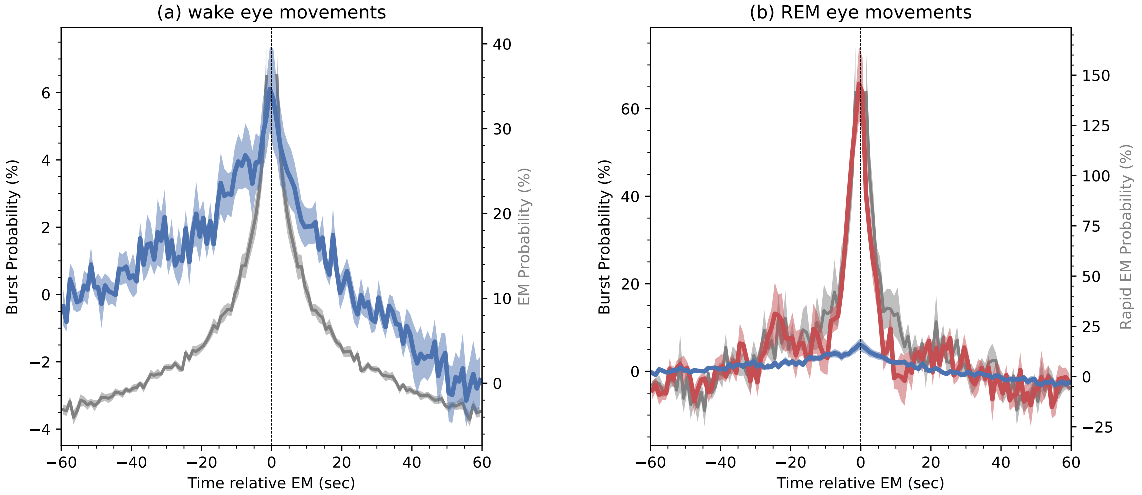


Supplementary Figure 8: **Thalamic Bursts and Eye Movements in wake**. Panel (a) shows, in blue, the average peri-event histogram of occurrence probability of thalamic bursts (in %) relative to the time of EMs, averaged over all 14 patients that showed the wake- and REM- specific fast oscillations; the shaded band indicates the standard error on the mean. The grey curves indicate the probability distribution of other EMs relative to the current EM. Panel (b) shows the same, but for EMs in REM sleep (in red), with the peri-event histogram of bursts during wake also shown (in blue). Note that the curves for REM sleep in Panel (b) are the same as in Figure 3(E). The enhancement of the burst probability at the time of EM, relative to baseline activity, is found to be much higher in REM sleep than in wake.

approach and ground-truth database (e.g., ref.[15] and references therein). There are only a few anatomical landmarks that are typically available in MRI images to clearly delineate the boundaries between different thalamic nuclei, implying the exact boundaries of the nuclei in individual patients may not be entirely accurate (see, for example, ref.[16], for a discussion on the difficulty of targeting the ANT due to the high individual variability in the location and size of the nucleus). However, the algorithms are indeed designed to ensure that the segmentation boundaries are, *on average*, correct.

We had chosen to use for this work a native-space algorithm[17]. In our testing, we found native space segmentations to be far superior to an MNI-space atlas (such as those typically used with the standard LeadDBS workflow[18]), as it entirely avoids the native-to-MNI coregistration process, an additional step that adds to the localization error. Note that patient brains are prone to asymmetries that are particularly problematic when converting from native to MNI space. While this makes the localization more accurate, it makes it impossible to accurately visualize all contacts on a single MRI. Supplementary Figure 12–46 shows the localization for each of the 136 thalamic contacts on the native-space MRI of the 17 individual patients.

However, for the ease of visualization, we have also created an overview plot by performing the additional step of native-to-MNI space coregistration. We emphasize again that the results thus obtained do not reflect the native-space localization. For example, we show the MNI-space coordinates of the contacts that were localized in the native space to the Anterior Thalamus and the Central Thalamus in Supplementary Figure 9, with the MNI space atlas overlaid on it. The figure clearly shows that the localization in the MNI space inaccurately depicts the native-space localization in the native space of the patients. With this declaimer stated, we also used a similar approach to help visualize the location of all DBS contacts that were used in this work, with color coding to indicate the contacts where we detected the fast thalamic oscillation versus those where we did not do so (see Supplementary Figure 10.

The other critical issue in understanding anatomical specificity comes from the fact that we are recording with macroelectrodes (a standard for most iEEG papers that do not have microwire recordings for single units). The size of each of the macroelectrode contacts was 1.5 mm in length and 1.27 mm in diameter, implying that in some cases the contact itself could be at the boundary of two nuclei and thus pick up signals from both. This issue is further complicated by the use of bipolar rereferencing, which, while necessary to derive spatially confined mesoscale field potentials, could imply that each of the contacts lies in two different thalamic nuclei. Bipolar rereferencing is widely used in human iEEG studies and is also widely believed to be the best approach to get a handle on local activity. However, it is likely that, when using a macroelectrode channel, the probability of detecting a neural signal from a given nucleus of origin remains non-zero outside the exact boundary of the nucleus, with the probability dropping with the distance of the contact to the nucleus.

The above issues, in both the thalamic segmentation itself and the specificity of macroelectrode recordings, informed our current analysis methodology, where we pooled all the contacts together to analyze if the detection probability of the wake- and REM- specific oscillations varied with distance from a particular nucleus. Such an approach takes into account the nature of the data we worked with, making the outcome robust against the above uncertainties in the specificity of an individual contact.

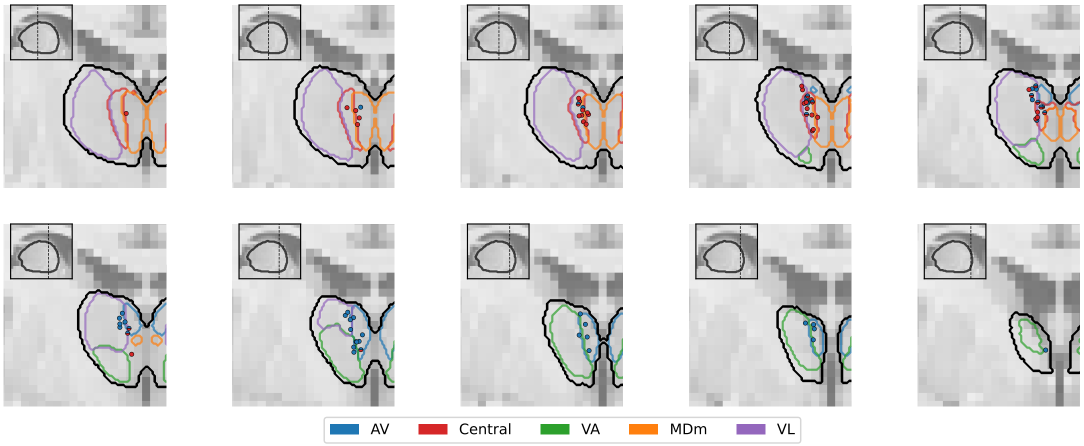


Supplementary Figure 9: **Thalamic Segmentation in MNI Space.** The figure shows different coronal slices on an MNI brain, with the plane of the slices shown in the top-left insets. The contours show the MNI space boundaries of the different nuclei in the Iglesias et al.[17] atlas. The MNI-coordinates of the electrode contacts of all 17 patients in our cohort, which were classified in the native patient space as being in the AV, are shown with blue-filled circles and those in the Central Thalamus with red-filled circles. Note that we transformed all coordinates to the left hemisphere for a clearer visualization. The contacts shown with half red and blue fillings are those that lie within 1 mm of both the AV and the Central Thalamus. Note that 1 mm is similar to the length of the DBS contacts as well as the typical resolution of MRI. The figure illustrates that transforming patient MRIs to the MNI space introduces additional errors that lead to inaccuracies in localization. See Supplementary Figure 12–46 for the individual native-space localizations of all contacts for the 17 patients in our cohort.

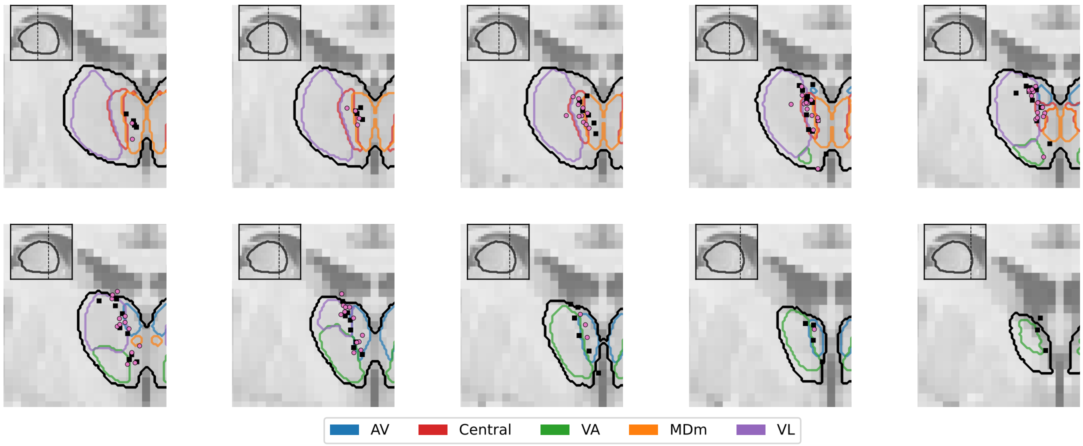


Supplementary Figure 10: **Visualization of contact localizations in the MNI Space.** The same as in Supplementary Figure 9, but now with the MNI coordinates of all 132 contacts from the 17 patients being plotted. Note that this figure is for visualization purposes only and that the plot does not accurately reflect the native space localizations (See Supplementary Figure 9). The magenta points indicate the contacts where the Wake and REM- Sleep specific oscillation was detected. Note that the oscillation was detected with bipolar montages, and we thus indicated both contacts that were part of a bipolar montage with a magenta circle. The black squares indicate contacts where we did not detect the oscillation. See Supplementary Figures 12–46 for an accurate visualization of the individual native-space localizations of all contacts for the 17 patients in our cohort.

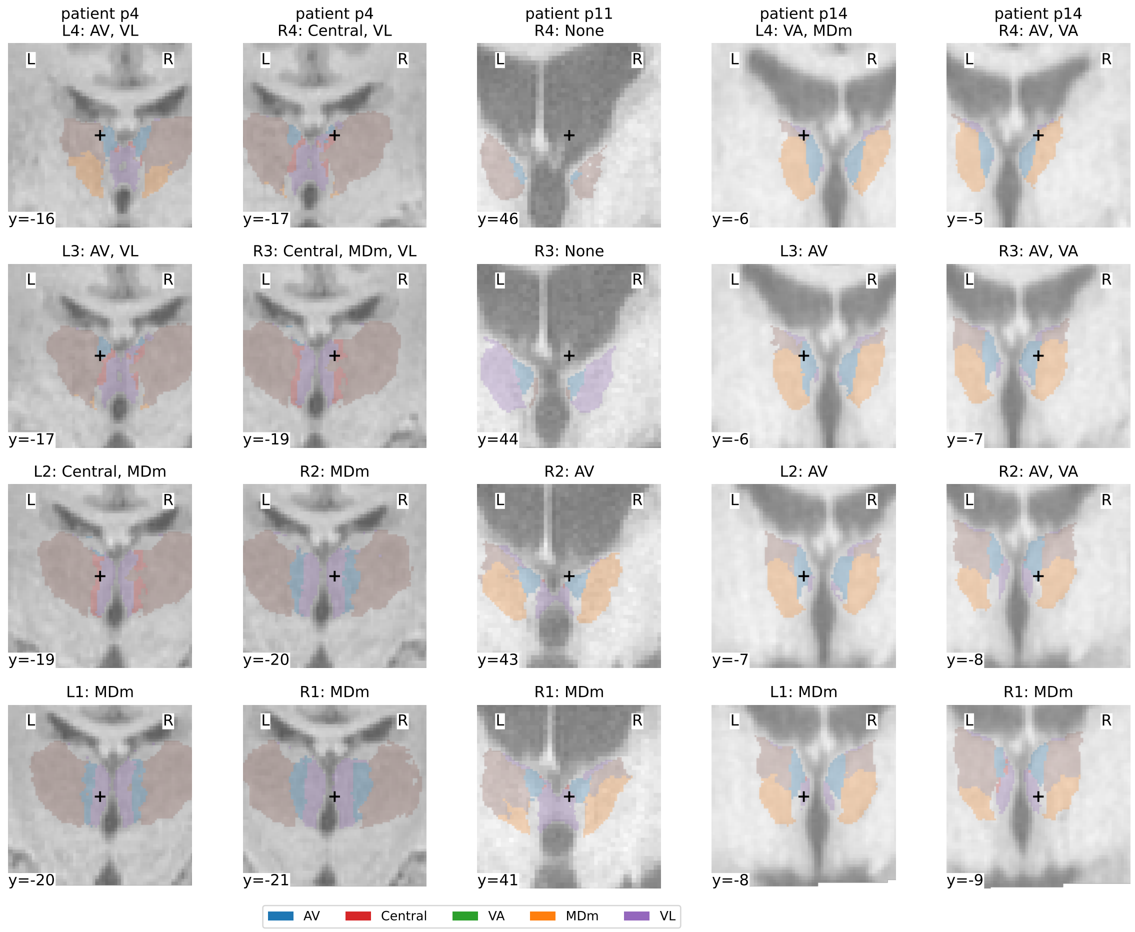


Supplementary Figure 11: **Localization of contacts for patients for whom we did not detect the wake and REM- sleep- specific thalamic oscillations.** Each row shows a particular DBS contact in a particular patient. Note that for p11, we did not have access to the recording from the left hemisphere and have thus shown here only the relevant right electrode. The black cross indicates the location of the contact on the sagittal slice of the native-space patient T1-weighted MRI (for a view of all slices, see the Supplementary Figures 18, 19, 34, 39, 40). The filled contours indicate the segmentation obtained from the Freesurfer algorithm[17]. The labels on top of each slice also indicate the thalamic region(s) that the contact was localized to; note that we have taken into account here the 1 mm length of the contacts.

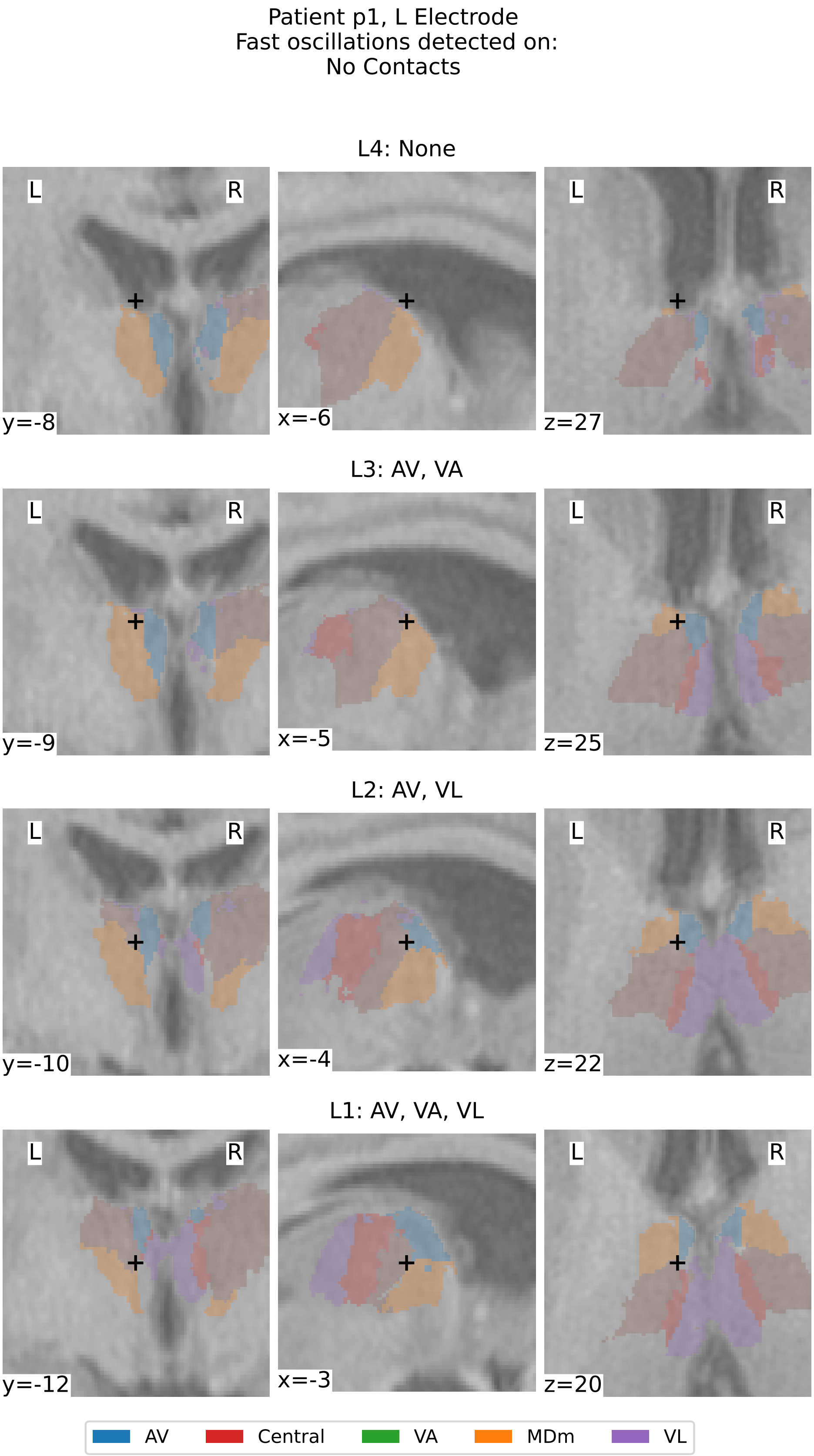


Supplementary Figure 12: **Thalamic Segmentation and Localization of contacts for patient p1, left electrode.** Each row shows a particular DBS contact in the patient. The black cross indicates the location of the contact on slices of the patient's native-space T1-weighted MRI. The filled contours indicate the segmentation obtained from the Freesurfer algorithm[17]. The labels on top of each slice also indicate the thalamic region(s) that the contact was localized to; note that we have taken into account here the 1 mm length of the contacts. AV: Antroventral, VA: Ventral Anterior, MDm: MedioDorsal medial, VL: Ventral Lateral.

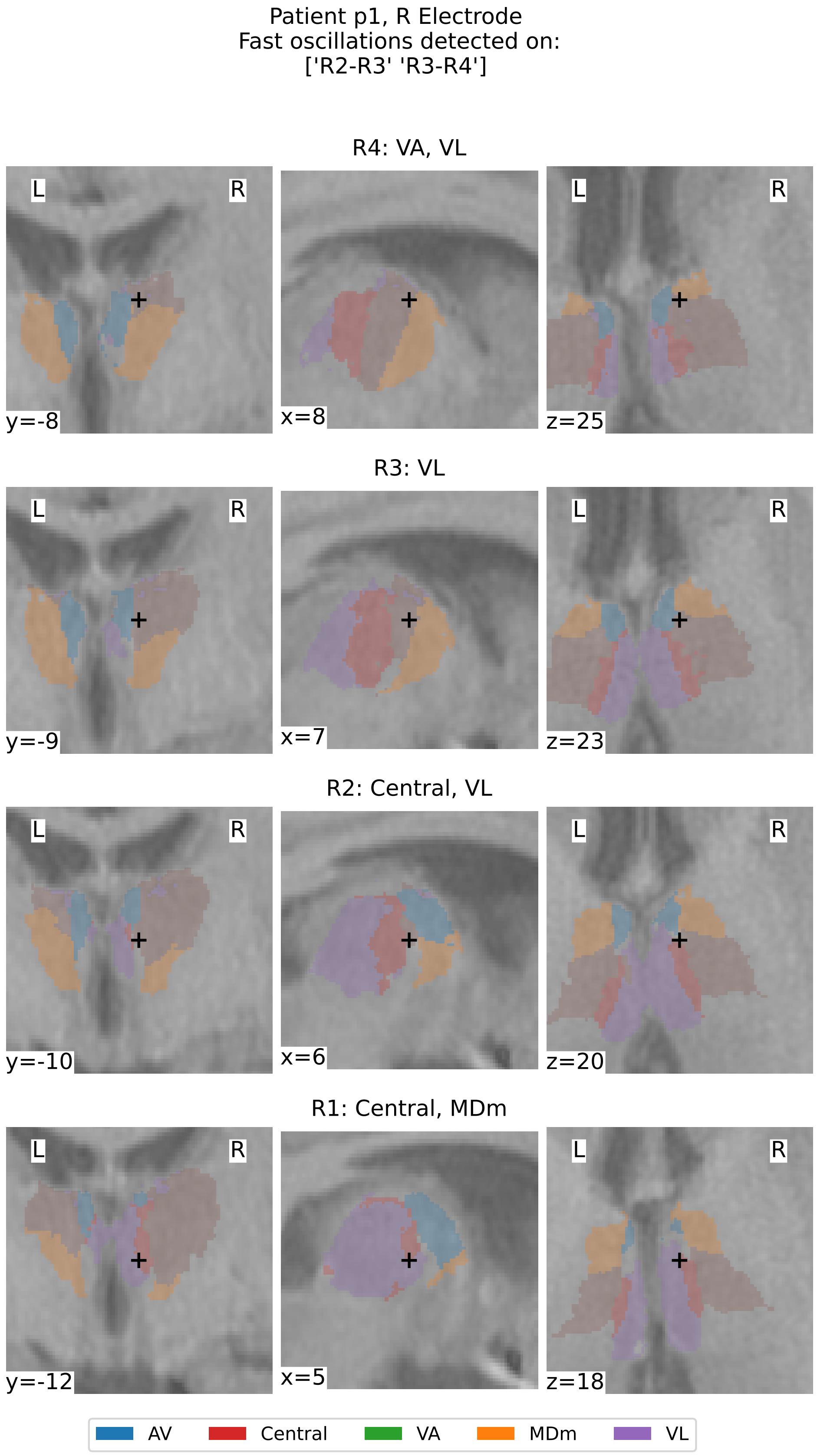


Supplementary Figure 13: **Thalamic Segmentation and Localization of contacts for patient p1, right electrode.** See caption of Supplementary Figure 12.

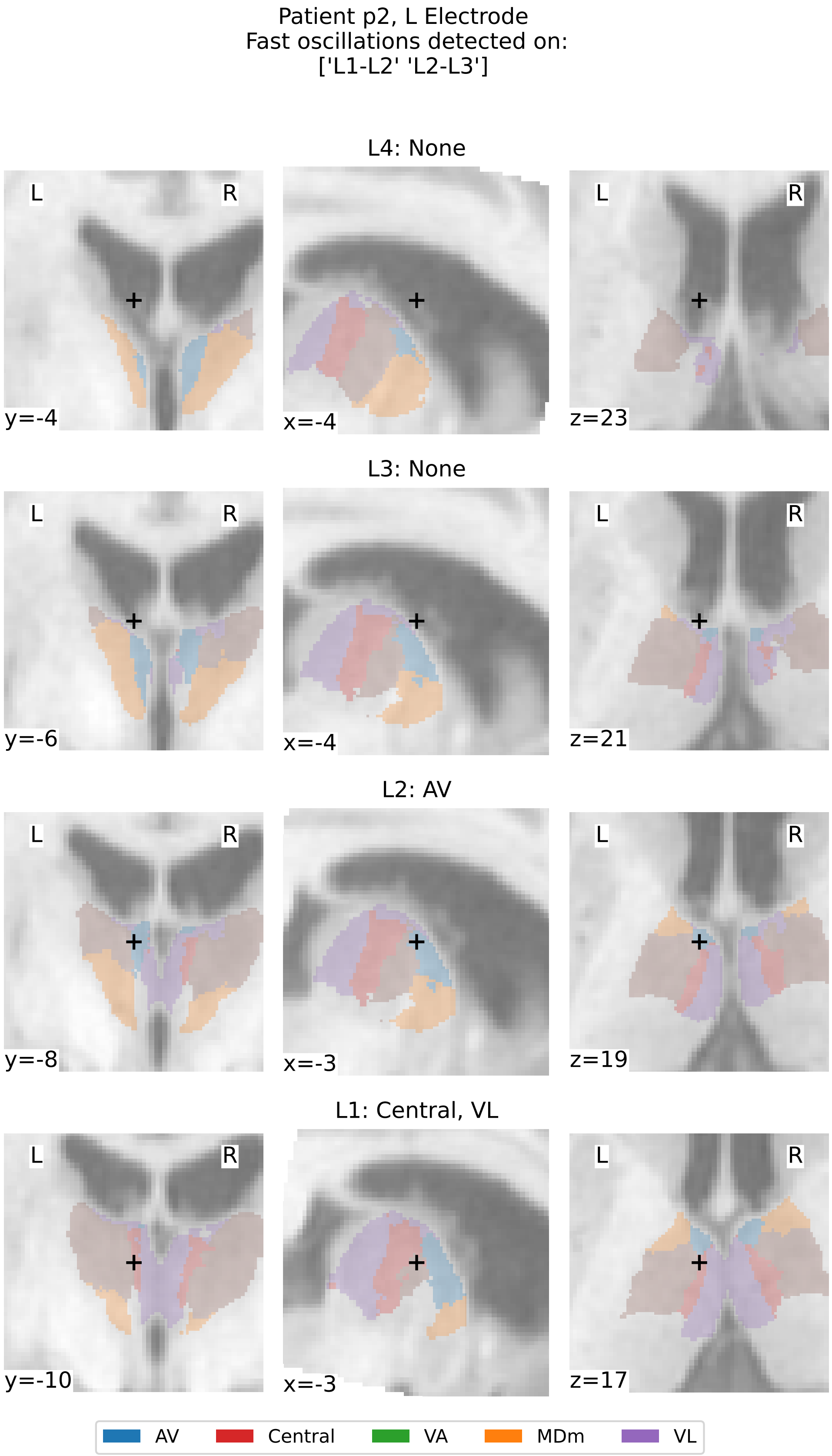


Supplementary Figure 14: **Thalamic Segmentation and Localization of contacts for patient p2, left electrode.** See caption of Supplementary Figure 12.

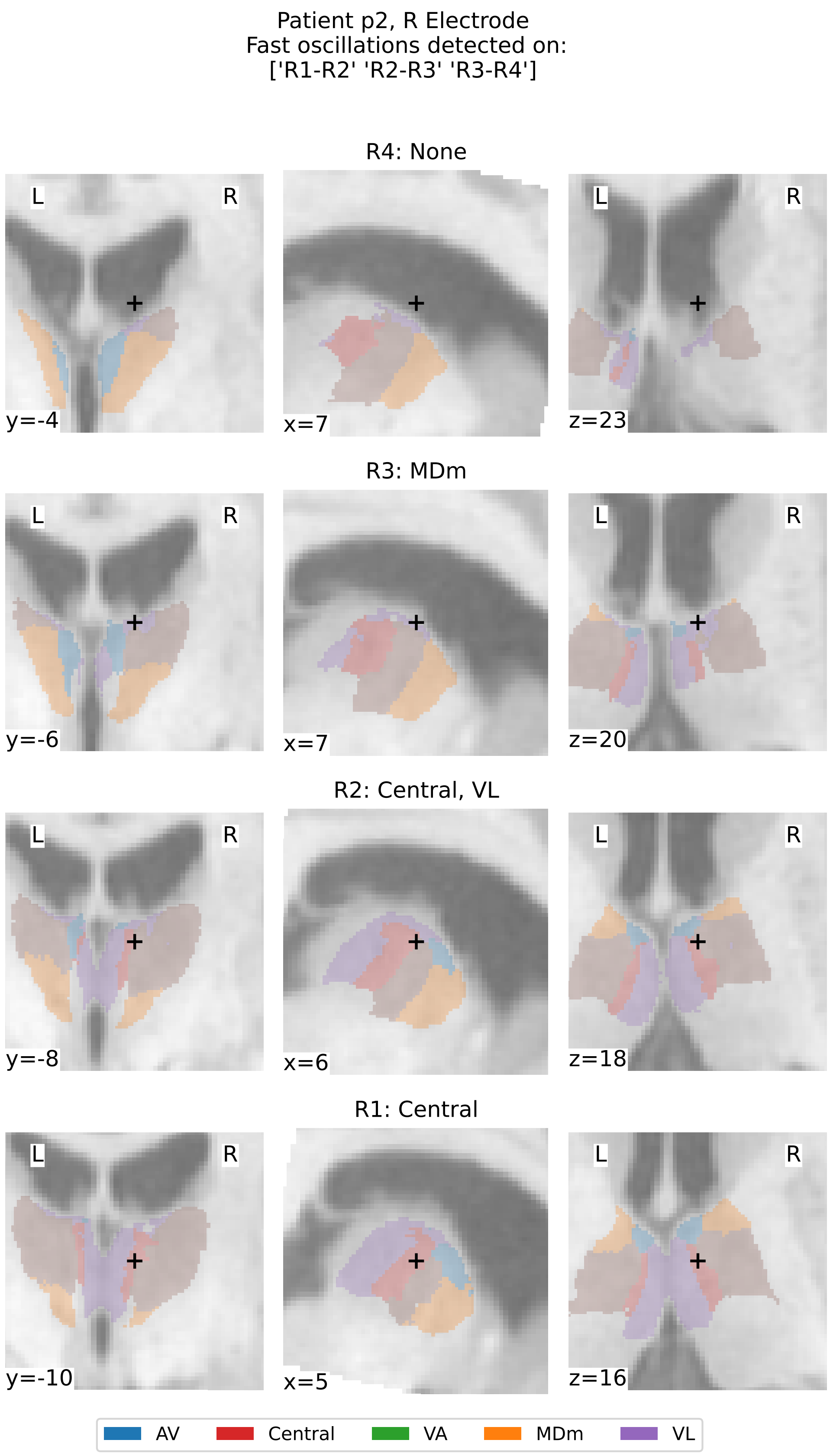


Supplementary Figure 15: **Thalamic Segmentation and Localization of contacts for patient p2, right electrode.** See caption of Supplementary Figure 12.

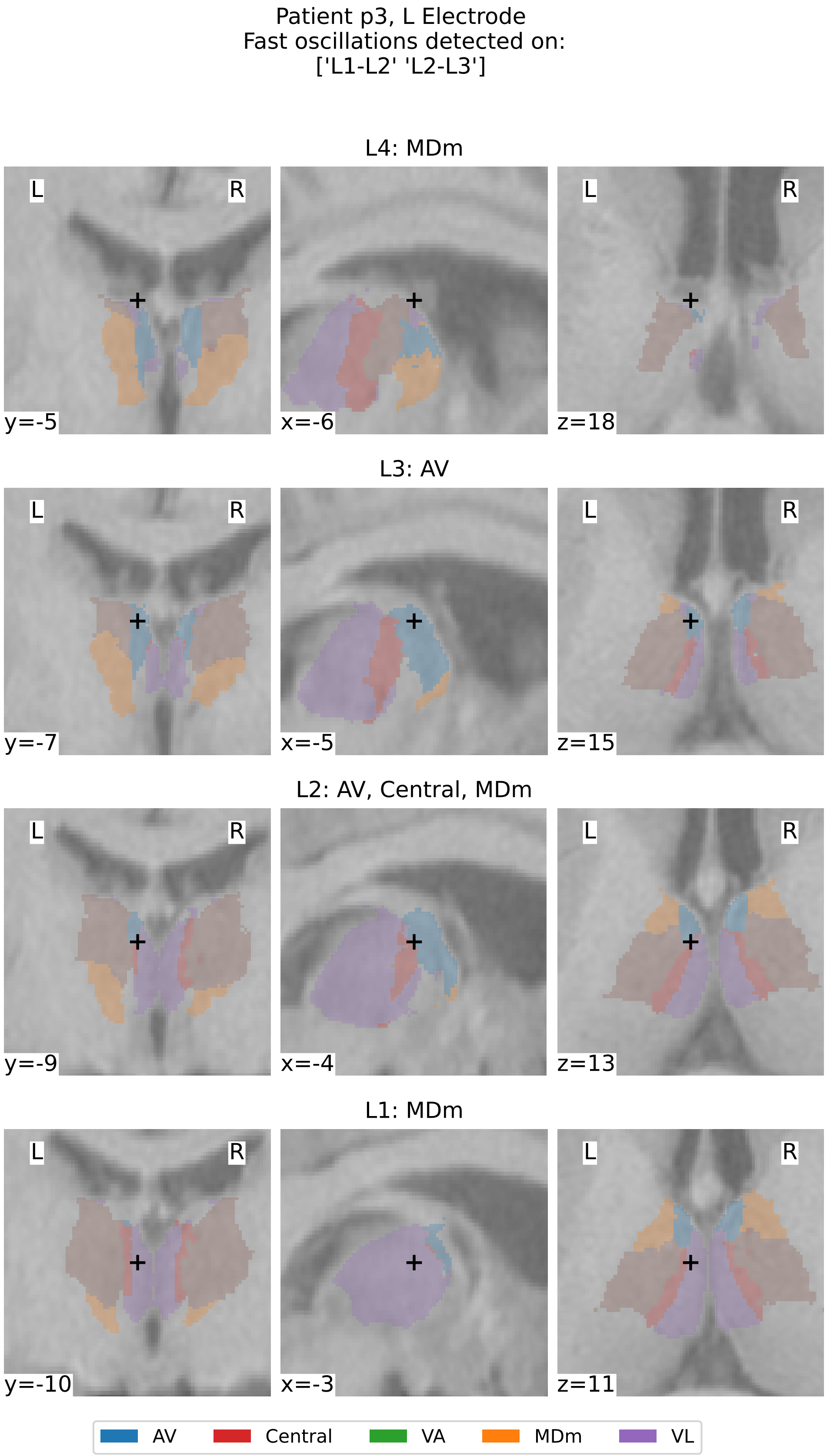


Supplementary Figure 16: **Thalamic Segmentation and Localization of contacts for patient p3, left electrode.** See caption of Supplementary Figure 12.

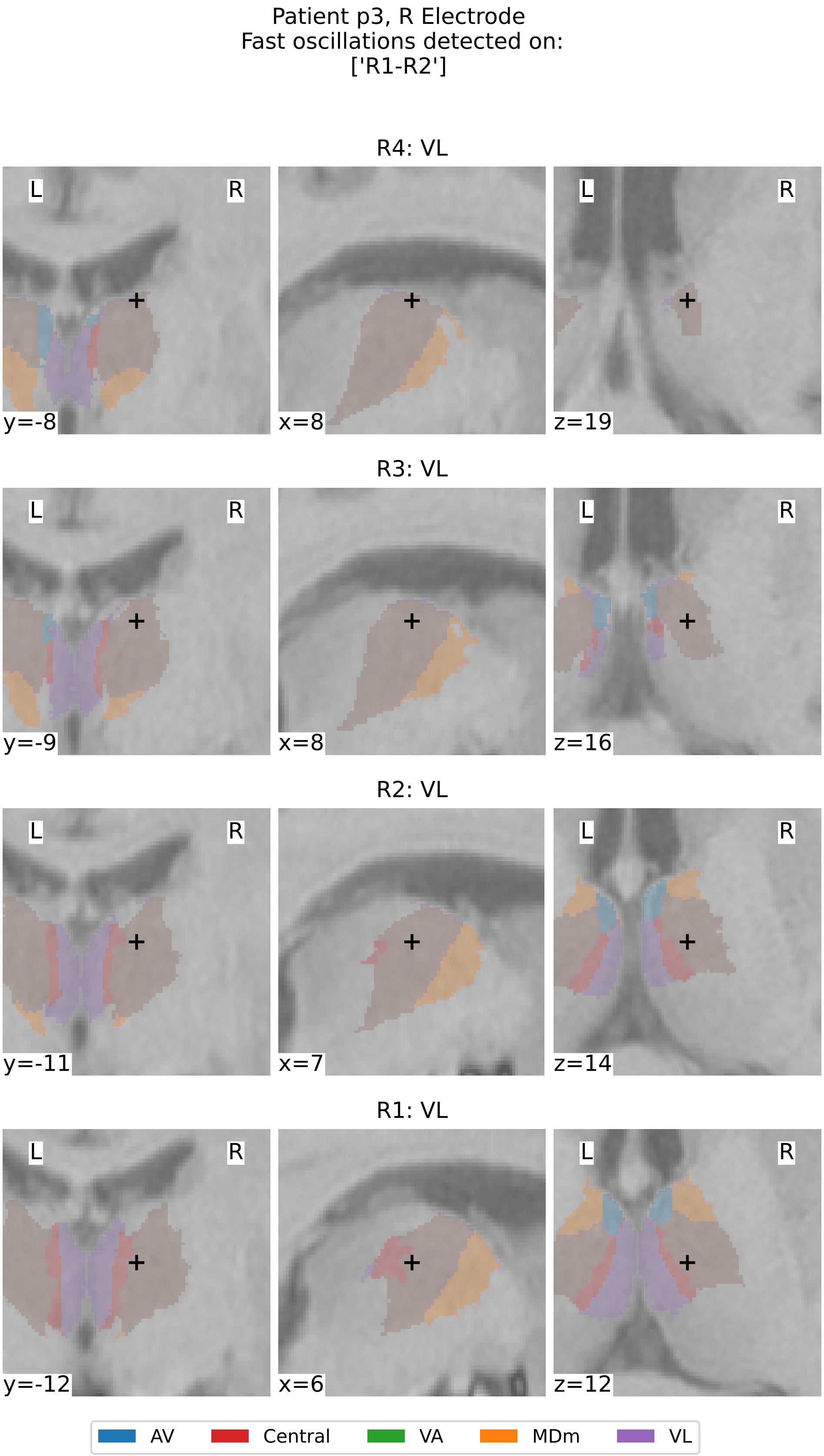


Supplementary Figure 17: **Thalamic Segmentation and Localization of contacts for patient p3, right electrode.** See caption of Supplementary Figure 12.

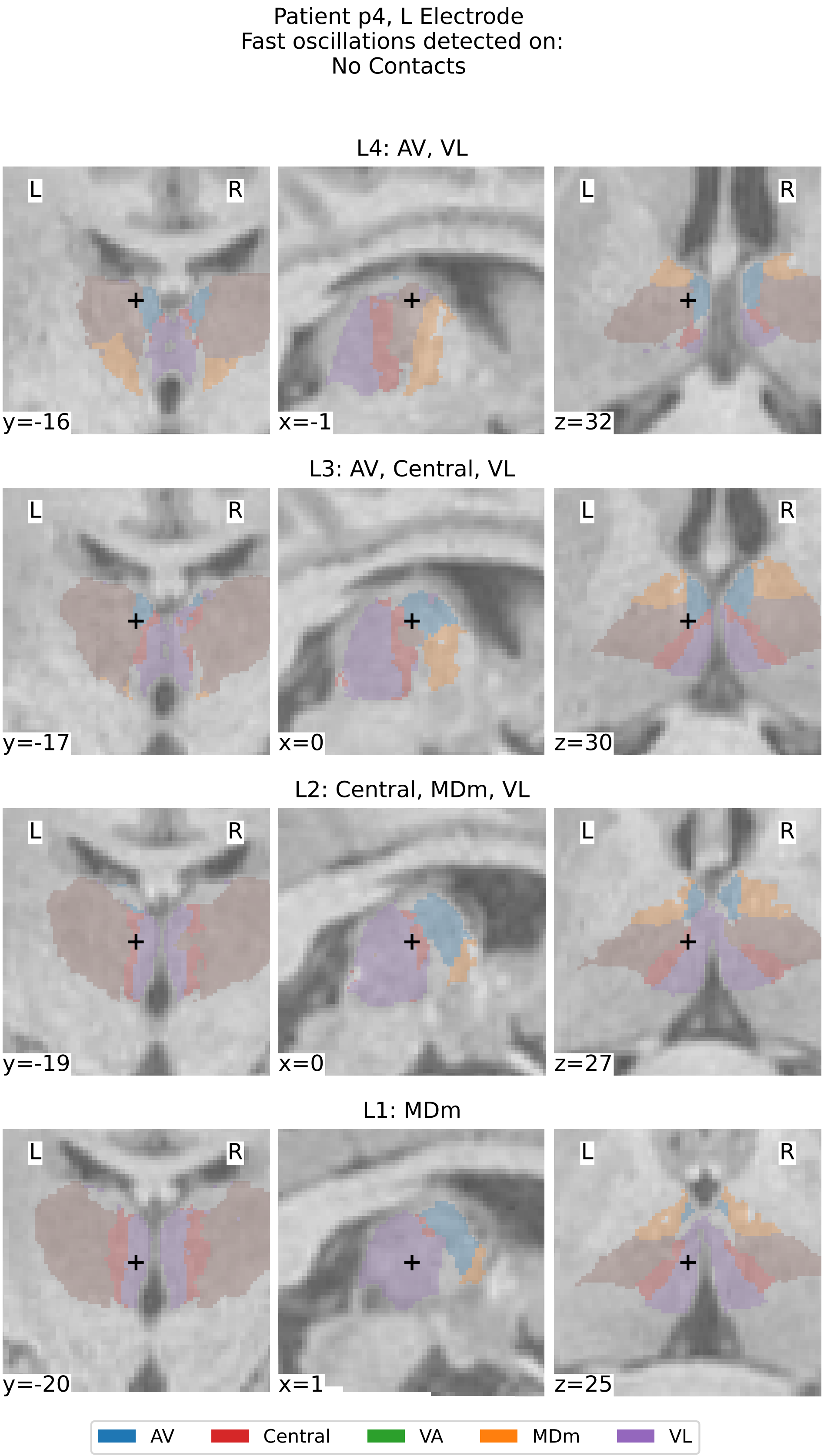


Supplementary Figure 18: **Thalamic Segmentation and Localization of contacts for patient p4, left electrode.** See caption of Supplementary Figure 12.

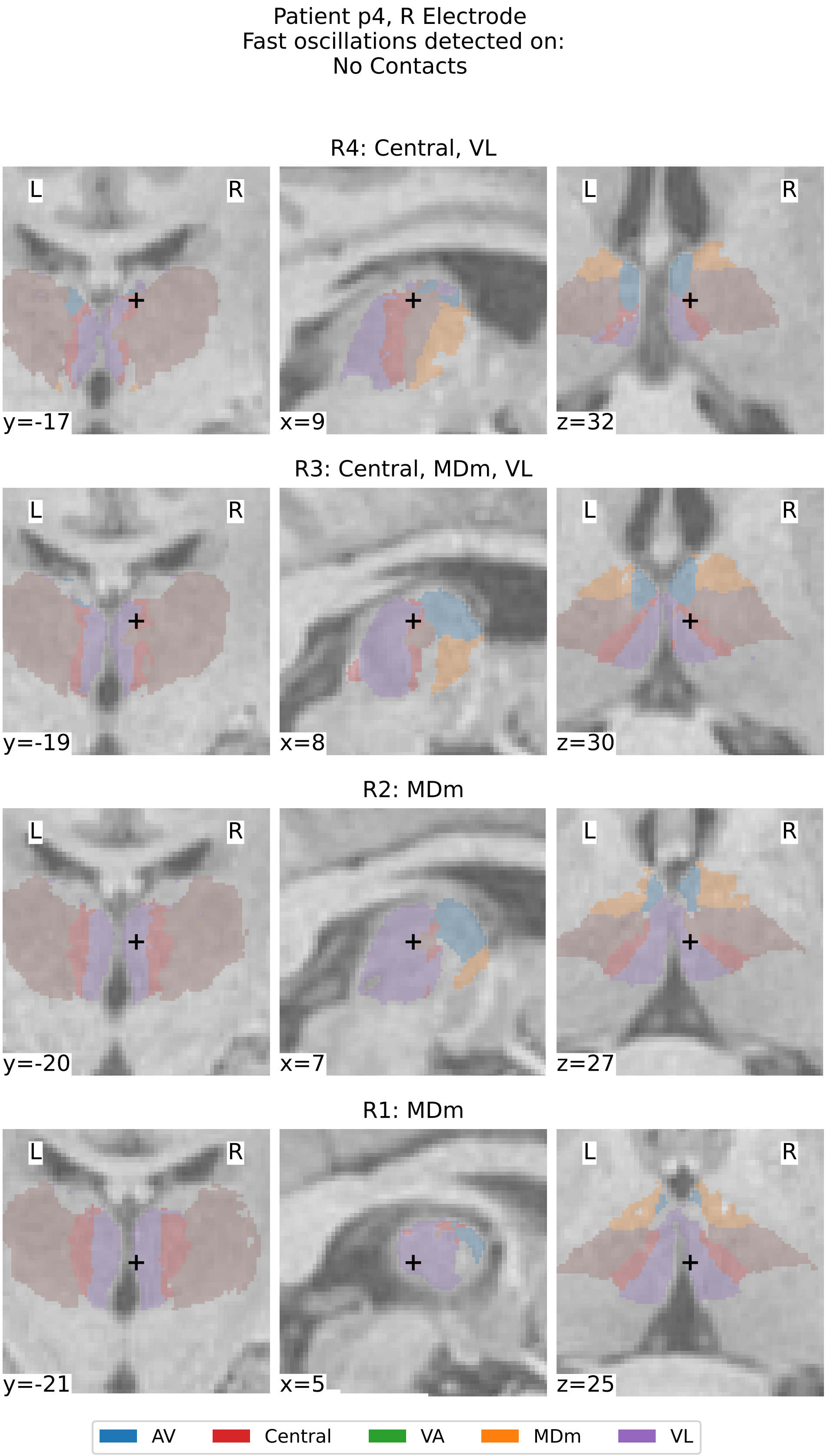


Supplementary Figure 19: **Thalamic Segmentation and Localization of contacts for patient p4, right electrode.** See caption of Supplementary Figure 12.

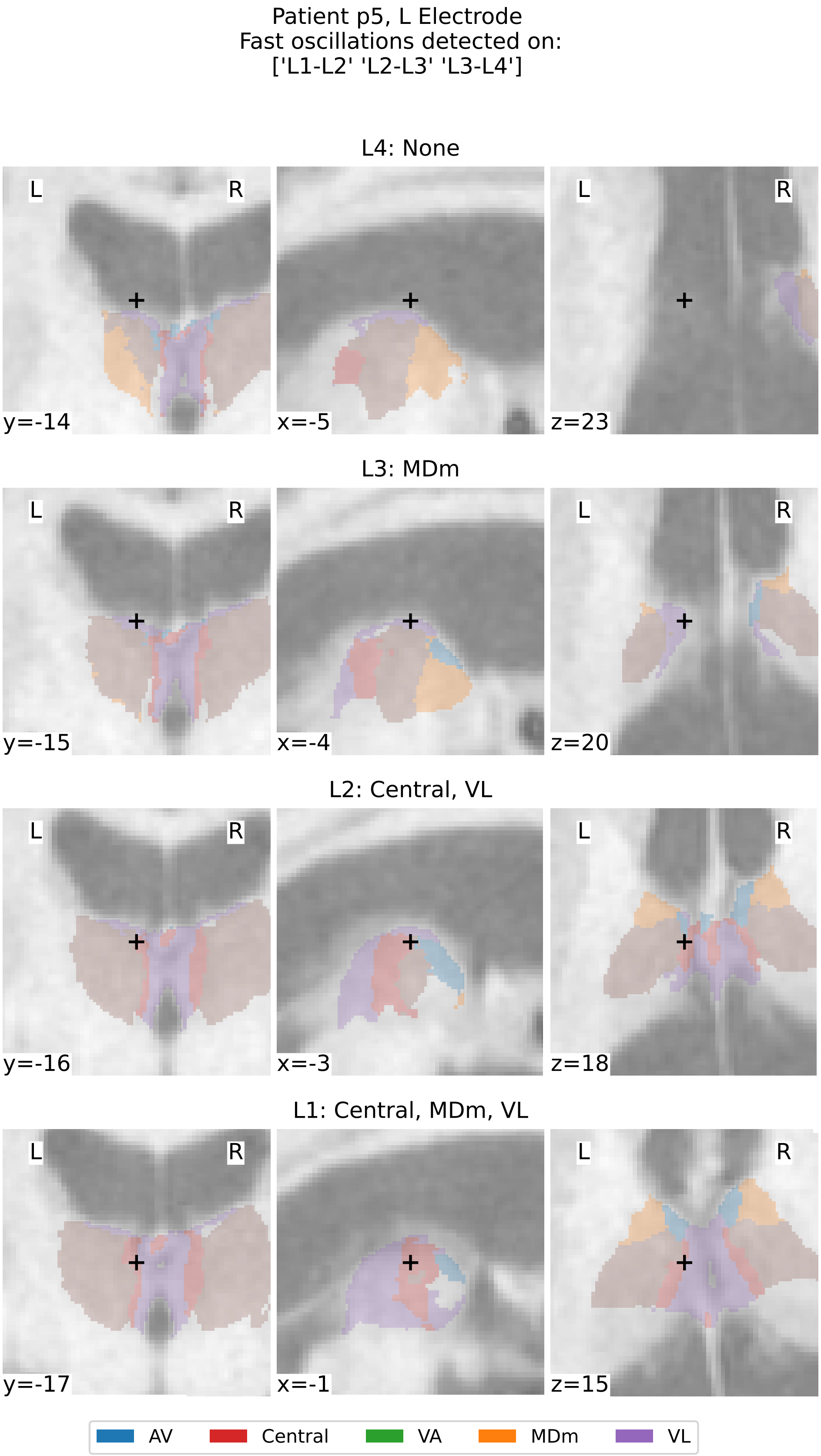


Supplementary Figure 20: **Thalamic Segmentation and Localization of contacts for patient p5, left electrode.** See caption of Supplementary Figure 12.

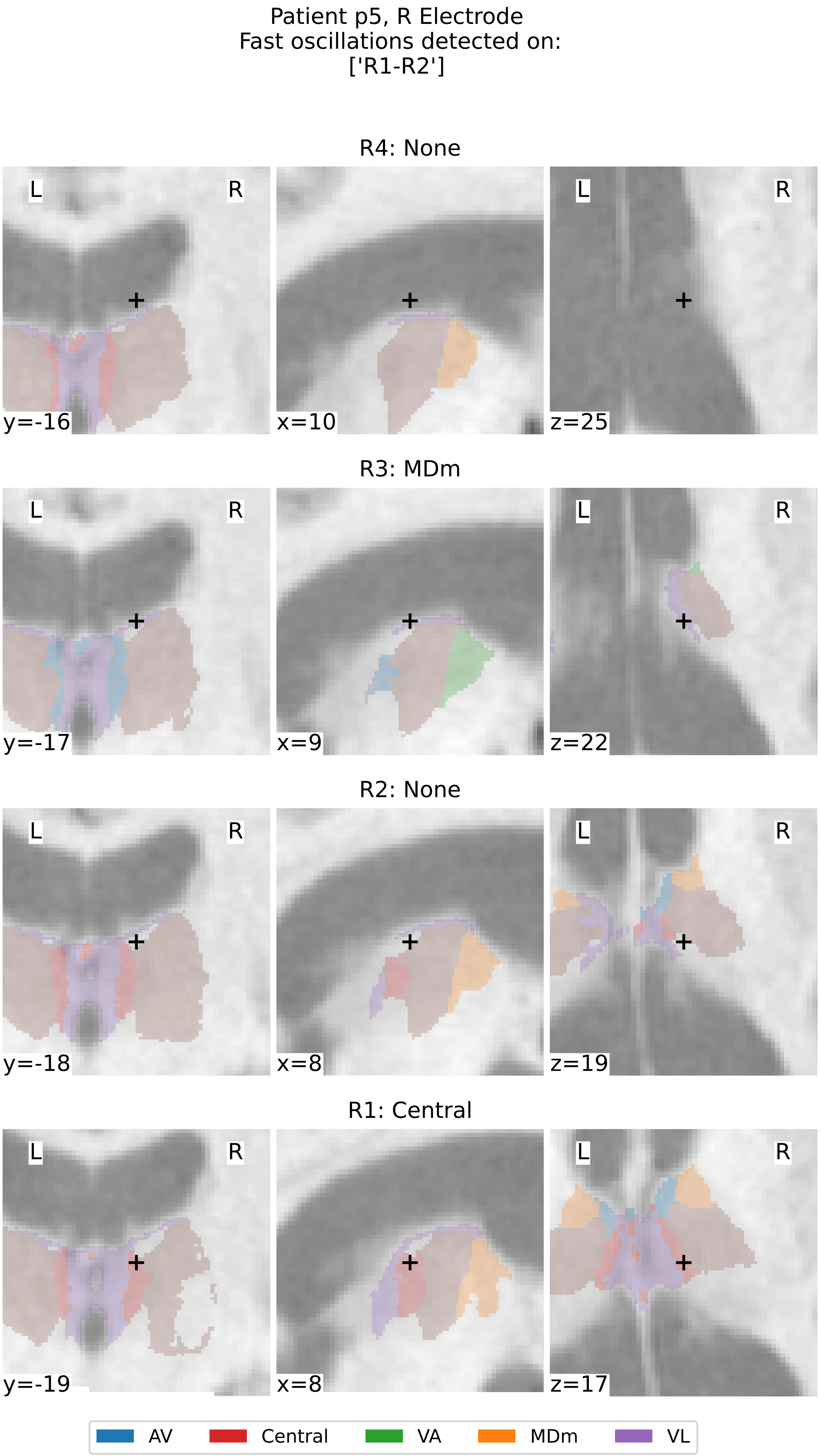


Supplementary Figure 21: **Thalamic Segmentation and Localization of contacts for patient p5, right electrode.** See caption of Supplementary Figure 12.

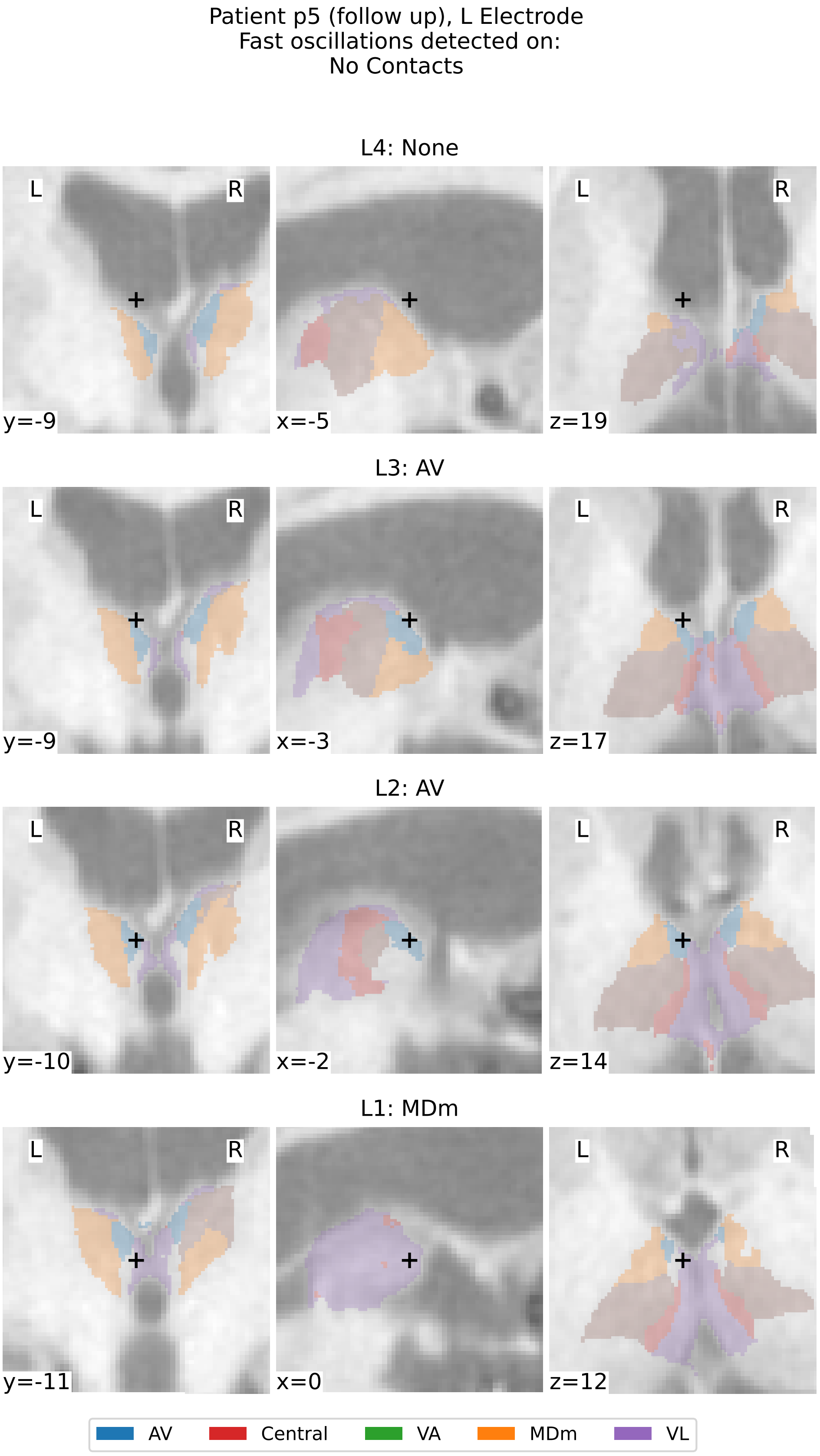


Supplementary Figure 22: **Thalamic Segmentation and Localization of contacts for patient p5 (reimplanted epoch), left electrode.** See caption of Supplementary Figure 12.

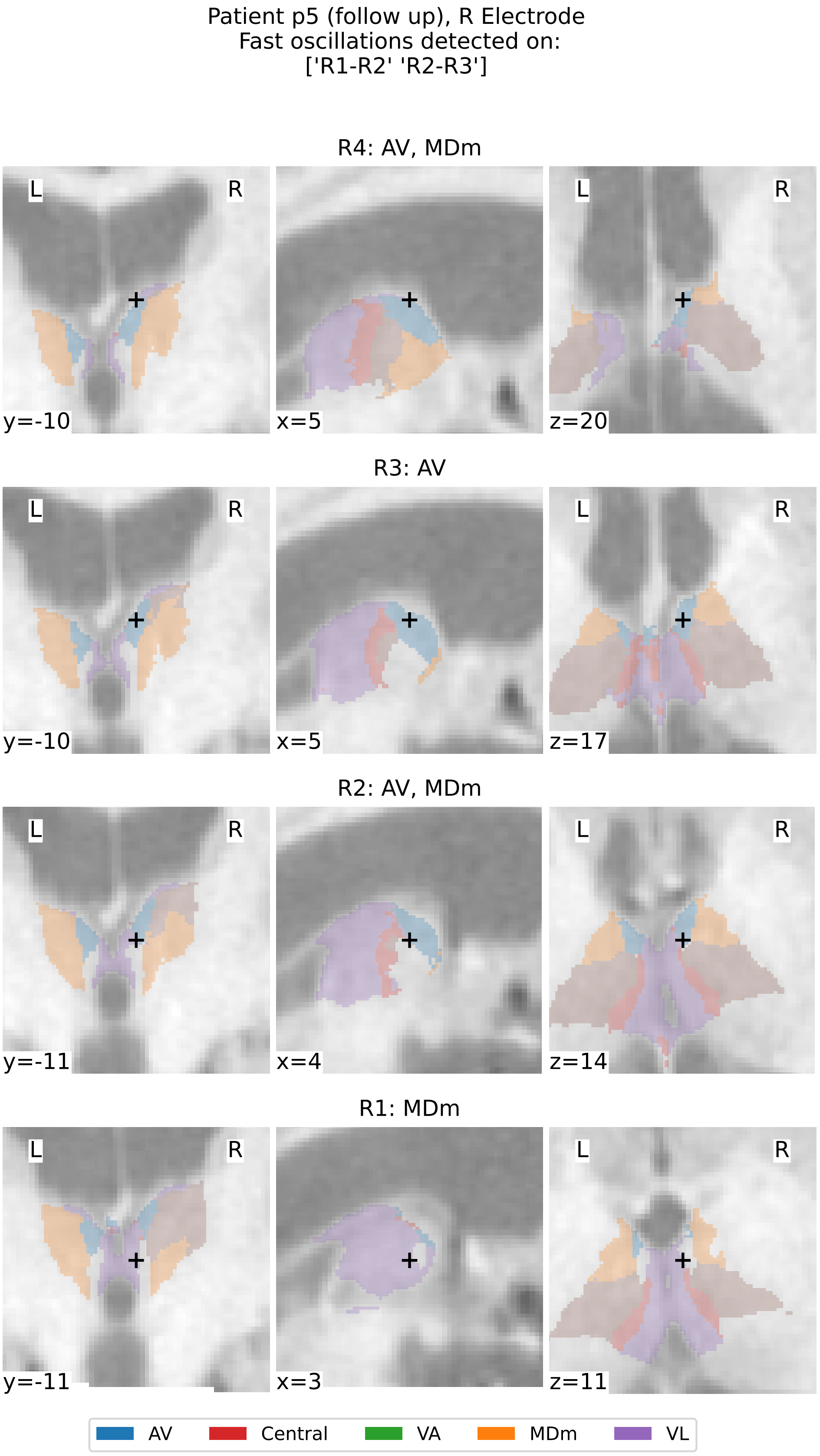


Supplementary Figure 23: **Thalamic Segmentation and Localization of contacts for patient p5 (reimplanted epoch), right electrode.** See caption of Supplementary Figure 12.

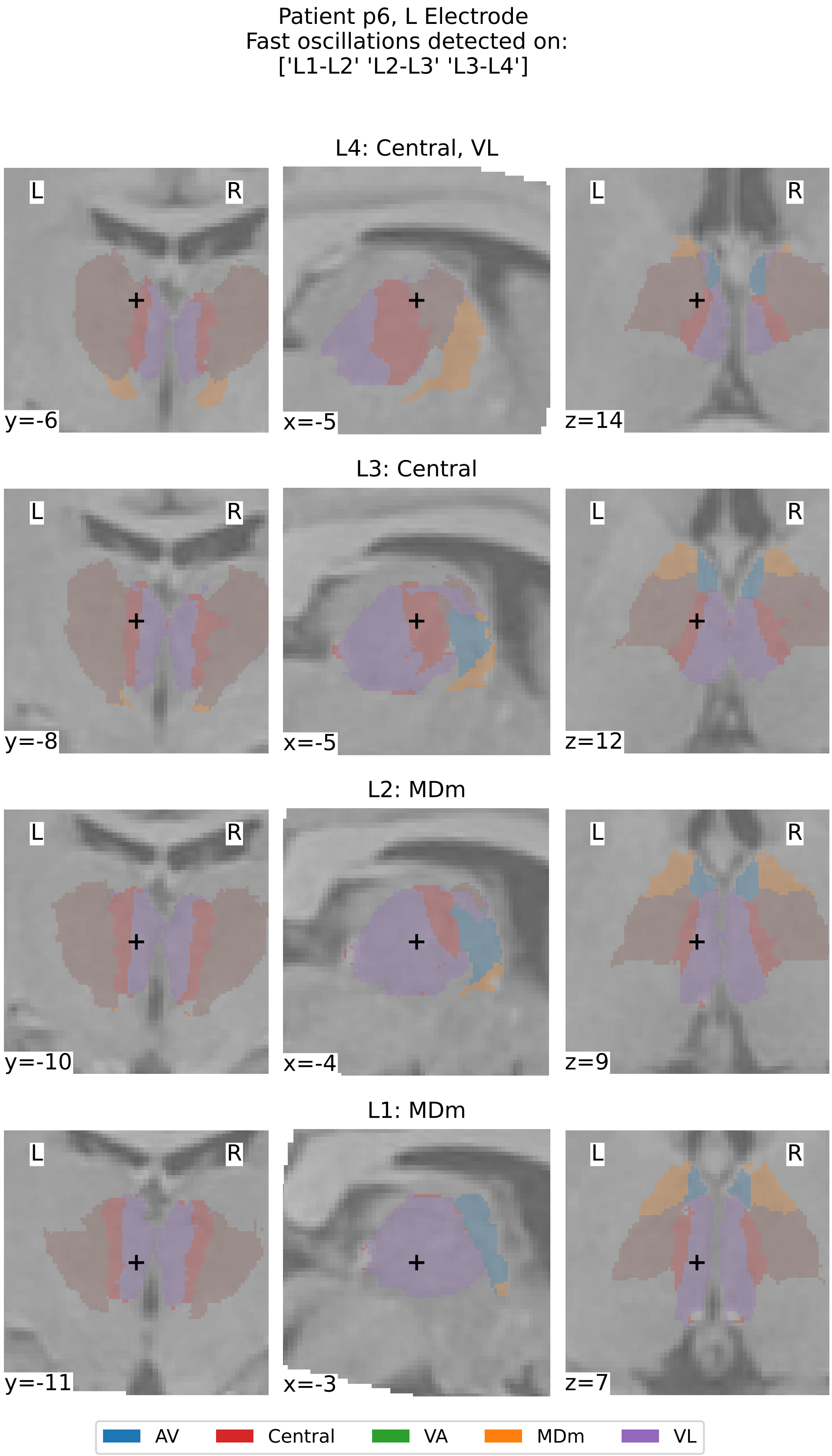


Supplementary Figure 24: **Thalamic Segmentation and Localization of contacts for patient p6, left electrode.** See caption of Supplementary Figure 12.

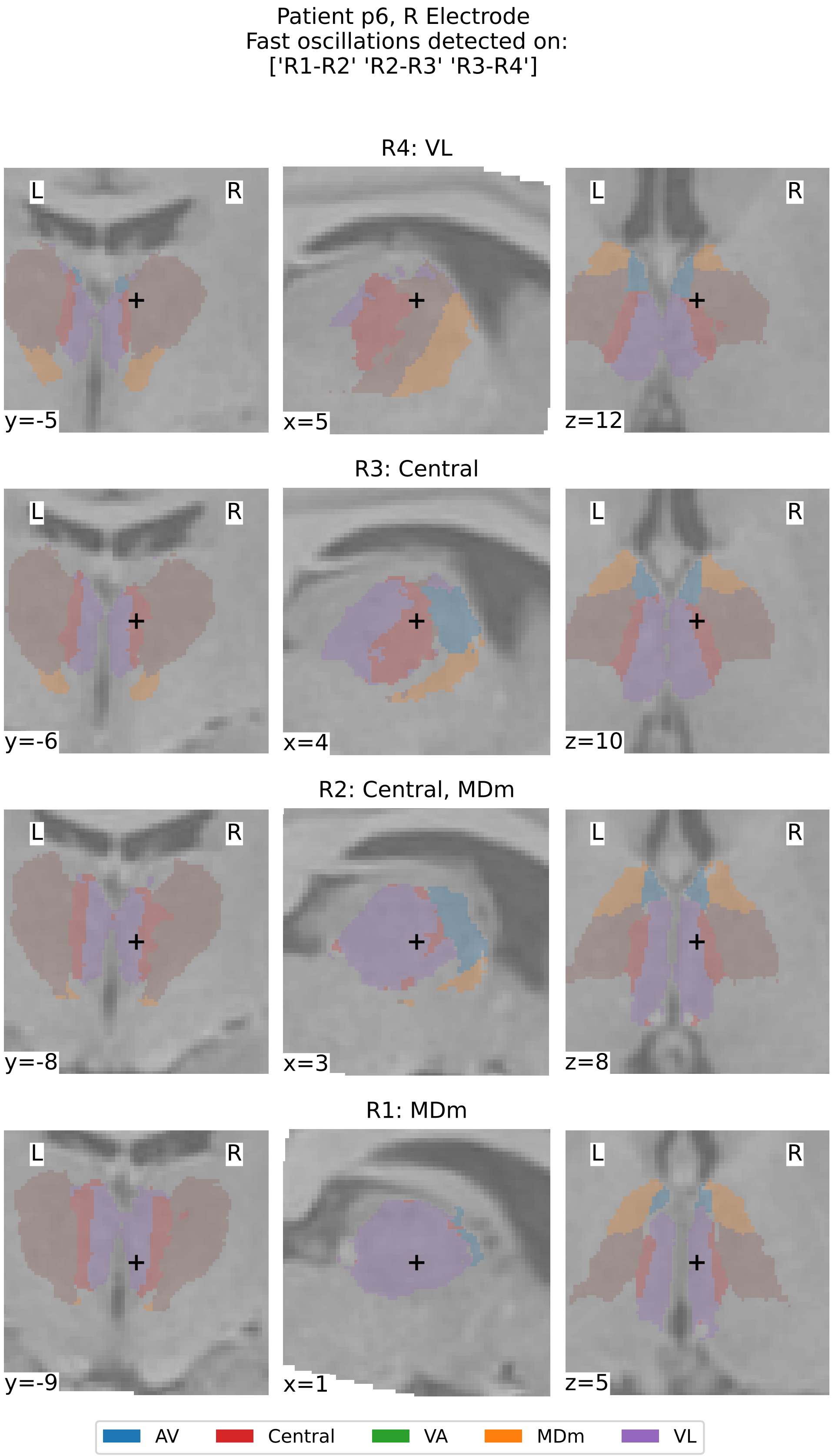


Supplementary Figure 25: **Thalamic Segmentation and Localization of contacts for patient p6, right electrode.** See caption of Supplementary Figure 12.

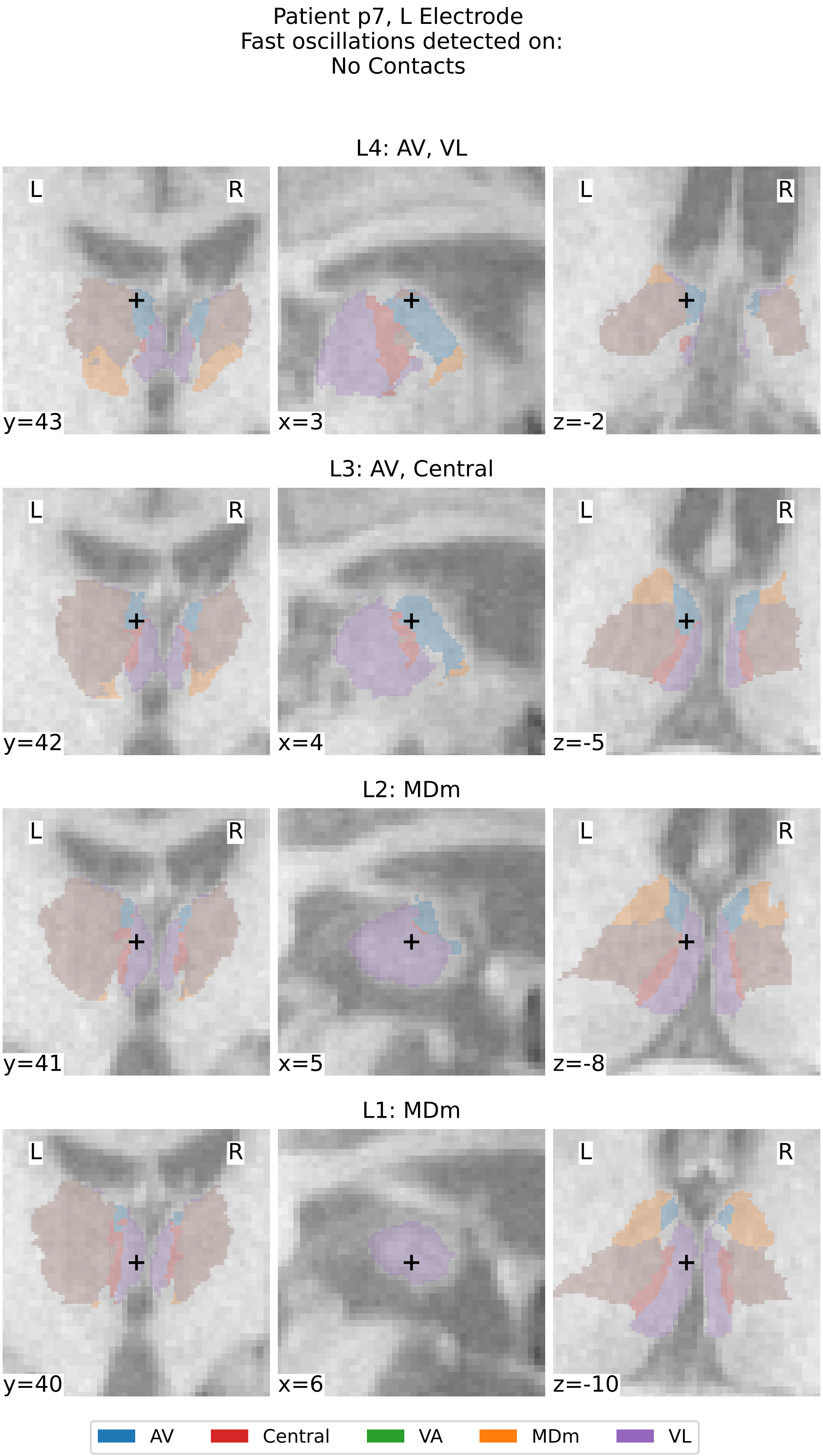


Supplementary Figure 26: **Thalamic Segmentation and Localization of contacts for patient p7, left electrode.** See caption of Supplementary Figure 12.

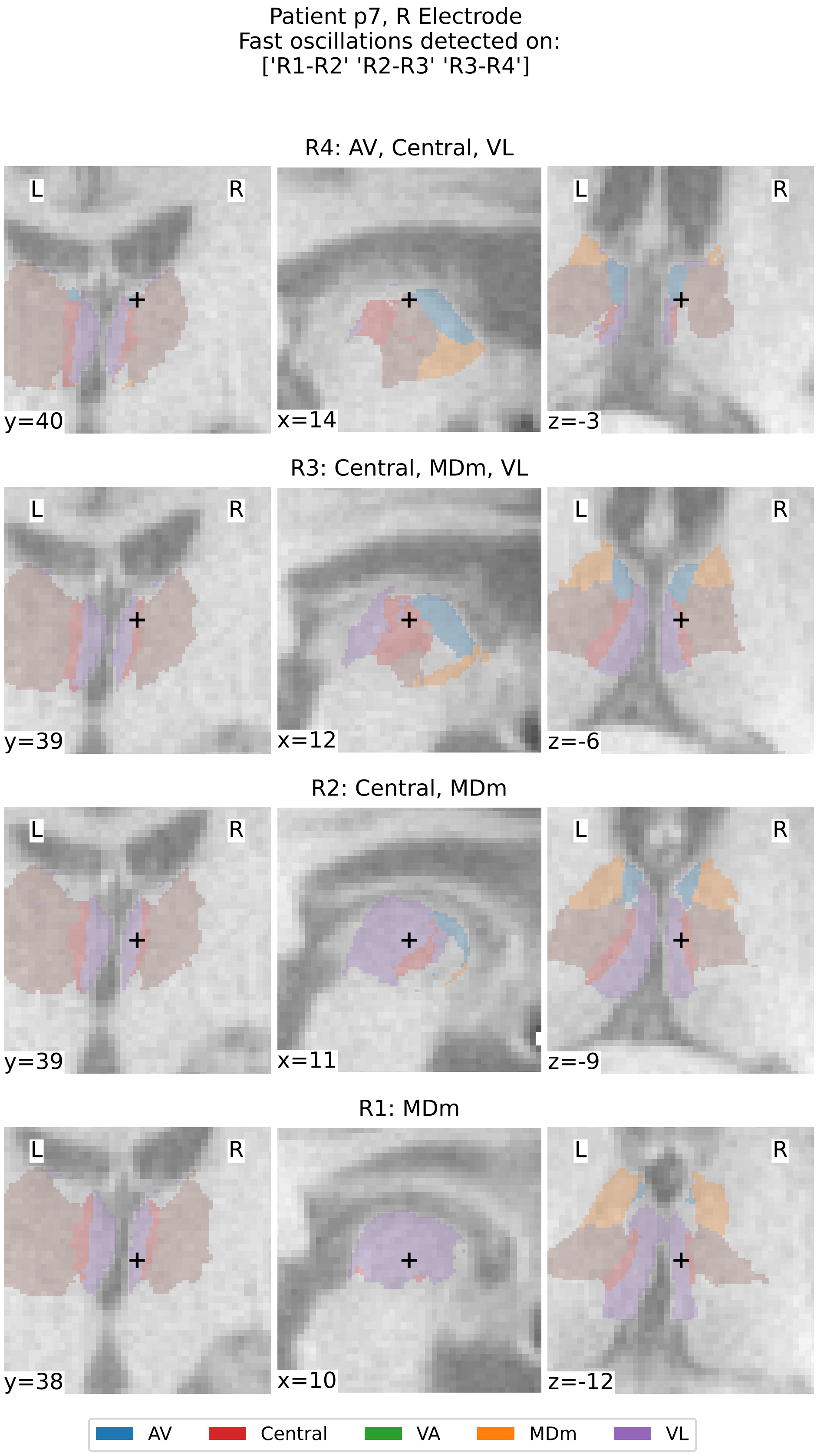


Supplementary Figure 27: **Thalamic Segmentation and Localization of contacts for patient p7, right electrode.** See caption of Supplementary Figure 12.

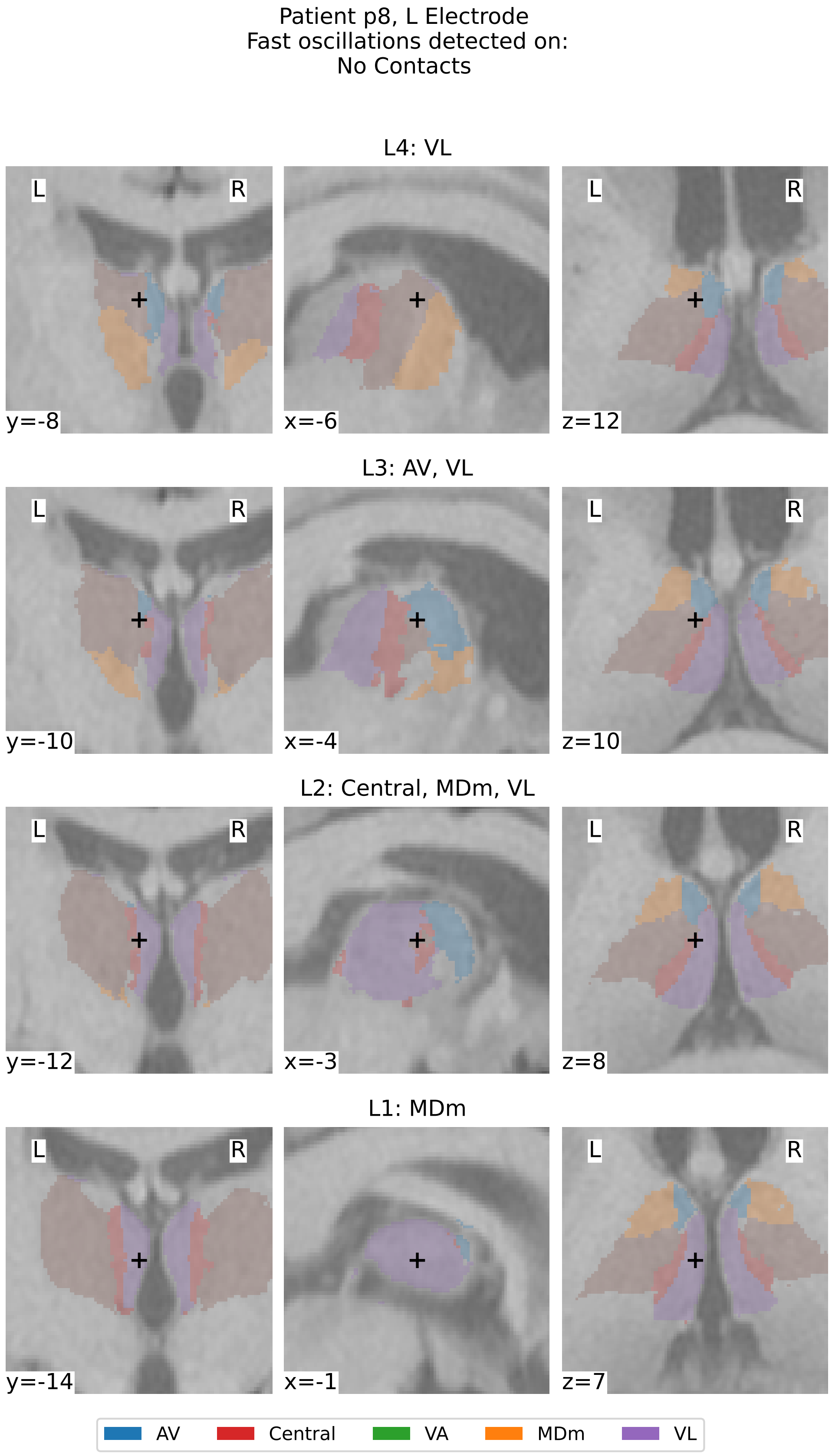


Supplementary Figure 28: **Thalamic Segmentation and Localization of contacts for patient p8, left electrode.** See caption of Supplementary Figure 12.

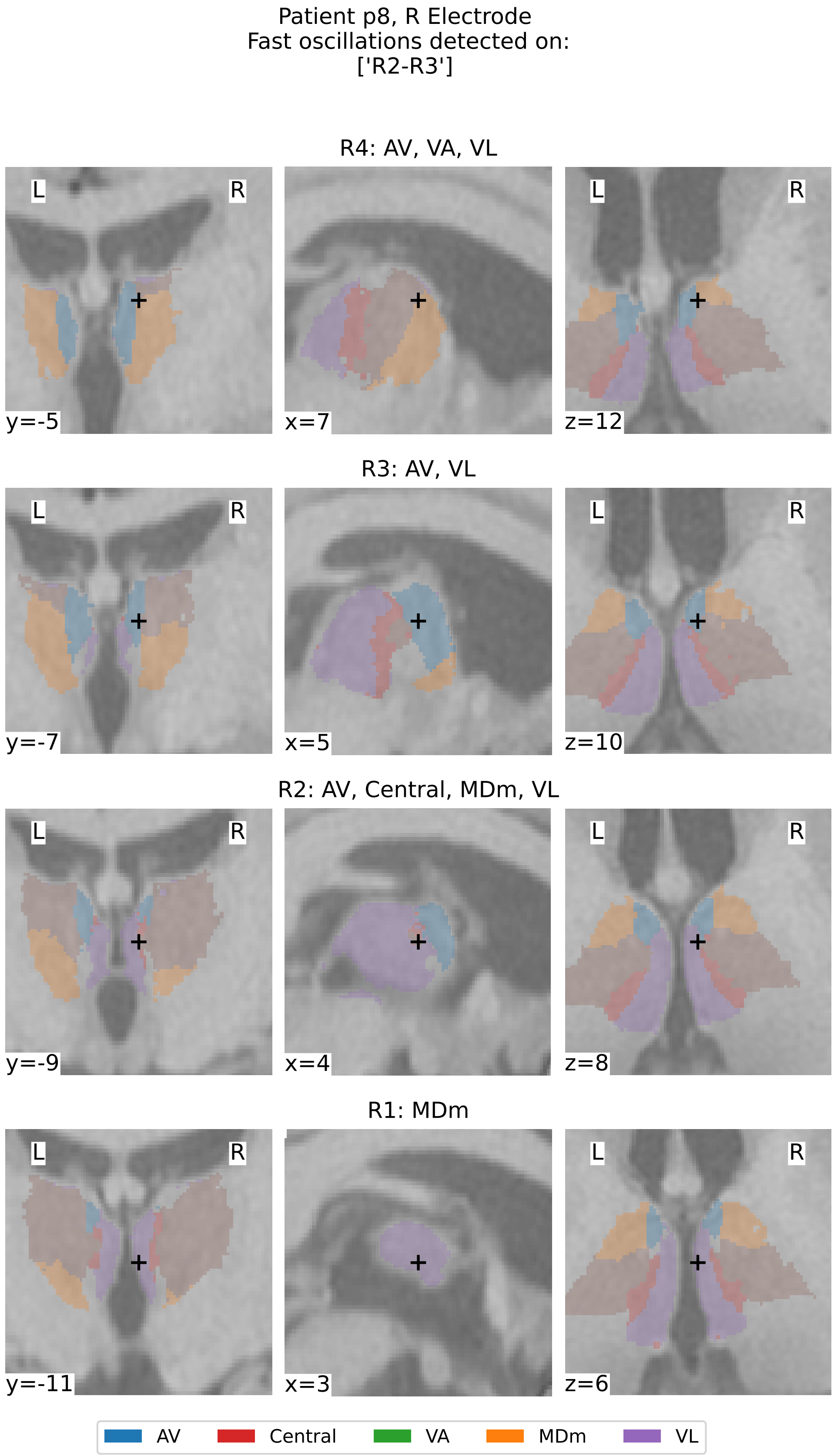


Supplementary Figure 29: **Thalamic Segmentation and Localization of contacts for patient p8, right electrode.** See caption of Supplementary Figure 12.

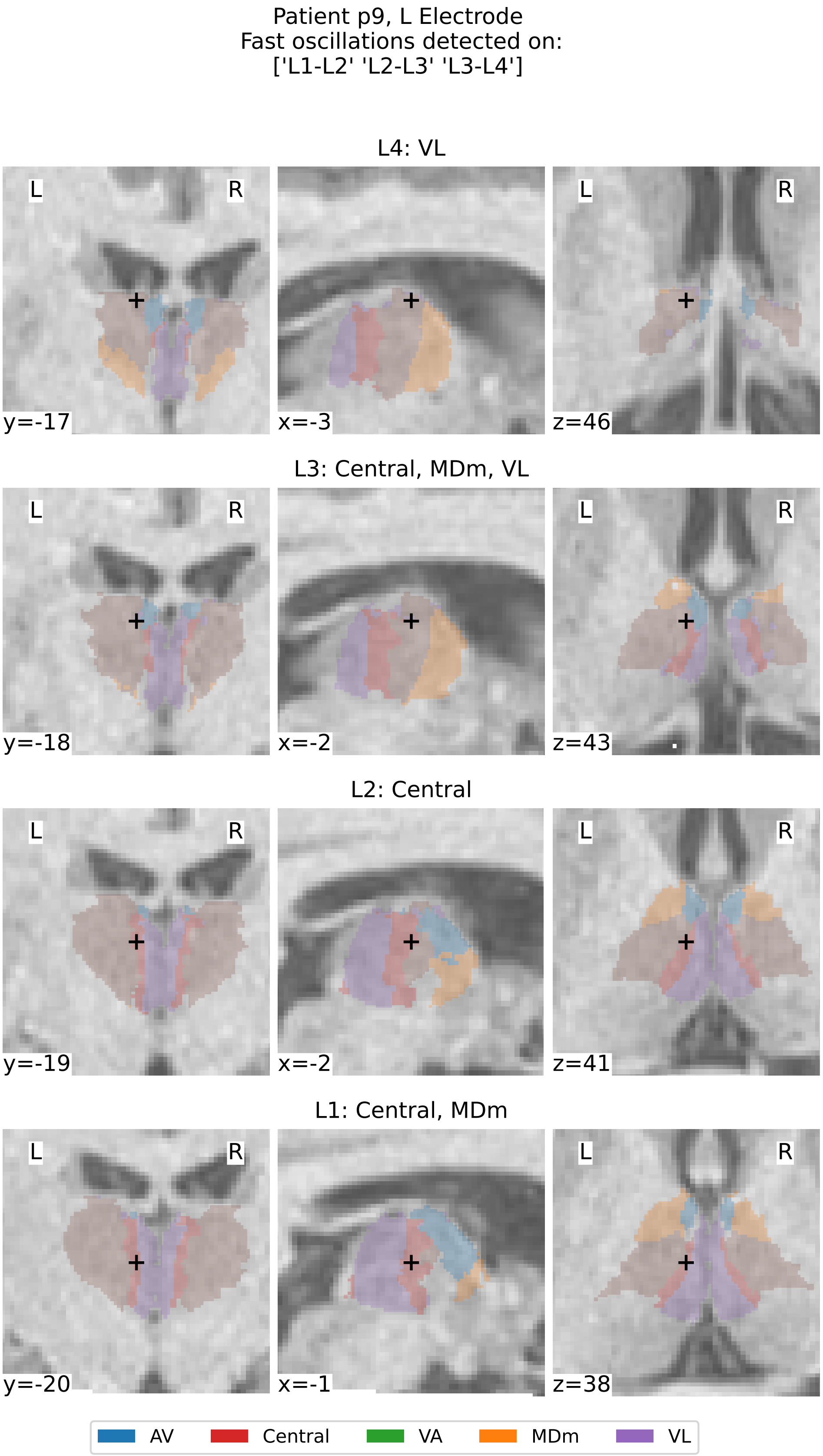


Supplementary Figure 30: **Thalamic Segmentation and Localization of contacts for patient p9, left electrode.** See caption of Supplementary Figure 12.

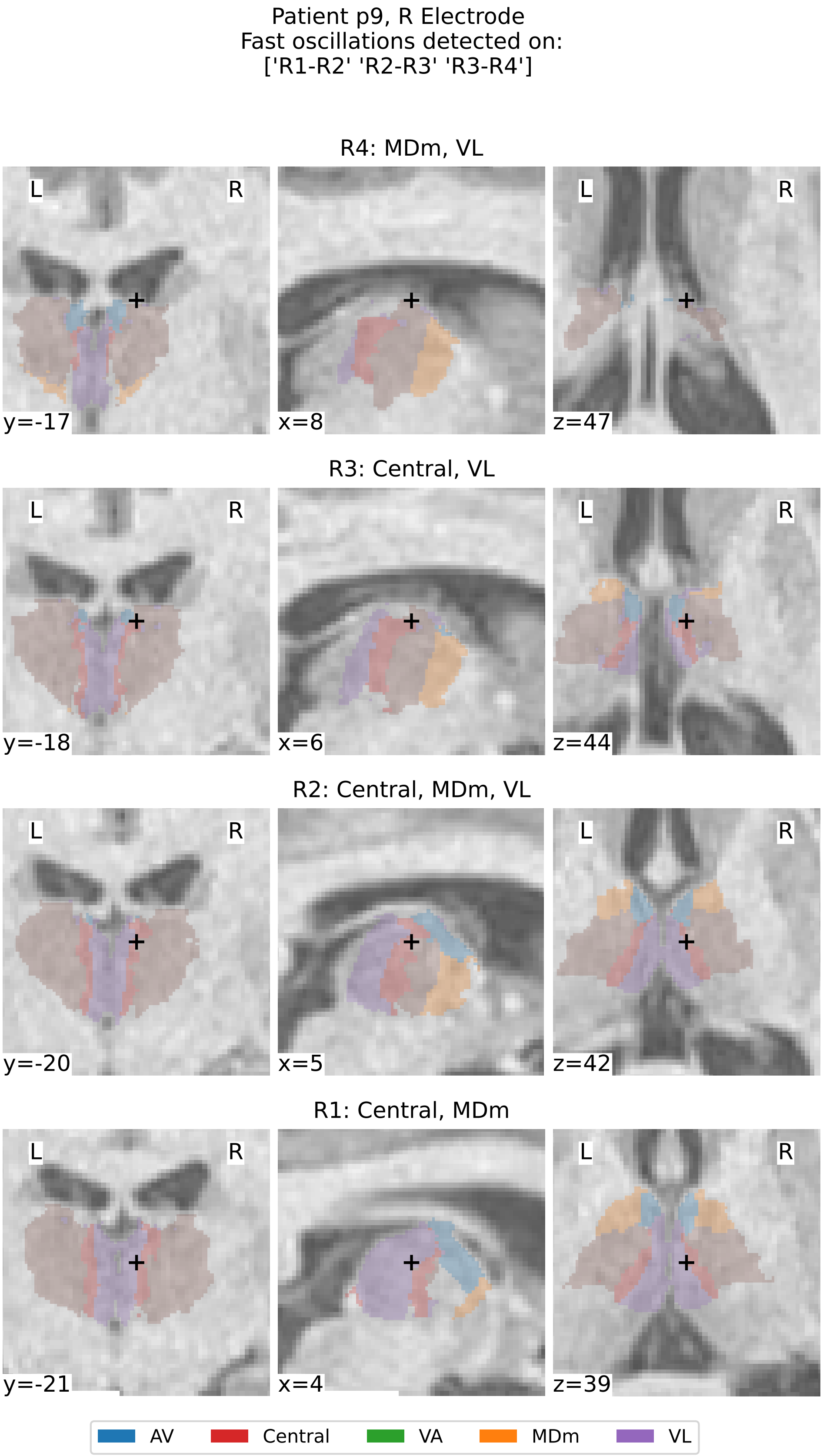


Supplementary Figure 31: **Thalamic Segmentation and Localization of contacts for patient p9, right electrode.** See caption of Supplementary Figure 12.

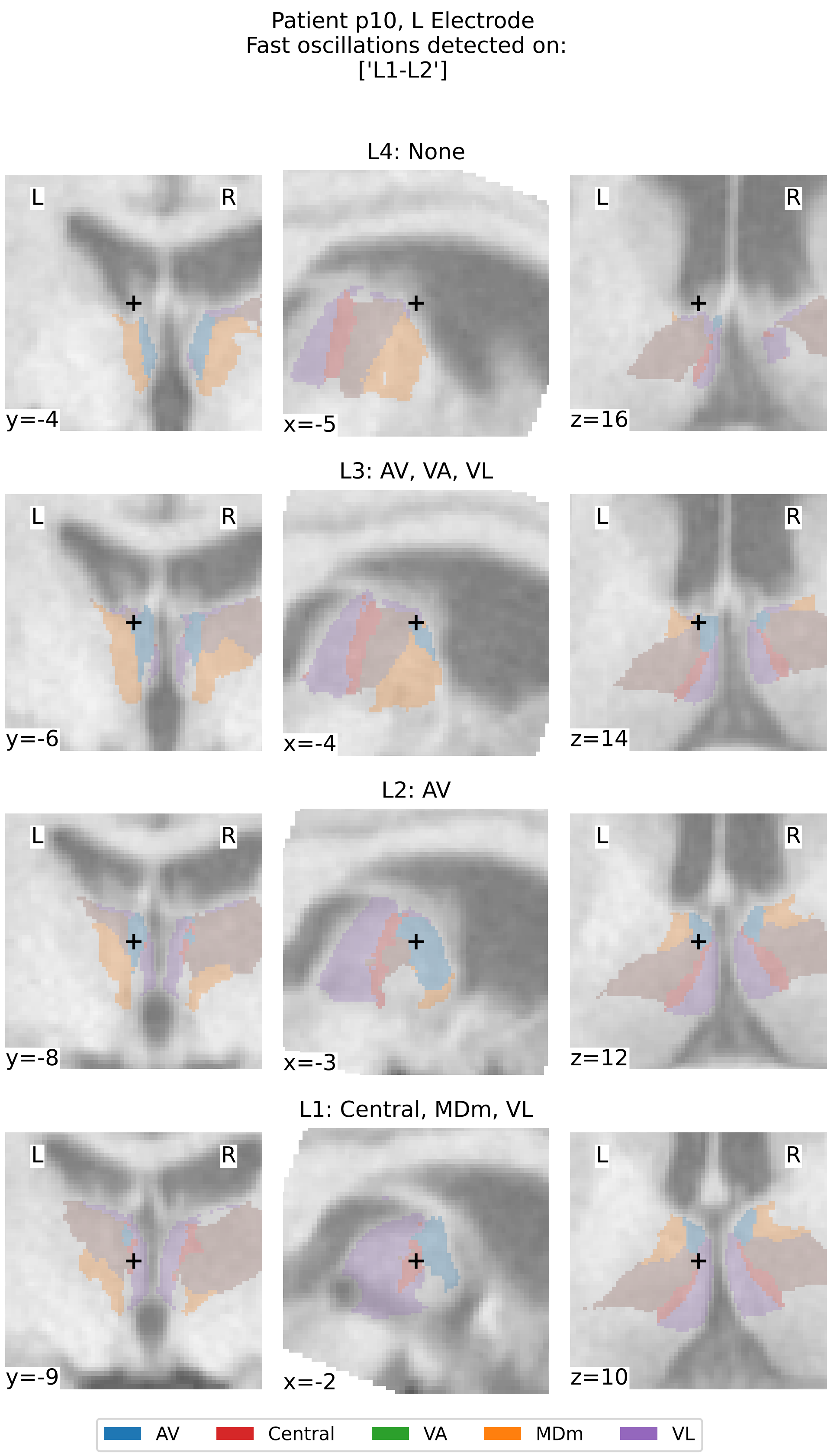


Supplementary Figure 32: **Thalamic Segmentation and Localization of contacts for patient p10, left electrode.** See caption of Supplementary Figure 12.

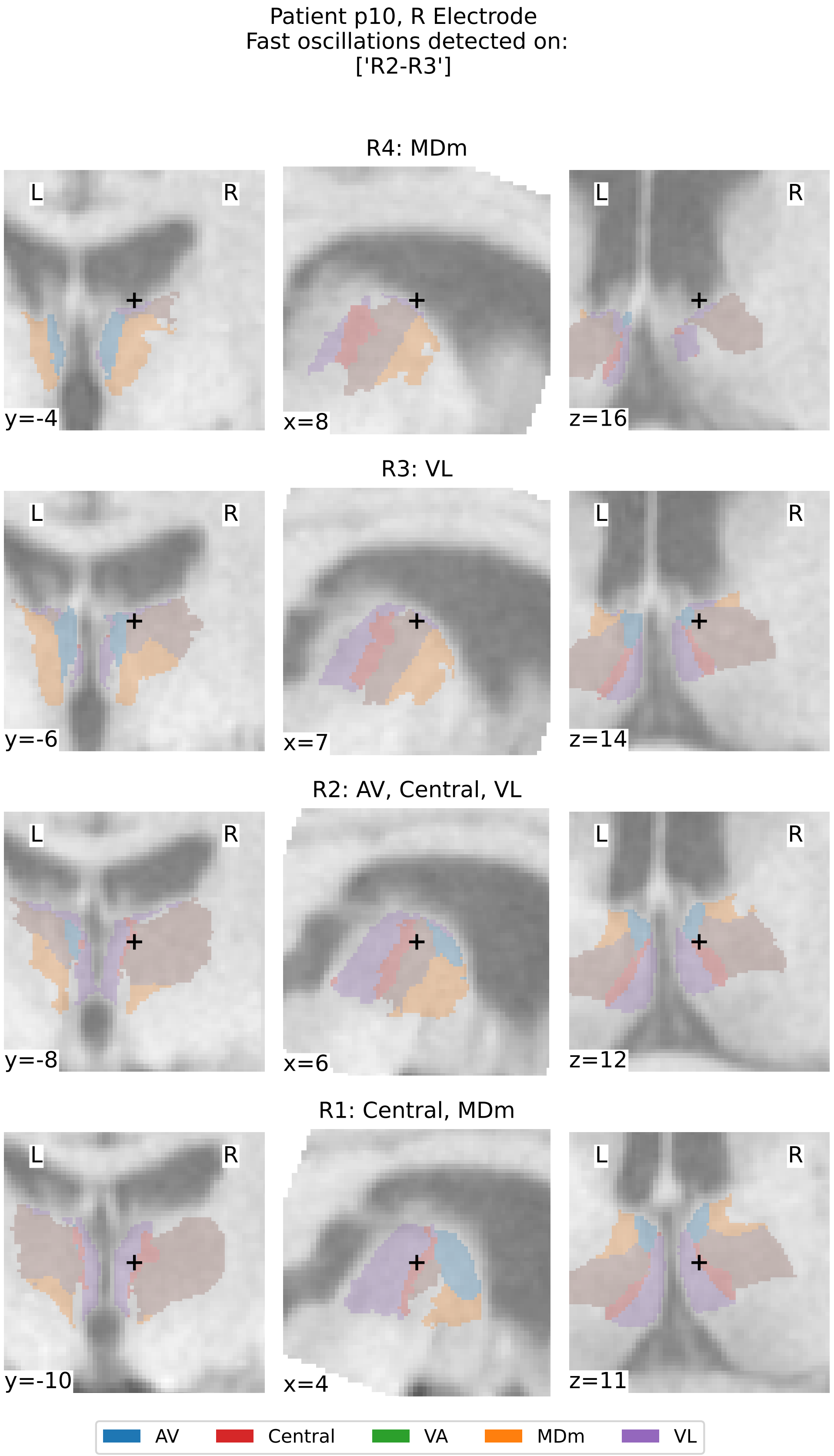


Supplementary Figure 33: **Thalamic Segmentation and Localization of contacts for patient p10, right electrode.** See caption of Supplementary Figure 12.

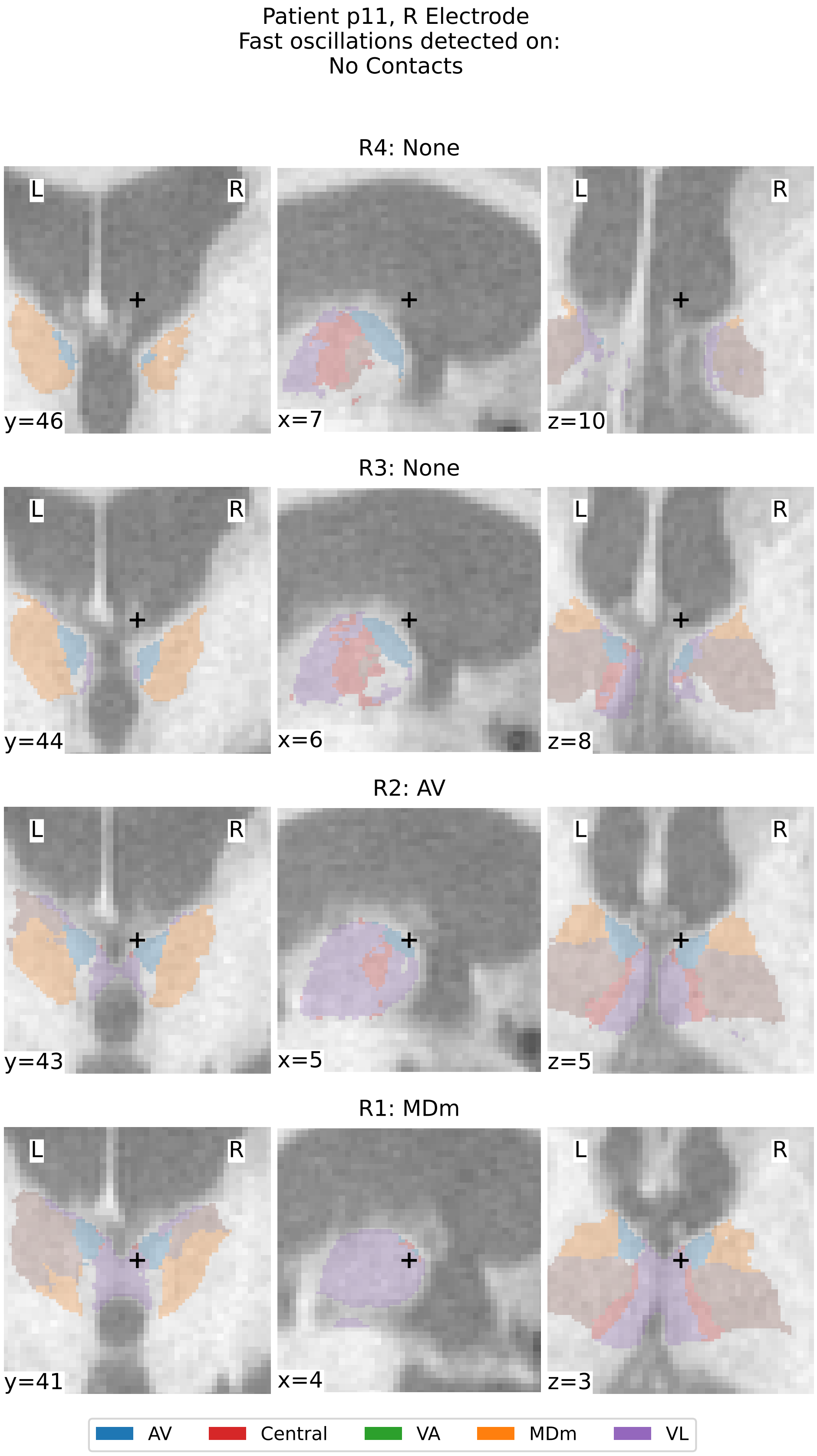


Supplementary Figure 34: **Thalamic Segmentation and Localization of contacts for patient p11, right electrode.** See caption of Supplementary Figure 12.

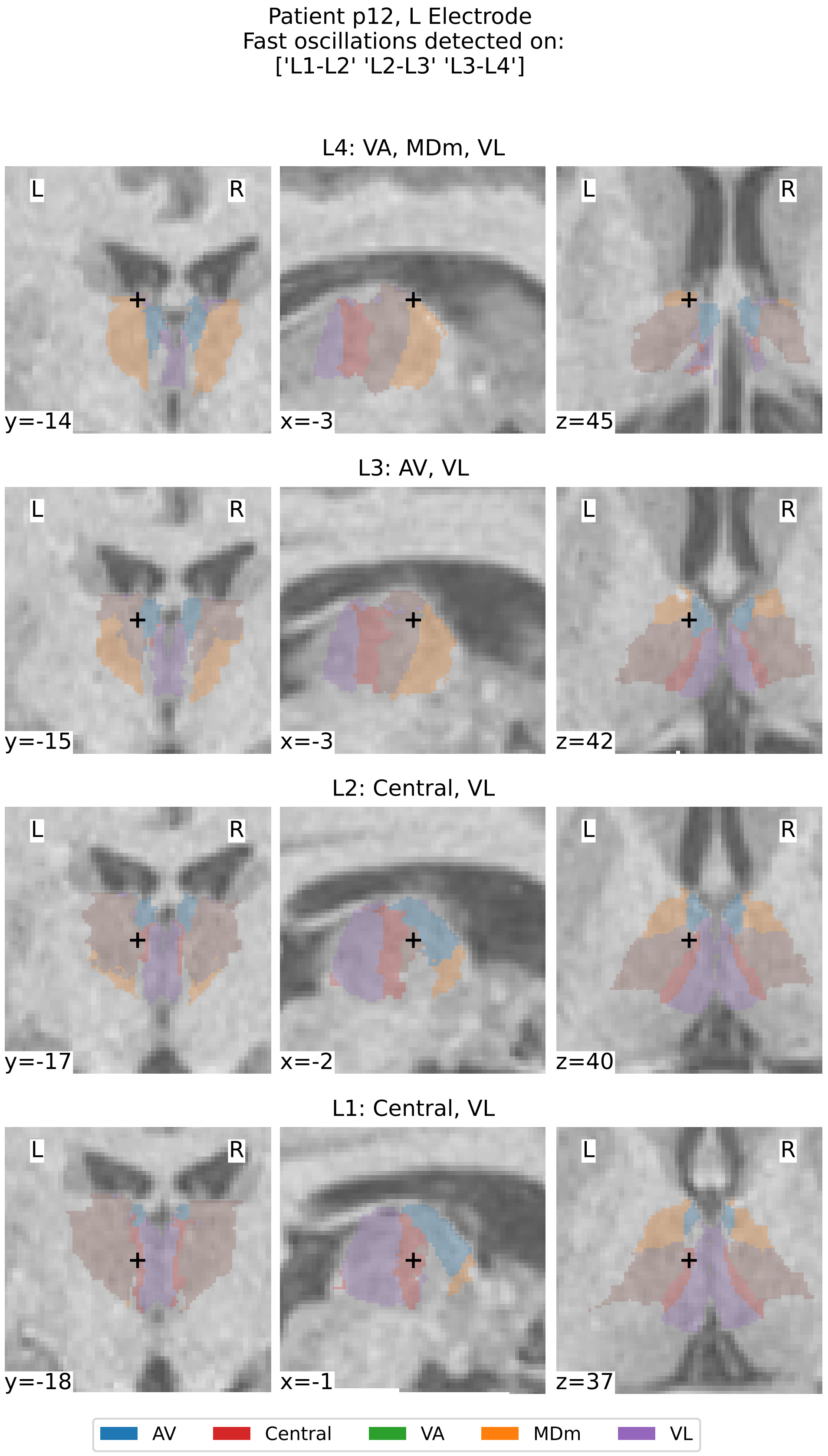


Supplementary Figure 35: **Thalamic Segmentation and Localization of contacts for patient p12, left electrode.** See caption of Supplementary Figure 12.

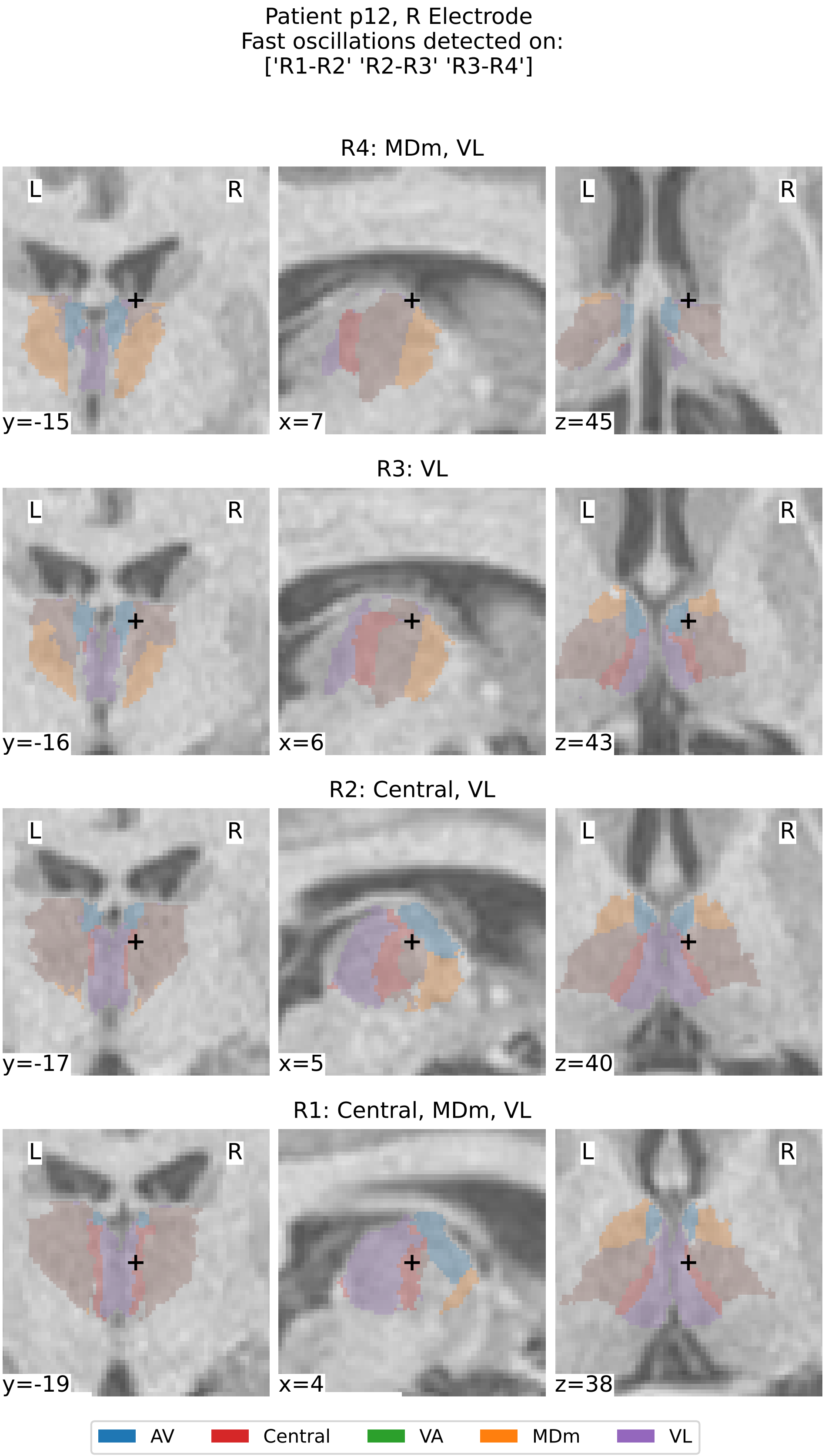


Supplementary Figure 36: **Thalamic Segmentation and Localization of contacts for patient p12, right electrode.** See caption of Supplementary Figure 12.

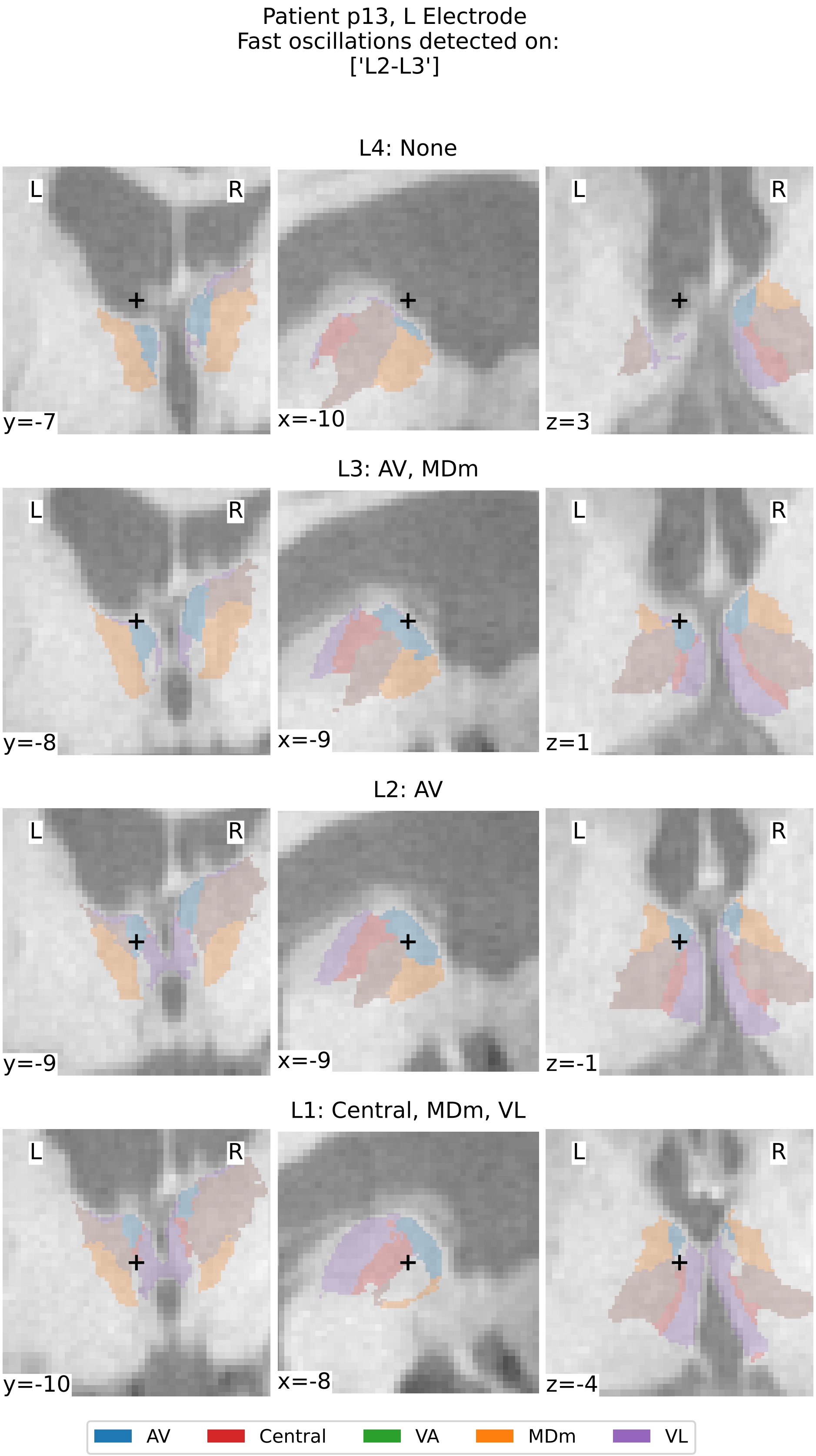


Supplementary Figure 37: **Thalamic Segmentation and Localization of contacts for patient p13, left electrode.** See caption of Supplementary Figure 12.

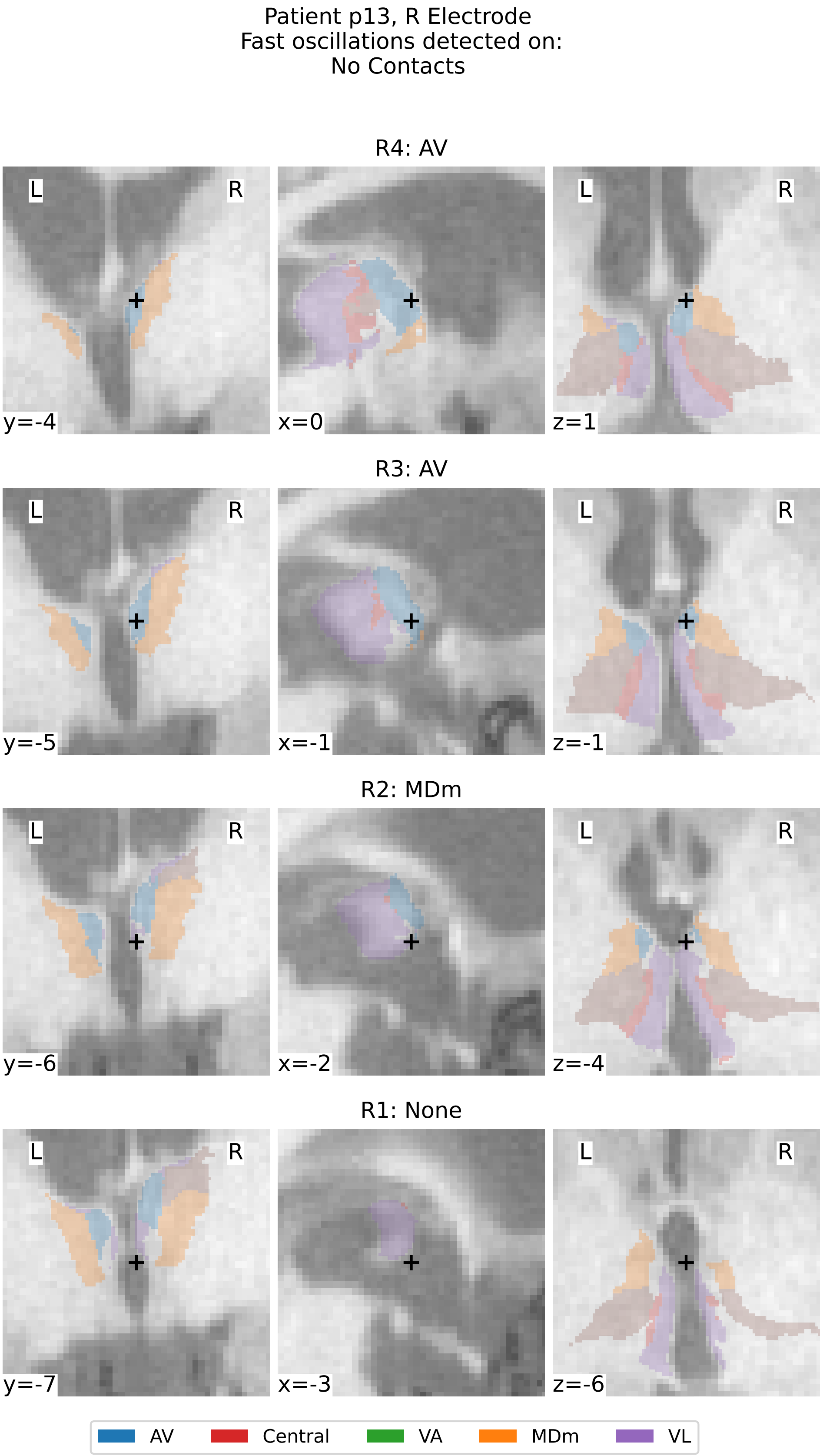


Supplementary Figure 38: **Thalamic Segmentation and Localization of contacts for patient p13, right electrode.** See caption of Supplementary Figure 12.

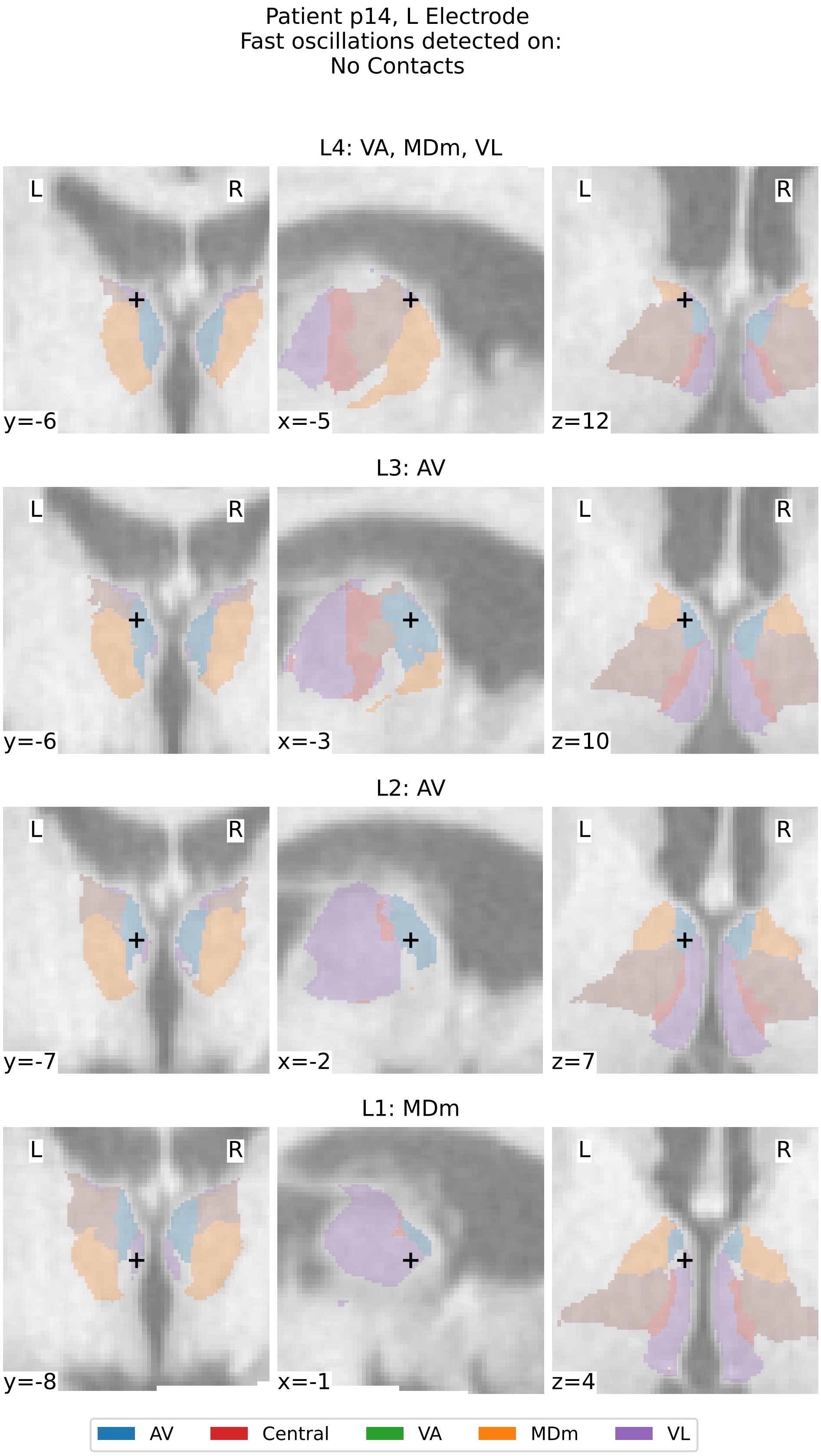


Supplementary Figure 39: **Thalamic Segmentation and Localization of contacts for patient p14, left electrode.** See caption of Supplementary Figure 12.

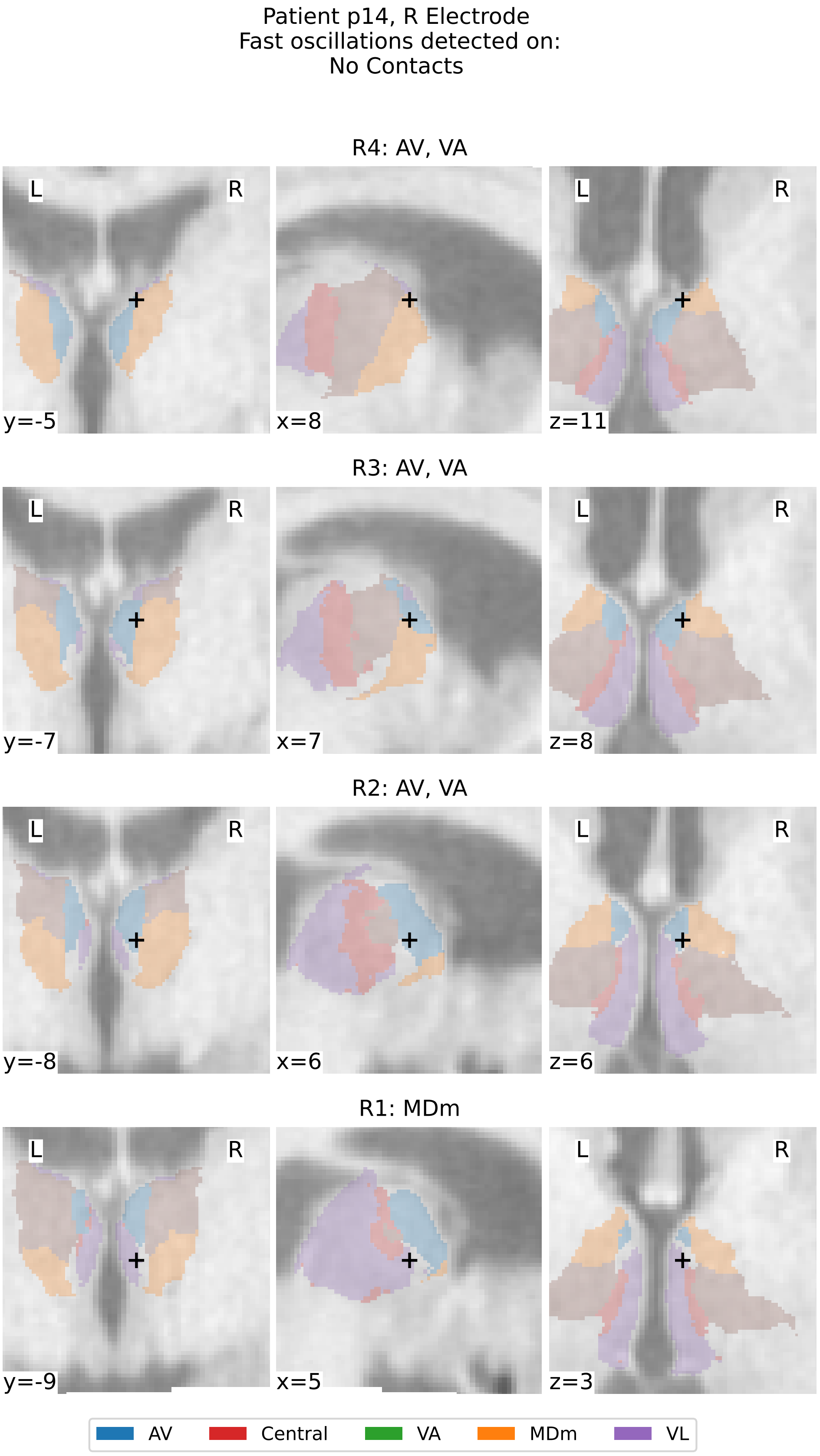


Supplementary Figure 40: **Thalamic Segmentation and Localization of contacts for patient p14, right electrode.** See caption of Supplementary Figure 12.

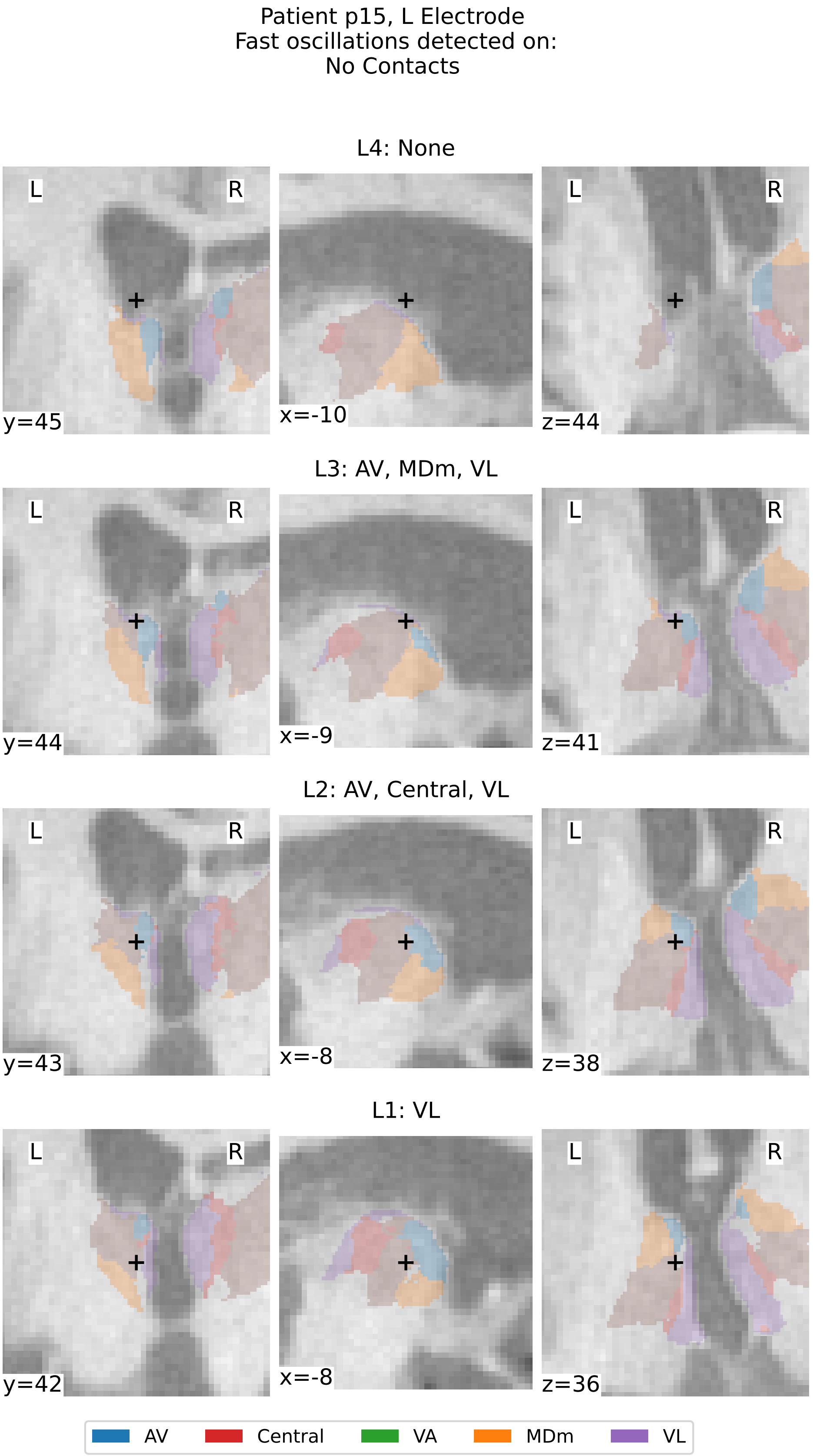


Supplementary Figure 41: **Thalamic Segmentation and Localization of contacts for patient p15, left electrode.** See caption of Supplementary Figure 12.

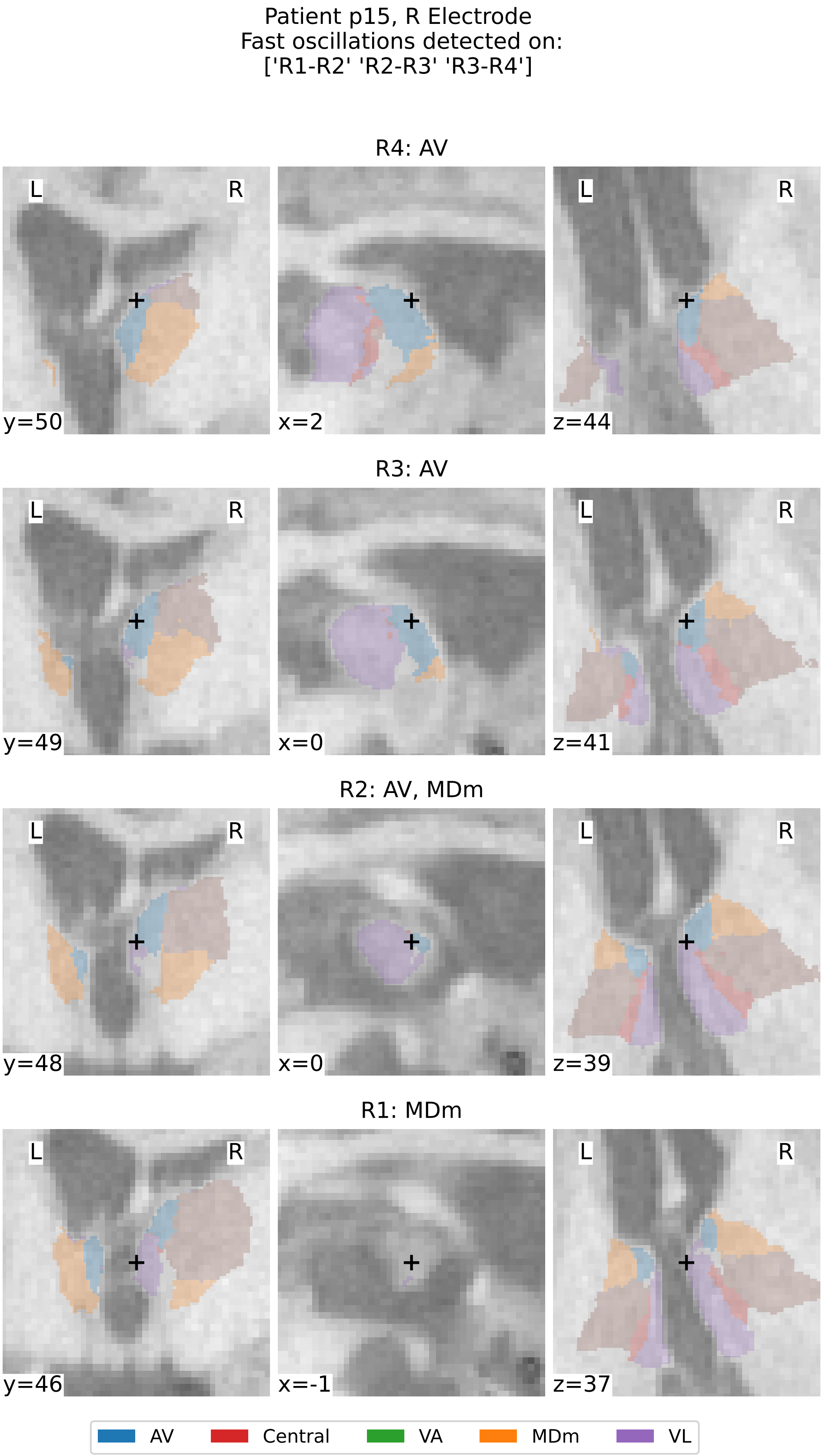


Supplementary Figure 42: **Thalamic Segmentation and Localization of contacts for patient p15, right electrode.** See caption of Supplementary Figure 12.

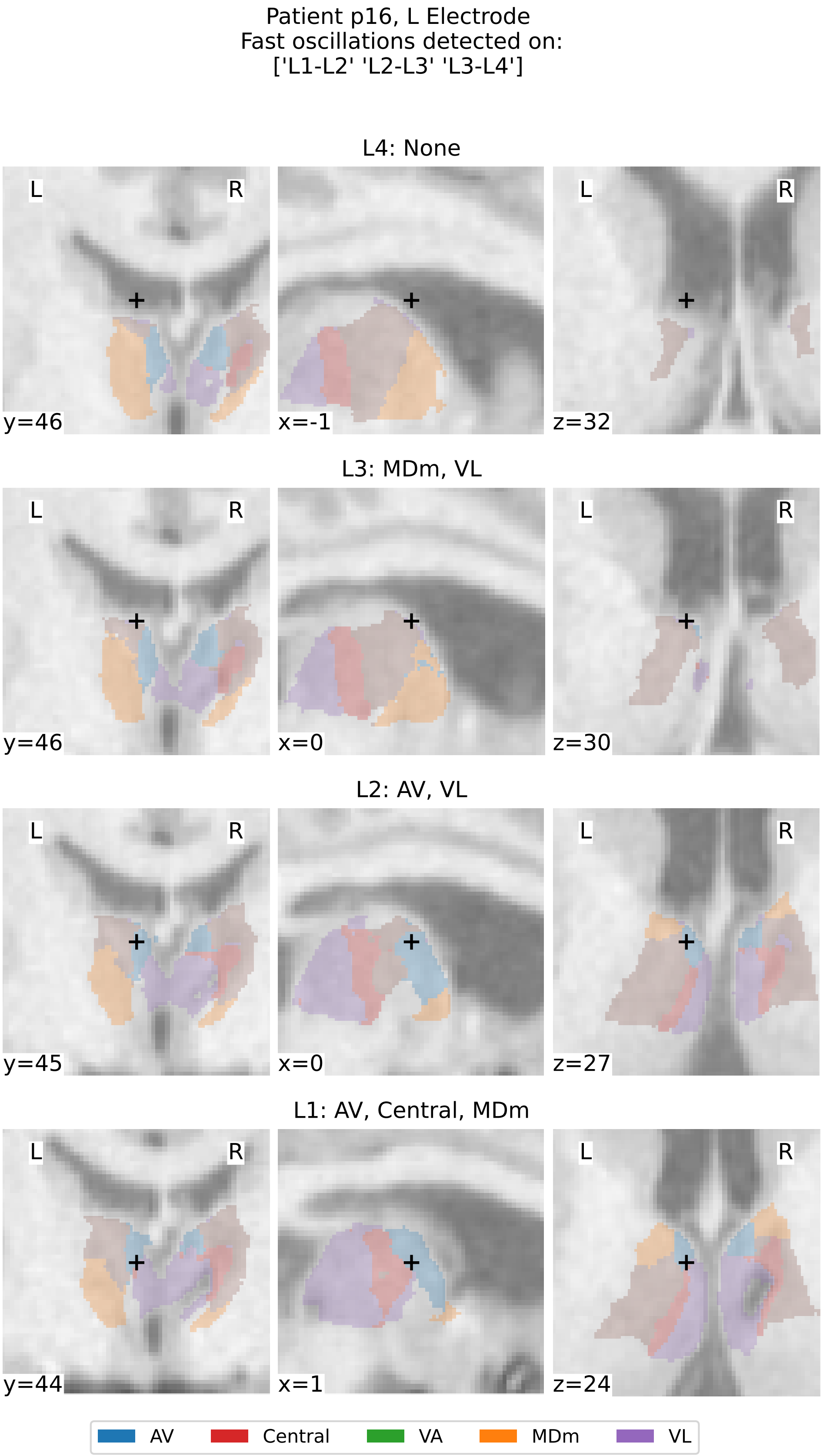


Supplementary Figure 43: **Thalamic Segmentation and Localization of contacts for patient p16, left electrode.** See caption of Supplementary Figure 12.

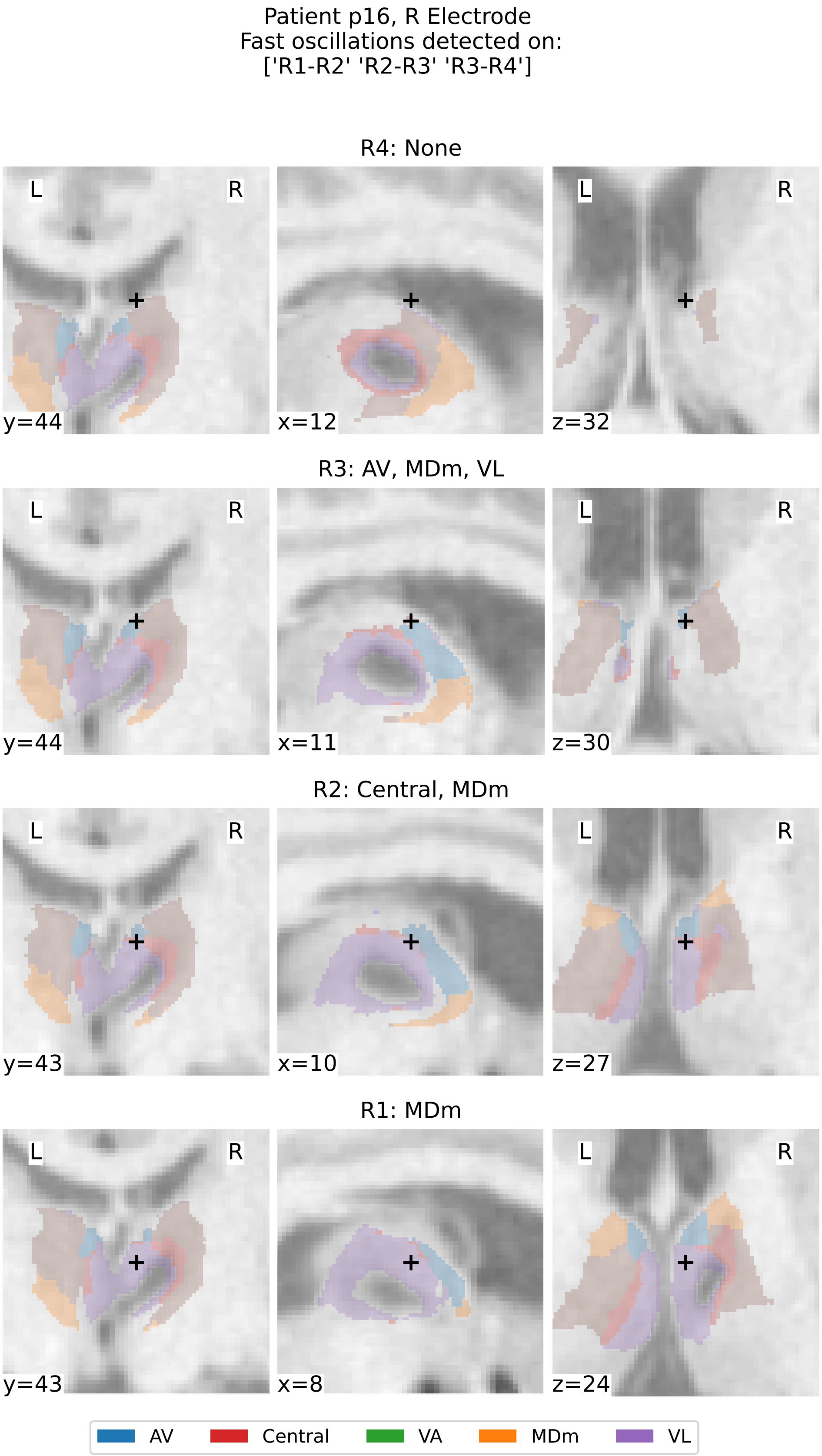


Supplementary Figure 44: **Thalamic Segmentation and Localization of contacts for patient p16, right electrode.** See caption of Supplementary Figure 12.

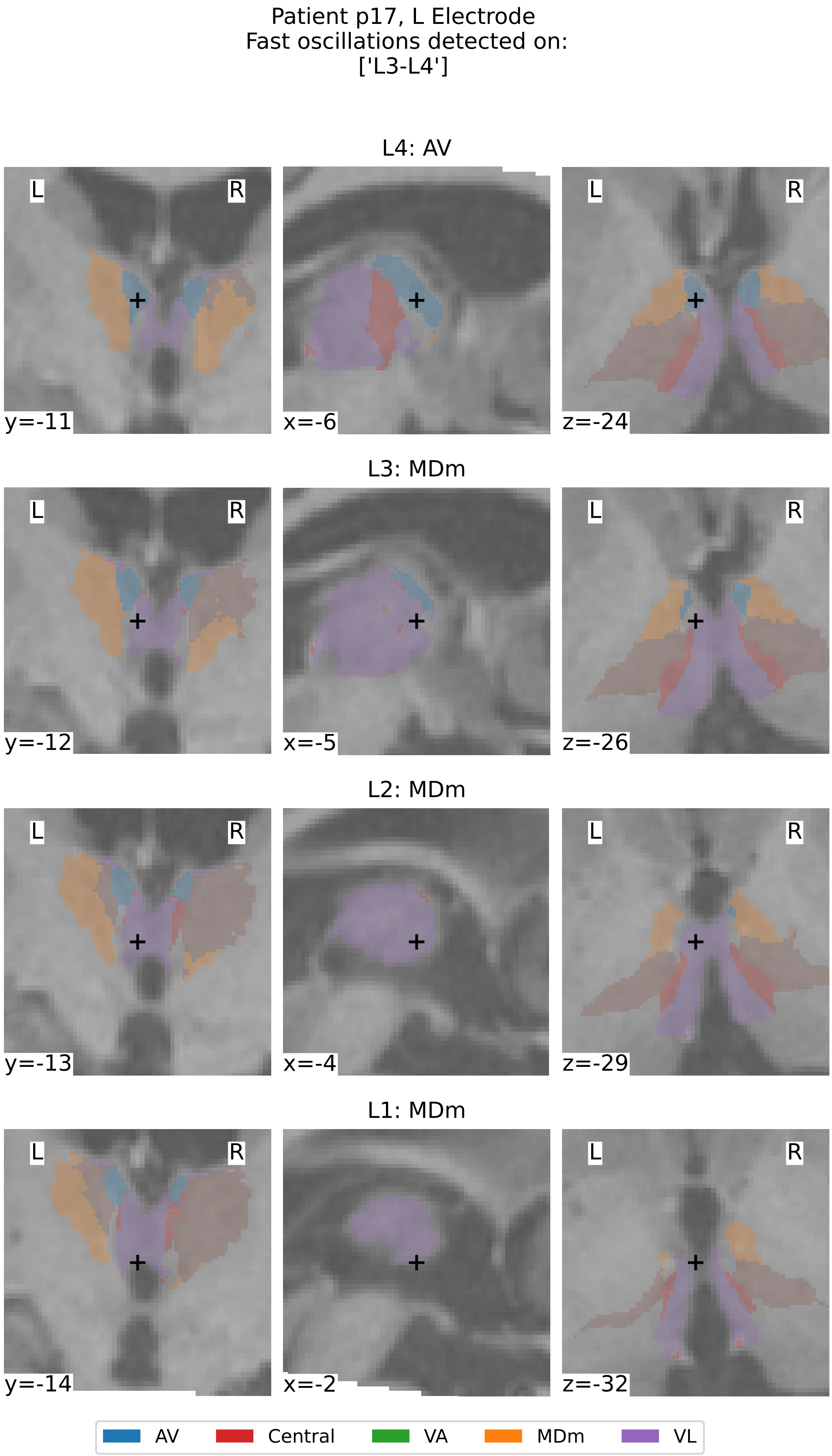


Supplementary Figure 45: **Thalamic Segmentation and Localization of contacts for patient p17, left electrode.** See caption of Supplementary Figure 12.

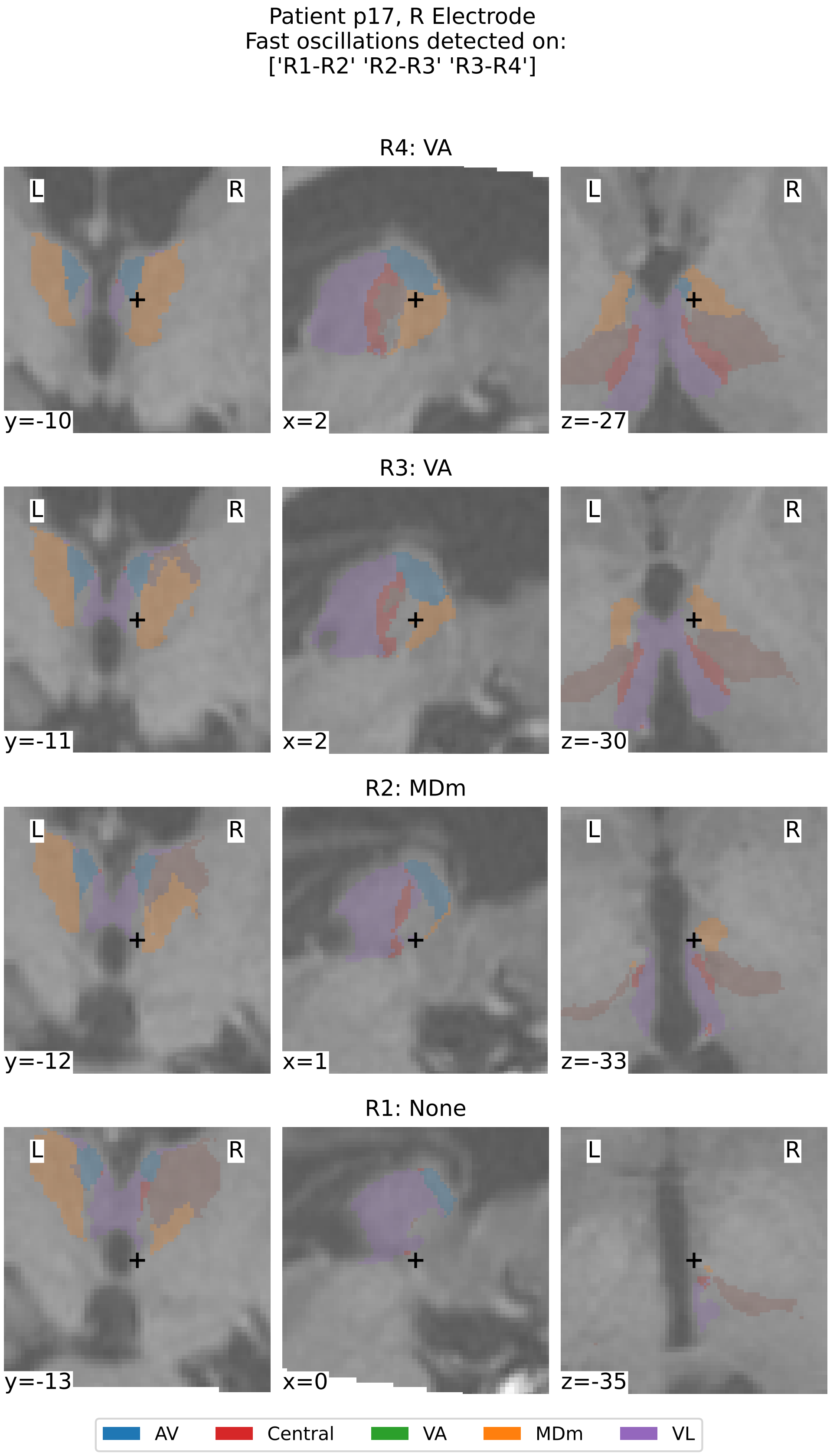


Supplementary Figure 46: **Thalamic Segmentation and Localization of contacts for patient p17, right electrode.** See caption of Supplementary Figure 12.
